# Promising Antimalarials targeting Apicoplast DNA Polymerase from *Plasmodium falciparum*

**DOI:** 10.1101/2022.03.07.483284

**Authors:** Pratik R. Chheda, Nicholas Nieto, Supreet Kaur, John M. Beck, Josh R. Beck, Richard Honzatko, Robert J. Kerns, Scott W. Nelson

## Abstract

Malaria is caused by the parasite *Plasmodium falciparum*, which contains an essential non-photosynthetic plastid called the apicoplast. A single DNA polymerase, apPOL, is targeted to the apicoplast, where it replicates and repairs the genome. apPOL has no direct orthologs in mammals and is considered a promising drug target for the treatment and/or prevention of malaria. We previously reported screening the Malaria Box to identify MMV666123 as an inhibitor of apPOL. Herein we extend our studies and report structure-activity relationships for MMV666123 and identify key structural motifs necessary for inhibition. Although attempts to crystallize apPOL with the inhibitor were not fruitful, kinetic analysis and crystal structure determinations of WT and mutant apo-enzymes, facilitated model building and provided insights into the putative inhibitor binding site. Our results validate apPOL as an antimalarial target and provide an avenue for the design of high potency, specific inhibitors of apPOL and other A-family DNA polymerases.

---

Malaria is a mosquito borne infectious disease caused by the parasites of the genus *Plasmodium*, with *P. falciparum* and *P. vivax* being the most common and deadly agents, causing more than 200 million cases of infection and 400,000 deaths annually.^1-3^ Despite the recent advances in medicine and insect control programs, malaria continues to remain endemic in several countries with nearly half of the world’s populations at risk of developing infection.^2^ Upon introduction into the vertebrate host during the bite of an infected mosquito, the parasites initially migrate to the liver and undergo a developmental transition before being released into the circulation where they infect red blood cells to produce all of the pathologies of the disease.^4^ If left untreated, the growing parasite load can lead to intravascular hemolysis and multiple organ failure ultimately leading to death. Currently, chemotherapy is the only means to manage the infection with Artemisinin-combination therapies (ACTs) being the most efficacious treatment available.^2,5^ However, resistance to Artemisinin and other co-administered drugs is established in Southeast Asia and is emerging in Africa where most severe disease and deaths occur, jeopardizing global malaria control programs.^6-9^ This accentuates the need for development of new drugs that target novel aspects of the parasite’s biology and could be used either alone or in combination for the management of Malaria.

*Plasmodium spp*. belong to the phylum Apicomplexa, which also includes many related pathogenic organisms (e.g., *Toxoplasma* and *Cyclospora*), the majority of which have an unusual organelle called the apicoplast.^10^ The apicoplast is evolutionarily related to the chloroplast, arising putatively through a secondary endosymbiotic event with red algae.^11^ The apicoplast is the site for several biochemical processes including fatty acid biosynthesis, heme synthesis, Fe-S cluster maturation and isoprenoid biosynthesis, making it indispensable for the parasite.^12-13^ In previous studies, *Plasmodium spp*. treated with antibiotics targeting the apicoplast prokaryotic-like ribosome, like azithromycin^14^, clindamycin^15^ and tetracyclines^16^ lead to parasites that grow and divide but produce progeny of schizonts that inherited non-functional apicoplasts and are unable to differentiate into merozoites that are necessary to invade new erythrocytes, causing a “Delayed-Death” phenomenon.^17-19^ Thus, drugs targeting the apicoplast can be used as monotherapy for prophylaxis or in combination with other rapid acting antimalarials for management of active malarial infection.

The apicoplast has its own 35 kb genome and hosts nearly 600 proteins, most of which are encoded by genes belonging to the nuclear genome of the parasite.^20^ Of the several apicoplast replication proteins, the apicoplast DNA polymerase (apPOL) is essential to the parasite and is clearly of prokaryotic origin with its nearest homolog outside of Apicomplexa being the replicative polymerase from the cyanobacteria *Cyanothece* sp. PCC 8802 (35% protein sequence identity).^21-23^ The most similar human polymerases are the lesion bypass polymerases theta and nu having 23% and 22% sequence identity, respectively, providing the basis for selectivity. On the other hand, apPOL from *P. falciparum* and *P. vivax* have 84% homology, suggesting that drugs targeted against the *Pf-*apPOL would be effective in treating *P. vivax* infections as well, making apPOL an attractive target for development of novel antimalarials.^24^ To identify novel inhibitors of apPOL, we had screened the Open Access Malaria Box, using our high-throughput assay, and reported identification of MMV666123 (**5a**) as a selective, micro molar (IC_50_∼3 μM) inhibitor of apPOL (Fig. 1).^25-26^ Herein we report the rational design, synthesis and biological evaluation of a series of **5a** derivatives leading to extensive SAR and understanding the enzymatic mechanism of inhibition of apPOL by these inhibitors. We also report insights into the putative inhibitor binding site with the identification of urea analogs of **5a** as the potential candidates for further lead optimization.

**Figure 1.**
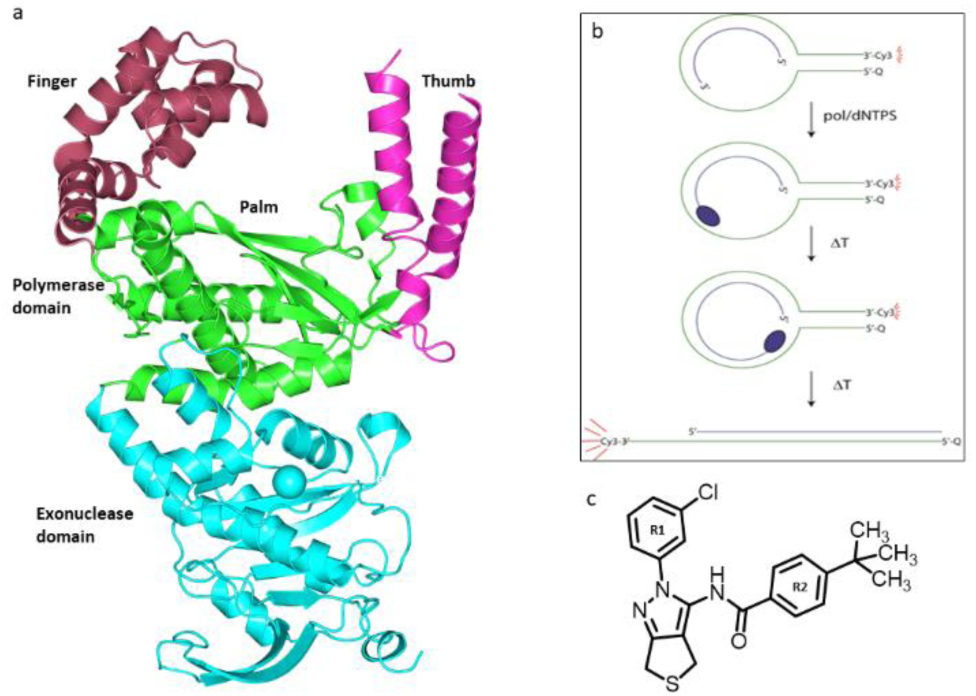
a) apPOL structure, b) Schematic showing DNA substrate used in molecular beacon assay to quantitate apPOL activity, c) structure of compound 5a

## Chemical Synthesis

Structural diversity was introduced to peripheral substituents of **5a** by synthesizing variably substituted 2-phenyl-4,6-dihydro-2H-thieno[3,4-c]pyrazol-3-amine cores following modification of a 2-step protocol reported by Baraldi, PR. et al.^27^ Commercially available methyl 2-mercaptoacetate (**1**) was stirred with acrylonitrile (**2**) in presence of sodium methoxide in methanol to give tetrahydrothiophene **3**, which on reaction with variably substituted phenyl hydrazines in refluxing ethanol gave 3-amino pyrazoles **4a-f** in moderate to good yields (**Scheme 1**). Acylation of **4a-f** with commercially available 4-tertbutyl benzoyl chloride under basic conditions furnished the amide linked analogs **5a-f** in moderate yields (**Scheme 1**). Alternatively, treating intermediates **4a-f** with either 4-tertbutyl benzene sulfonyl chloride or 4-tertbutyl phenyl isocyanate under basic conditions furnished the desired sulfonamide (**6a-f**) and urea linked (**7a-f**) analogs.(**Scheme 1**).

**Scheme 1:**
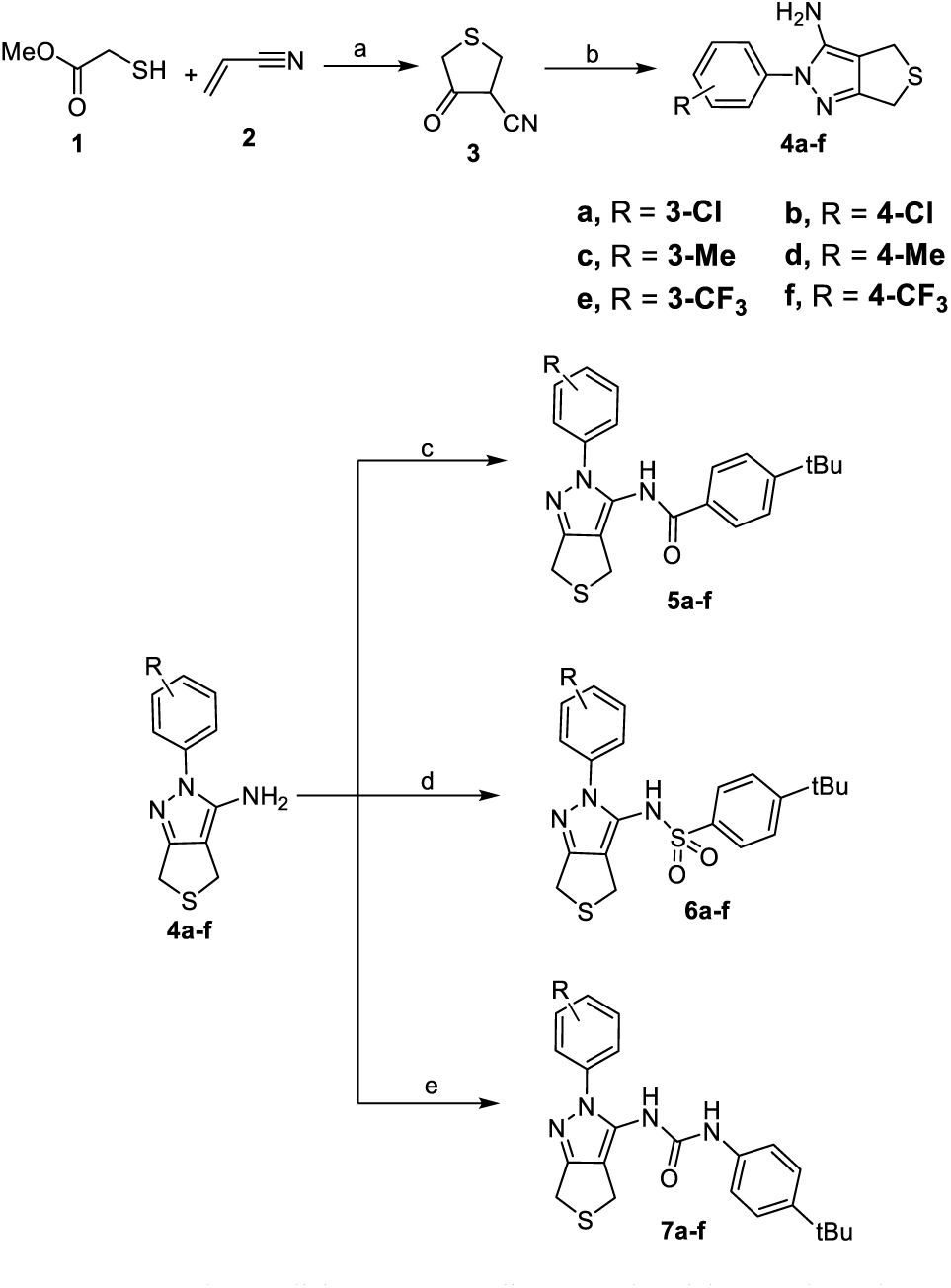
Synthesis of 5a-f, 6a-f, 7a-f ^a, *b*^. ^*a*^ Reagents and Conditions: (a) Sodium methoxide, methanol, 0 °C-RT, 42%; (b) substituted phenyl hydrazine, ethanol, reflux, 54-76%; (c) 4-tertbutyl benzoyl chloride, di-isopropyl ethyl amine, anhydrous dichloromethane, RT, 58-69%; (d) 4-tertbutyl benzene sulfonyl chloride, sodium hydride, anhydrous dimethylformamide, RT, 52-59%; (e) 4-tertbutyl phenyl isocyanate, sodium hydride, anhydrous dimethylformamide, RT, 42-73%. ^*b*^ For complete structures, see Table 1.

Upon extended storage and several freeze-thaw cycles of the stock solution of the initial hit, we observed slow oxidation of the tetrahydrothiophene thioether to sulfoxide. To evaluate if the thioether is key for activity, we synthesized sulfoxide derivative **8** by treating **5a** with hydrogen peroxide and acetic anhydride in the presence of silica gel in dichloromethane (**Scheme 2**).

**Scheme 2:**
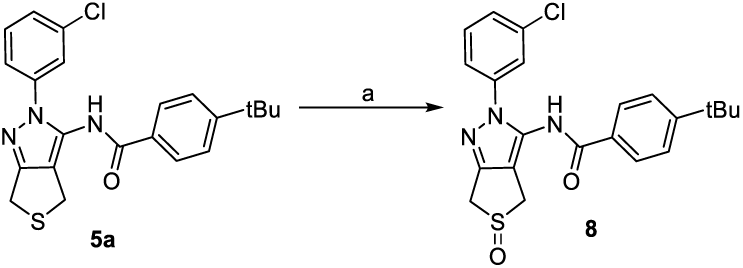
Synthesis of 8. Reagents and Conditions: (a) Hydrogen peroxide, silica, acetic anhydride, dichloromethane, r.t., quantitative; (b) meta-chloroperoxybenzoic acid, dichloromethane, r.t., quantitative.

Analogs having a substituted 2-phenyl-4,6-dihydro-2H-furo[3,4-c]pyrazol-3-amine core were synthesized from reaction between commercially available methyl 2-hydroxyacetate (**9**) with acrylonitrile (**2**) under basic conditions to give tetrahydrofuran **11** in 48% yield, which on refluxing with variably substituted phenyl hydrazines in ethanol afforded amino pyrazoles **12a-c** in moderate yields (**Scheme 3**). Intermediates **12a-c** were then reacted with variably substituted phenyl isocyanates to furnish urea linked analogs **13a-c**.

**Scheme 3:**
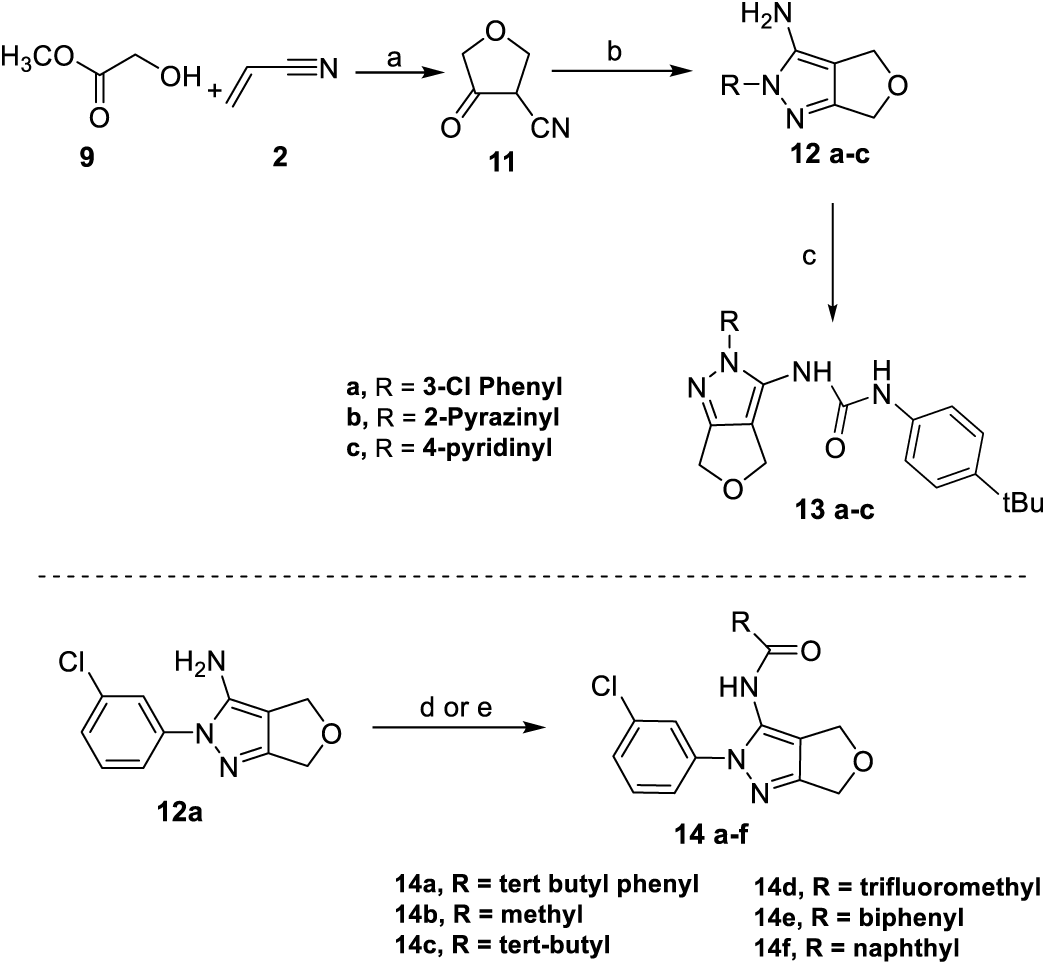
Synthesis of tetrahydrofuran analogs 13a-c, 14a-f ^a, b^. ^*a*^ Reagents and Conditions: (a) Sodium hydride, anhydrous tetrahydrofuran, 0 °C-RT, 48%; (b)substituted phenyl hydrazine, ethanol, reflux, 52-68%; (c) 4-tertbutyl benzoyl chloride, di-isopropyl ethyl amine, anhydrous dichloromethane, RT, 58-69%; (d) substituted acid chloride, di-isopropyl ethyl amine, anhydrous dichloromethane, RT; (e) N-methyl imidazole, tetramethylethylenediamine, potassium carbonate, 2-naphthoyl chloride or [1,1’-biphenyl]-4-carbonyl chloride, acetonitrile 0 °C. ^*b*^ For complete structures, see Table 2.

It is notable that yields for forming the urea linkage were low because of formation of bis-urea side products under the reaction conditions. Lastly, variably substituted target compounds **14a-d** were synthesized following amide coupling procedures as describedfor compound **5a**. Alternatively, the target compounds substituted with naphthyl (**14f**) and biphenyl (**14e**) groups were synthesized by employing a mixture of three bases (N-methyl imidazole, tetramethylethylenediamine and potassium carbonate) in acetonitrile to furnish the desired products in moderate yields (**Scheme 3**).

All synthesized analogs, summarized in **Table 1**, were first tested *in vitro* for their potency to inhibit purified *Pf*. apPOL^exo−^ polymerase activity using our Molecular Beacon Assay.^26^ In short, a stem-loop oligonucleotide, with a Cy3 dye at 5’ and Black Hole Quencher at the 3’ end, and a primer annealed to the loop region, was used as a substrate for the DNA polymerase assays. Inhibitor potency was measured by their ability to prevent strand displacement synthesis by apPOL^exo−^, which opens the DNA duplex separating the Cy3 fluorophore from the quencher resulting in an increased fluorescence. Overall, *IC*_*50*_ values for the diverse analogs varied by approximately 130-fold with most derivatives displaying poorer inhibition compared to **5a** (**Table 1**).

**Table 1:** Summary of Activity.

| Compound | Scaffold | X | R<sup>1</sup> | apPOL<br>IC<sub>50</sub><br>(μM) | *P.falciparum*<br>EC<sub>50</sub> (μM) | Exonuclease<br>specific<br>activity | Exonuclease<br>EC<sub>50</sub> (μM) |
| --- | --- | --- | --- | --- | --- | --- | --- |
| - | - | - | - | - | - | 2.2 ±0.09 | - |
| **5a** | 1 | S | 3-Cl | 2.9 ±0.3 | 4.0 ±1.1 | 1.6 ±0.1 | - |
| **5b** | 1 | S | 4-Cl | 11 ±3 | 4.2 ±0.6 | 2.4 ±0.2 | - |
| **5c** | 1 | S | 3-Me | 4.8 ±1.2 | 5.9 ±0.07 | 2.3 ±0.1 | - |
| **5d** | 1 | S | 4-Me | 6.5 ±1.2 | >20 | 1.5 ±0.05 | - |
| **5e** | 1 | S | 3-CF<sub>3</sub> | 6.1 ±1.4 | 2.0 ±0.03 | 2.4 ±0.1 | - |
| **5f** | 1 | S | 4- CF<sub>3</sub> | 5.9 ±1.5 | 4.8 ±0.8 | 2.0 ±0.1 | - |
| **6a** | 2 | S | 3-Cl | 8.3 ±3.4 | >20 | 0.4 ±0.05 | 40 ±5.7 |
| **6b** | 2 | S | 4-Cl | 63 ±12 | >20 | 0.06 ±0.03 | 21 ±2.5 |
| **6c** | 2 | S | 3-Me | 55 ±8 | >20 | 2.3 ±0.1 | - |
| **6d** | 2 | S | 4-Me | 25 ±6 | >20 | 2.2 ±0.05 | - |
| **6e** | 2 | S | 3-CF<sub>3</sub> | 38 ±3 | >20 | 0.06 ±0.001 | 41 ±6.7 |
| **6f** | 2 | S | 4- CF<sub>3</sub> | 36 ±12 | >20 | 0.01 ±0.004 | 16 ±0.9 |
| **7a** | 3 | S | 3-Cl | 3.9 ±0.5 | 1.3 ±0.04 | 2.6 ±0.03 | - |
| **7b** | 3 | S | 4-Cl | 2.4 ±0.4 | 1.0 ±0.09 | 2.1 ±0.2 | - |
| **7c** | 3 | S | 3-Me | 5.6 ±0.6 | 3.1 ±0.3 | 1.2 ±0.2 | - |
| **7d** | 3 | S | 4-Me | 5.3 ±0.5 | 4.5 ±0.8 | 1.8 ±0.2 | - |
| **7e** | 3 | S | 3-CF<sub>3</sub> | 3.0 ±0.4 | 4.3 ±0.1 | 1.4 ±0.3 | - |
| **7f** | 3 | S | 4- CF<sub>3</sub> | 6.4 ±0.6 | 1.2 ±0.7 | 2.1 ±0.2 | - |
| **8** | 1 | sulfoxide | 3-Cl | 65 ±14 | 4.8 ±0.7 | 1.9 ±0.2 | - |
| **13a** | 3 | O | 3-chlorophenyl | 8.3 ±0.8 | 0.74 ±0.01 | 2.4 ±0.3 | - |
| **13b** | 3 | O | 2-Pyrazinyl | 20 ±7.8 | 2.5 ±0.8 | 1.7 ±0.2 | - |
| **13c** | 3 | O | 4-Pyridinyl | 35 ±7.9 | 3.3 ±0.08 | 1.3 ±0.1 | - |
| **14a** | 4 | O | 4-*tert*butyl<br>phenyl | 8.7 ±4.5 | 4.6 ±0.2 | 1.2 ±0.1 | - |
| **14b** | 4 | O | -CH<sub>3</sub> | 317 ±69 | >20 | 2.2 ±0.1 | - |
| **15c** | 4 | O | -C(CH<sub>3</sub>)<sub>3</sub> | 253 ±85 | >20 | 2.6 ±0.3 | - |
| **14d** | 4 | O | -CF<sub>3</sub> | 262 ±59 | >20 | 2.3 ±0.1 | - |
| **14e** | 4 | O | 1,1'-biphenyl | 7.5 ±2.6 | >20 | 1.2 ±0.1 | - |
| **14f** | 4 | O | 2-naphthyl | 21 ±4.5 | 8.0 ±0.002 | 1.8 ±0.2 | - |

The sulfonamide analogs were in general 2-10-fold less active than **5a**, whereas the urea linked analogs retained potency similar to **5a**. Only **6a** being the exception for sulfonamide analogs, the 3-Cl R-substituent of **6a** retained inhibition constants similar to amide-and urea-linked derivatives. Overall, the nature and position of the substituent on the phenyl ring of scaffolds 1-4 did not affect inhibitory activity of the analogs, however, introduction of heteroatoms into the phenyl ring (**13b-c**) was detrimental to activity. This indicates the phenyl ring likely occupies a hydrophobic pocket. The sulfoxide analog **8** was 10-fold less active than **5a**, whereas the urea-linked tetrahydrofuran analog **13a** was only 2-fold less active, indicating that sulfur in the saturated ring is a possible metabolic liability and this liability can be alleviated by replacing oxygen for sulfur without destroying the apPOL inhibitory activity. Based on this data, the second series of analogs were synthesized (**Scheme 3**), all of which have a tetrahydrofuran core and introduce structural variation in the tertbutyl phenyl region of **5a**. Testing a small set of analogs with different size substituents (**14a-f**) quickly indicated that a bulky tertbutyl phenyl group is necessary to retain activity; analogs containing either a small methyl, trifluoromethyl, or even a t-butyl substituent (**14a-b**) were >100-fold less active than **5a**. Overall, three urea linked compounds **7a, 7b**, and **7e** retained the inhibitory properties of the parent compound (**Figure 2a**) and were more soluble than **5a** based on a visual analysis.

**Figure 2.**
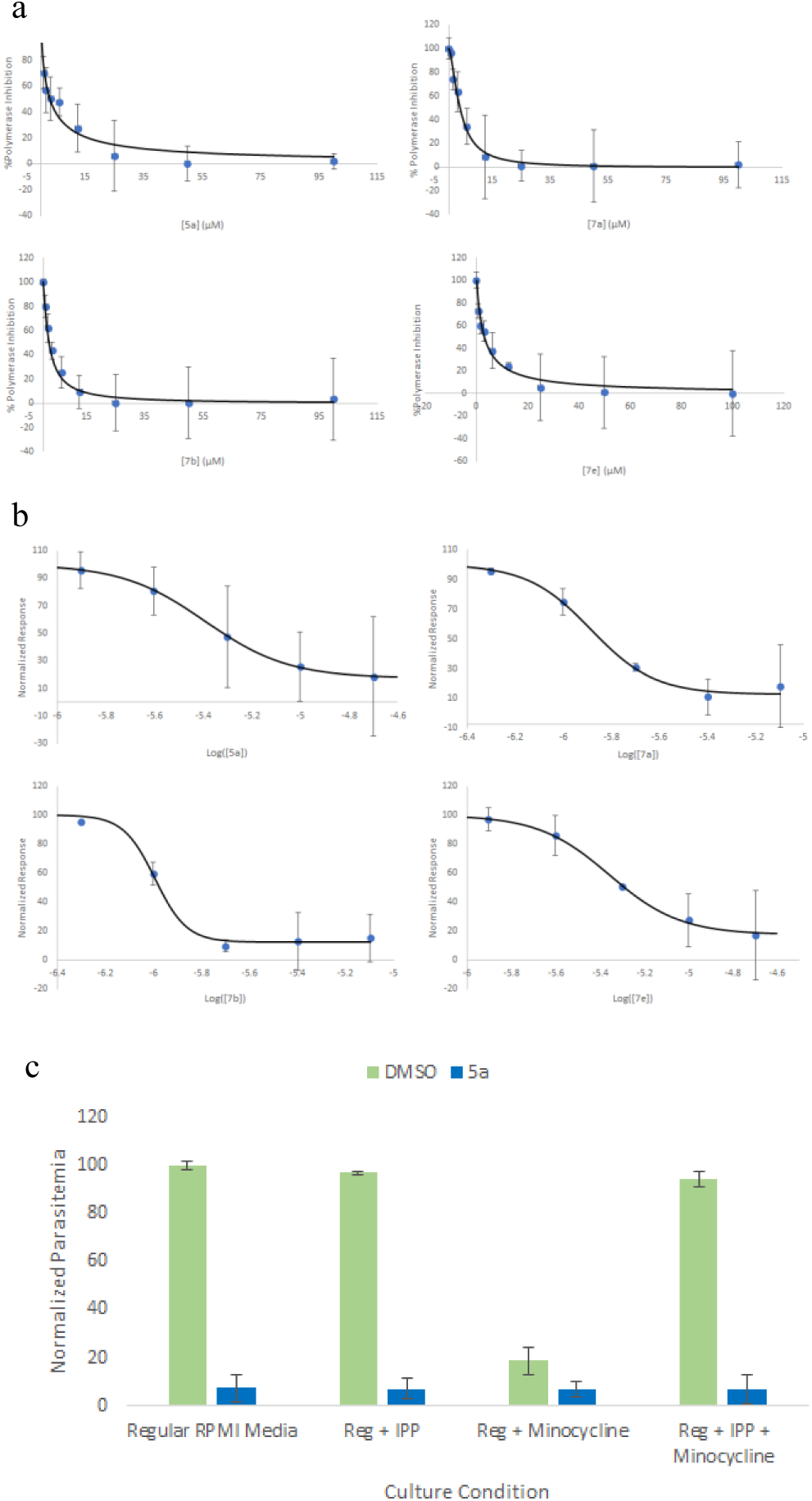
IC_50_, EC_50_, and IPP Rescue data. A) IC_50_, curves for **5a, 7a, 7b**, and **7e**. B.) EC_50_ data for **5a, 7a, 7b**, and **7e** C) IPP rescue data for **5a**

The apPOL inhibitors were simultaneously evaluated for their ability to kill *P. falciparum* parasites using NF54^attB^ parasites cultured in red blood cells in presence of increasing inhibitor concentrations followed by measuring parasitemia at 72-hour time points using flow cytometry (**For additional details – see SI**).^28^ The majority of urea and amide-linked compounds (**5a-f, 7a-f**) gave EC_50_ values within single digit µM range that corresponded to their in vitro IC_50_ values (**Table 1**). Similar to apPOL inhibitory activity, the potency of compounds decreased when linker was changed to sulfonamide (**6a-f**) or when the bulky t-butylphenyl group was replaced with smaller non-bulkier moieties (**14b-e**). Gratifyingly, the same urea linked compounds **7a, 7b** and **7e** which had best overall apPOL inhibition were also among those with the lowest EC_50_ in the parasite killing assay (**Figure2b)**. Additionally, the tetrahydrofuran analog **13a**, which had apPOL inhibition similar to **5a**, but lacked its metabolic liability and had improved solubility was the most potent inhibitor in the parasite killing assay with a nanomolar EC_50_ of 740 nM.

Isoprenoids (IPP) are essential for the replication and survival of *Plasmodium* parasites during the blood-stage of infection and the apicoplast is the only site for isoprenoid precursor biosynthesis. IPP supplementation in parasite cultures treated with apicoplast-targeting antimalarials rescues parasite death following loss of the organelle, this is an essential tool to validate candidate compounds that specifically target and disrupt apicoplast function.^13^ To evaluate this, IPP rescue assays were performed initially on the parent compound by growing parasites in either 20 μM inhibitor (highest concentration used for EC_50_ analysis) or DMSO under four different conditions and measuring parasitemia by flow cytometry after 96 hours (**For additional details – see SI**).^28^ Surprisingly, **5a** retained its lethality in presence of IPP and did not display IPP rescue compared to a positive control using minocycline, suggesting a secondary target for these inhibitors outside of the apicoplast (**Figure 2c**). The most likely secondary target protein could be Polγ, which is another A-family DNA polymerase found in the mitochondria of plasmodial parasites. Currents attempts to clone the mitochondrial polymerase to characterize potential inhibition by these compounds have thus far not been fruitful and we continue b our efforts towards this. Due to a lack of significant improvement in inhibitor potency for the derivative library, subsequent IPP rescue was not performed on any of the parent compound derivatives.

apPOL is a A-family DNA polymerase and contains a 3’-5’ exonuclease domain, therefore, we wanted to evaluate the compound libraries selectivity to inhibit the polymerase activity over exonuclease activity. 3’-5’ Exonuclease activity of apPOL was quantified using a blunt-ended double stranded oligonucleotide containing a 2’aminopurine at position one on 3’end as the substrate for apPOL.^29^ Incubating with DNA substrate, WTapPOL, containing intact exonuclease domain, binds the 3’end of DNA substrate and proofreads in 3’-5’ direction releasing the 2-aminopurine at position 1 resulting in an increase fluorescence over time. Single substrate DNA concentrations were used to determine the specific activity of apPOL exonuclease domain in the presence and absence of 100 µM inhibitors to compare their strengths (**Table 1**). Compounds displaying nearly 100% inhibition were subsequently analyzed for IC_50_ determination (**For additional details – see SI**). Majority of compounds, including all urea-and amide-linked compounds, were selective and did not inhibit exonuclease enzyme activity more than 50% at 100µM. Interestingly, only the sulfonamide linked compounds with electron withdrawing substituents (**6a**,**b**,**e**,**f**) showed >90% exonuclease inhibition at 100µM (with para substituents showing increased affinity over the meta analogs). These results suggest subtle changes within the inhibitor scaffold, in this case the compound linker, can result in redirecting compound binding affinity to a completely different region of the enzyme which is remarkable.

Having identified analogs with minor improvements in activity against apPOL and *P. falciparum* parasites compared to **5a**, we next wanted to gain structural information on inhibitor binding site(s) to guide the next round of inhibitor design. To enable this, we decided to perform enzyme kinetics and crystallography studies in parallel to gain further insights into the mechanism of inhibition. Kinetic mechanism of apPOL polymerase inhibition was determined for **5a** and three urea-analogs **7a, 7b** and **7e** using the molecular beacon assay and Dynafit software.^30^ Data was collected at various dNTP and inhibitor concentrations and fitting to several kinetic models was analyzed using the Dynafit model discrimination analysis feature.^28^

A combination of visual analysis and Model discrimination analysis using the Akaike information criterion^31^ (AIC) and the Bayesian information criterion^32^ (BIC) indicated that the data fit best to a non-competitive model (**Figure 3a**). However, we consistently observed that the data points for the series at the highest inhibitor concentration were lower than predicted by a simple non-competitive model, suggesting a more complex model may be necessary (**Figure 3b&c**). To better fit the data, we evaluated other models involving binding of two inhibitor molecules to the polymerase including non-interacting and cooperative models.

**Figure 3.**
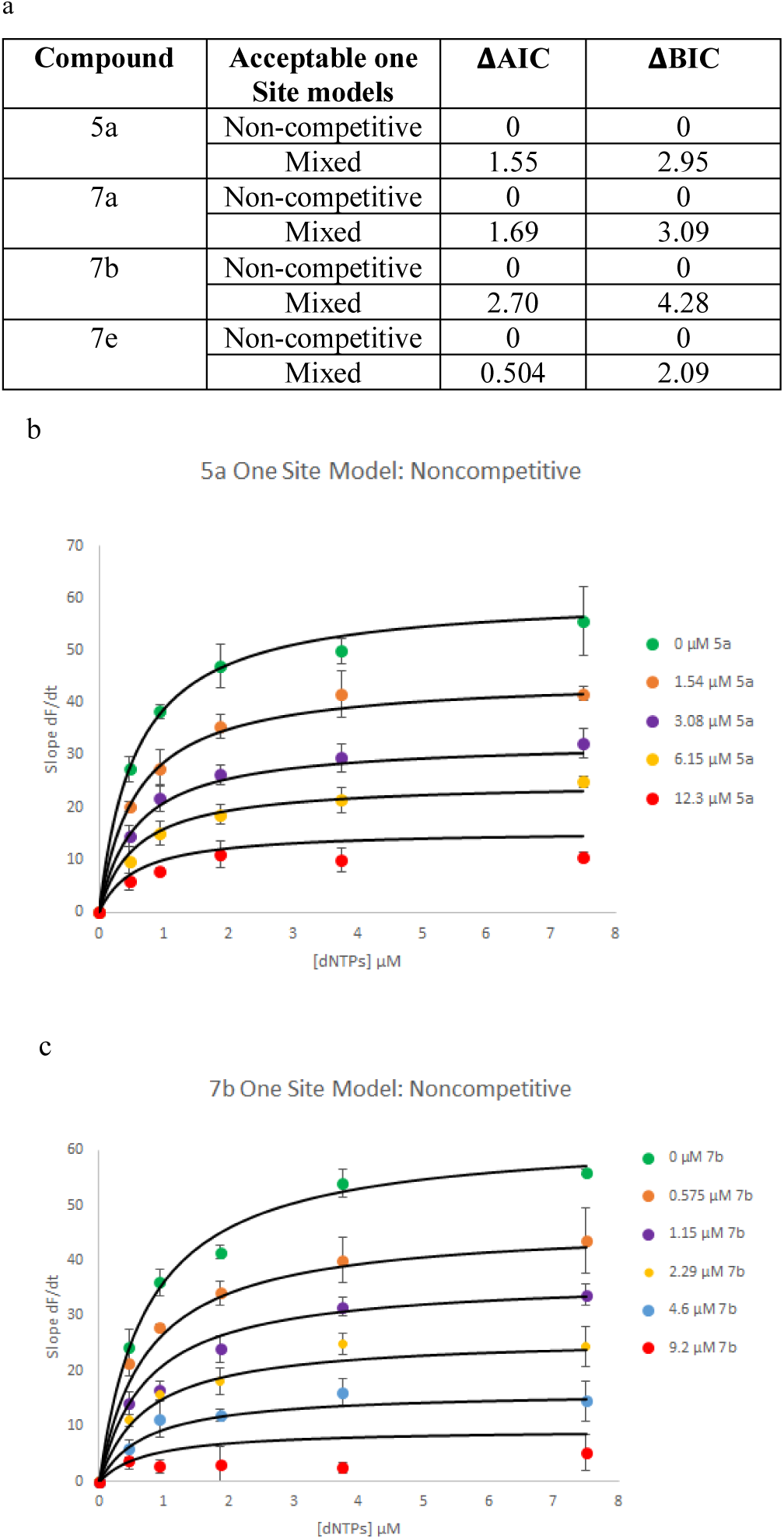
Dynafit results for fitting mechanism data to single site models of inhibition (competitive, noncompetitive, uncompetitive, mixed). a.) Table summarizing results from dynafit including acceptable models and statistical values AIC and BIC that were used to distinguish which model was most consistent with the data. b.) 5a mechanism data graph showing experimental data points (colored circles) plotted with fitted data from dynafit (black smooth lines) for a noncompetitive single site model. c.) 7b mechanism data graph showing fitted data is weakly fitting to the experimental data at highest concentration of 7b indicating a more complicated mechanism may be involved.

For compound **5a**, the best-fitting mechanism was a model with two non-identical and independent sites, both of which are non-competitive against dNTPs with K_I_ for high-affinity binding site being 6.1 μM and the K_II_ of low-affinity binding site being 15.9 μM (**Figure 4a**). Similar analysis on urea analogs indicated that **7a** and **7b** fit best to the same model as **5a** with K_I_ values of 2.7 and 2.0 μM for **7a** and **7b**, respectively, and K_II_ values of 12.2 and 15.0 μM, respectively. Interestingly, the data for inhibition by **7e** fit best to an independent two-site model, similar to the other inhibitors, but in this case, only the high affinity site (K_I_) was non-competitive against dNTPs, whereas the lower affinity site (K_II_) was competitive. Overall, the data from SAR studies suggest modifications made so far to scaffold lead to only 2 to 5-fold improvement in the activity compared to **5a** and the mechanistic studies indicate presence of a second binding site, further complicating the receptor binding. Further optimizing inhibitor potency rationally would require structural characterization of enzyme-inhibitor binding to understand key interacting amino acids in the pocket and so we tried to resolve the crystal structure of the inhibitor bound apPOL.

**Figure 4.**
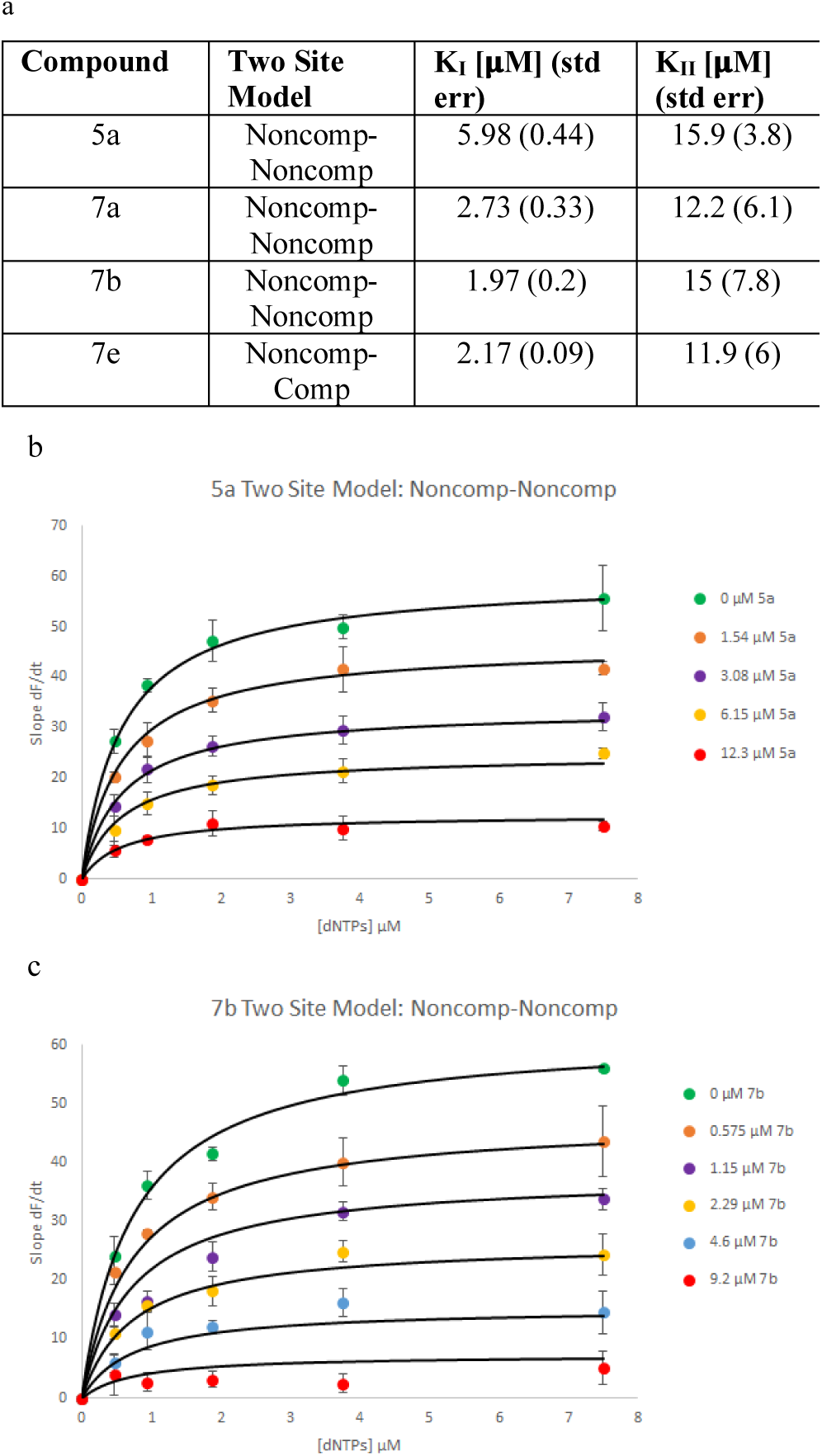
Dynafit results for fitting Mechanism data to two site models. a.) Table summarizes inhibitor equilibrium constants K_I_ and K_II_ for parent compound and three urea derivatives that retained parental activity. 5a, 7a, and 7b data fit best to a two-site model that is noncompetitive at both sites while 7e was noncompetitive at one site and now competitive at the second site b.) Inhibitor mechanism data graph of 5a experimental data (colored circles) plotted with fitted data (solid black lines) from dynafit for a two-site model that is noncompetitive at both sites. c.) Inhibitor mechanism data graph for 7b; a more complicated two-site model used resulted in a better fit of the theoretical data to the experimental data at the highest concentration of 7b used.

## Crystallography work and W512 mutant studies-

**Figure 5*a*** presents an overview of the apPOL structure. The enzyme can replicate a DNA template (polymerase activity) and hydrolyze 2’-deoxynucleotides from the 3’-end of oligonucleotides (exonuclease activity, a function that putatively eliminates miss-incorporated bases during polymerization). Compounds investigated here either inhibit polymerase activity (amide-and urea-linked compounds such as **7a**) or inhibit exonuclease activity (sulfonamide-linked compounds containing electron withdrawing groups on phenyl ring such as compound **6a**). Initial electron density maps revealed significant difference density between the two molecules of apPOL^exo–^ in the asymmetric unit of the P6_5_ crystal form. The electron density, attributed to bound inhibitors, is in proximity to Trp512. Subsequent mutations, W512A and W512F, demonstrated the impact of position 512 on polymerases activity (K_m_ for dNTPs increased 20-fold for W512A) (**Table 2**) and inhibition (K_I_ and K_II_ for W512F changed 6-8fold without a change in K_m_ for dNTPs) (**Table 2**). Further refinement, however, did not result in a satisfactory fit of inhibitor models to the electron density. Subsequent data collection on crystals without inhibitors revealed equivalent electron density and eliminated the possibility of electron density arising from the inhibitor. Rather, the electron density is attributed to low molecular weight components of polyethylene glycol 1000 that localized between the two polypeptide chains of apPOLexo^−^ (**Figure 5b**). Representative structures of apPOLexo^−^ (PDB ID 7SXL; 2.7 Å resolution; 0.19 R-work; 0.22 R-free; 0.97 Molprobity score) and W512F-apPOLexo^−^ (PDB ID 7SXQ; 2.5 Å resolution; 0.20 R-work; 0.22 R-free; 0.99 Molprobity score) were deposited in the Protein Data Base and served as a basis for the modeling of bound inhibitors.

**Figure 5:**
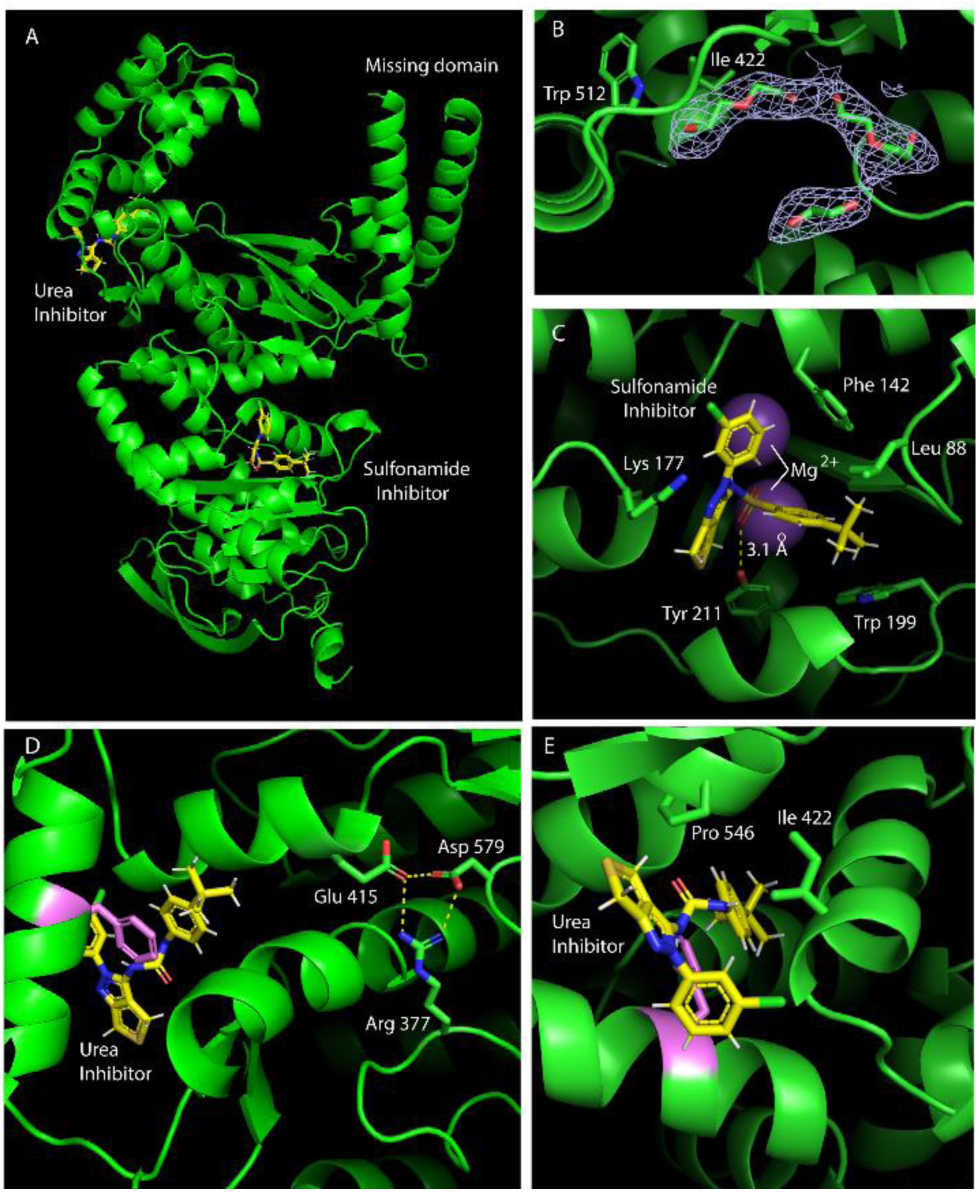
Crystallography images. A.) apPOL displayed with 7a and 6a docked in the W512 binding pocket and exonuclease active site respectively B.) Electron density from small MW fragments of PEG 1000 shown proximal to W512 indicating this regions potential as a hotspot for small molecule binding. C.) apPOL exonuclease domain active site with compound 6a and two Mg^2+^ ions modeled into active site. D.) Another view of apPOL^exo-^ W512F binding pocket displaying its proximity to -1 nucleotide position of the dNTP binding site that is ∼20A from W512. E.) 7a docked into W512F binding pocket showing F512, P546, and I422 forming a gate around the urea-based linker of 7a scaffold.

**Table 2.** Inhibitor equilibrium binding constants for apPOL and W512 mutants (noncompetitive-noncompetitive dual-site); compares the dNTP K_M_ and inhibitor equilibrium binding constants K_I_ and K_II_ between apPOL^exo-^W512 and mutants W512A and W512F.

| Enzyme | dNTPs $K_M$ [$\mu M$] | Two Site Model | **7a** $K_I$ [$\mu M$] (std err) | **7a** $K_{II}$ [$\mu M$] (std err) |
| --- | --- | --- | --- | --- |
| W512 | 0.593 | Noncomp-Noncomp | 2.73 (0.33) | 12.2 (6.1) |
| W512A | 28.8 | Noncomp-Noncomp | 4.65 (0.7) | 6.8 (2.5) |
| W512F | 0.811 | Noncomp-Noncomp | 16.6 (7.3) | 2.4 (1.4) |

Though we were unable to resolve inhibitor bound apPOL, Trp512 mutant studies indicate its involvement in inhibitor binding. Comparing the two crystal structures shows that the indole sidechain of Trp512 packs into a cluster of hydrophobic residues. In contrast, the phenyl sidechain of Phe512 does not pack into the site of the indole moiety of Trp512. Moreover, Ile422 undergoes a ∼120° rotation about its CA-CB bond (χ^1^). The principal result of the W512F mutation is the formation of a pocket containing 5 molecules of water. Our structure activity relationships indicate a critical role for the *t*-butyl phenyl moiety in the inhibition of polymerase activity. Docking compound **7a** into the putative binding site in W512F mutant **(Figure 5d)** shows *t*-butyl phenyl group of **7a** tightly fits into the pocket created by W512F mutation and leaves no empty space. The side chains of Phe512, Ile422 and Pro546 form a gate that closes over the amide and urea linkages **(Figure 5e)**. To provide access to the pocket for entry of the t-butyl-phenyl group, the gate must open transiently. The sulfonamide-linked inhibitor with its additional oxygen atom, sterically interferes with a closed gate, providing potential reason for its lack of inhibition of apPOL polymerase activity.

Additionally, the discovery of exonuclease inhibition by compound **6a** came about unexpectedly as all compounds studied here were derivatives of a parent compound (**5a**) that exhibits no inhibition of exonuclease activity. Structure-activity relationships unequivocally associate exonuclease inhibition with the sulfonamide moiety of compound **6a** (**Table 1**). Modelling **6a** into the exonuclease active site of apPOL (putative binding site) shows that **6a** has a limited number of conformations that avoid close internal contacts and retain favorable stereochemistry. The conformer exhibiting the greatest contact surface with the active site appears in **Figure 5c**. The pose shows one of the oxygen atoms of the sulfonamide moiety bridges the two metals of the active site, as the second oxygen atom (hydrogen bonded to Tyr211) occupies the space taken by the nucleophilic water responsible for the hydrolysis of the substrate (phosphodiester) linkage. The *t*-butyl-phenyl group fills the void for the 3’-ribose and base and the 3-chlorophenyl group occupies space for the base of the second nucleotide residue. And the fused ring system fills the space for the ribosyl and 5’-phosphoryl groups of the second nucleotide, providing a model for exonuclease inhibition.

Efforts from crystallography studies of pfapPOL^exo-^ though unsuccessful in identifying the binding site, still indicated that W512 plays a potential role in enzyme-inhibitor interactions. To test this theory, W512 was mutated to Ala using site-directed mutagenesis (SDM)^33^ with primers P3 and P4 (SI Primer list); mutant protein expression and purification was done the same as wild type. Mutating W512 to Ala resulted in 20-fold increase in dNTP K_M_ with no significant change in k_cat_ indicating W512, which sits ∼20A from polymerase active site, communicates with dNTP binding site via long-range interactions and could contribute significantly to substrate affinity.

To study a mutant that retains the substrate kinetics of the WT W512 protein, a residue that preserves the steric environment that Trp provides, such as Phe, was employed (primers P5 and P6 were used in SDM to generate W512F mutant). The W512F mutant displayed dNTP substrate kinetics similar to WT and was used to reanalyze IC_50_ and inhibition mechanism data for compound **7a**. The **7a** IC_50_ for W512F protein was within 2-fold of the WT value, and mechanism data fit consistently with a nonidentical, dual-site mechanism that is noncompetitive at both sites. Interestingly, the W512F mutant suggested the two binding sites are no longer independent and are rather coupled as exemplified by the significant increase in affinity going from K_I_ (16 µM) to K_II_ (2 µM) (**Table 2**). This binding synergism observed for compound **7a** against W512F suggests the W512F mutant promotes **7a** coupling between the two binding sites while W512 and W512A variants promote **7a** binding mechanisms that could be independent or antagonistic.

In summary, the data presented here establishes apPOL as a target for drug development. The parent compound, along with the amide-and urea-linked inhibitors clear *P. falciparum* infections from human blood cultures at dosage levels that correspond to 50% inhibition levels of polymerase activity. The inhibition kinetics data are consistent with two binding sites that are independent in the WT enzyme and synergistic in the W512F mutant. We provide evidence suggesting that one of the inhibitor binding sites is located in a hydrophobic pocket created when Trp512 rotates out of its accustomed position. This pocket is observed in crystals grown in 10% DMSO or by the W512F mutation and appears to be filled by low molecular weight compounds derived from PEG. The location of the second inhibitor binding site is unclear and will require further investigation.

The family of compounds studied here offers promise as leads for the development of new antimalarial drugs. In the case of sulfonamide-linked inhibitors, the path toward improved inhibitors is clear, but the biological consequences of potent inhibition of exonuclease activity are uncertain. Methods have been developed that enable replacement of the endogenous apPOL with an exonuclease deficient mutant to determine whether the loss of exonuclease activity impairs the viability of *P. falciparum*. Amide- and urea-linked inhibitors clearly impact the viability of *P. falciparum* in biological assays, but the binding loci for such inhibitors remains elusive. Further progress will rely in part on crystals of an apPOLexo^−^ complex with DNA, and the use of that complex to identify binding sites for polymerase-directed inhibitors.

## ASSOCIATED CONTENT

Supporting Information: The Supporting Information is available free of charge at https://pubs.acs.org/

Additional synthetic schemes, detailed synthetic protocols, detailed compound characterization, protocols for in vitro assays, protocols for crystallographic studies, protocols for site-directed mutagenesis.

### Accession Codes

Structure data associated with this study have been deposited to the RCSB Protein Data Bank (http://www.rcsb.org) and can be accessed with the following codes: apPOLexo^−^ PDB ID 7SXL and W512F-apPOLexo^−^ PDB ID 7SXQ. Authors will release the atomic coordinates and experimental data upon article publication

## AUTHOR INFORMATION

Authors:

**Pratik R. Chheda**: Division of Medicinal and Natural Products Chemistry, Department of Pharmaceutical Sciences and Experimental Therapeutics, University of Iowa, Iowa City, Iowa 52242, United States.

**Nicholas Nieto**: The Roy J. Carver Department of Biochemistry, Biophysics, and Molecular Biology, Iowa State University, Ames, IA, 50011, USA.

**Supreet Kaur**: The Roy J. Carver Department of Biochemistry, Biophysics, and Molecular Biology, Iowa State University, Ames, IA, 50011, USA.

**John M. Beck**: The Roy J. Carver Department of Biochemistry, Biophysics, and Molecular Biology, Iowa State University, Ames, IA, 50011, USA; Department of Biomedical Sciences, Iowa State University, Ames, Iowa, 50011, USA.

**Josh R. Beck**: Department of Biomedical Sciences, Iowa State University, Ames, Iowa, 50011, USA.

**Richard Honzatko**: The Roy J. Carver Department of Biochemistry, Biophysics, and Molecular Biology, Iowa State University, Ames, IA, 50011, USA.

## Author Contributions

P.R.C. designed research, carried out chemical synthesis and co-wrote the paper. N.N. designed and carried out in vitro experiments, crystallographic experiments, and co-wrote the paper. S.K. carried out in vitro experiments and crystallographic experiments. J.M.B. and J.R.B. carried out parasite killing experiments, R.H. carried out crystallographic and modelling experiments. S.W.N and R.J.K. designed and directed research, secured funding and co-wrote the paper, with assistance from all other authors.

## Notes

The authors declare no competing financial interest.

## ACKNOWLEDGMENTS

The work in the laboratory of R.J.K. was supported by the John L. and Carol E. Lach Chair in Drug Delivery Technology and University of Iowa College of Pharmacy research stimulation funds. The work in the laboratory of S.W.N. was supported by NIH grant R21AI127622. P.R.C acknowledges the support of a predoctoral fellowship from the University of Iowa Center for Biocatalysis and Bioprocessing affiliated with the NIH sponsored Predoctoral Training Program in Biotechnology (T32 GM008365).

## ABBREVIATIONS USED

apPOL: Apicoplast DNA polymerase
*P. falciparum*: Plasmodium falciparum
IPP: Isoprenoids
TFA: trifluoroacetic acid
ACN: acetonitrile
MeOH: methanol
DCM: dichloromethane
DMEM: Dulbecco’s modified Eagle’s medium
SHM: Seahorse medium
BSA: bovine serum albumin

## REFERENCES

1. de Koning-Ward, T. F.,, Dixon, M. W. A.,, Tilley, L.,, Gilson, P. R., Plasmodium species: master renovators of their host cells. Nature Reviews Microbiology 2016, 14, 494–507.

2. Organization, W. H., World Malaria Report 2017. 2017.

3. White, N. J.; Pukrittayakamee, S.; Hien, T. T.; Faiz, M. A.; Mokuolu, O. A.; Dondorp, A. M., Malaria. The Lancet 2014, 383 (9918), 723–735.

4. Gazzinelli, R. T.; Kalantari, P.; Fitzgerald, K. A.; Golenbock, D. T., Innate sensing of malaria parasites. Nature Reviews Immunology 2014, 14, 744–757.

5. Gogtay, N.; Kannan, S.; Thatte, U. M.; Olliaro, P. L.; Sinclair, D., Artemisinin-based combination therapy for treating uncomplicated Plasmodium vivax malaria. Cochrane Database of Systematic Reviews 2013, (10), CD008492. doi: 10.1002/14651858.CD008492.pub3.

6. Uwimana, A.; Umulisa, N.; Venkatesan, M.; Svigel, S. S.; Zhou, Z.; Munyaneza, T.; Habimana. R. M.; Rucogoza, A.; Moriarty, L. F.; Sandford, R.; Piercefield, E.; Goldman, I.; Ezema, B.; Talundzic, E.; Pacheco, M. A.; Escalante, A. A.; Ngamije, D.; Mangala, J. N.; Kabera, M.; Munguti, K.; Murindahabi, M.; Brieger, W.; Musanabaganwa, C.; Mutesa, L.; Udhayakumar, V.; Mbituyumuremyi, A.; Halsey, E. S.; Lucchi, N. W., Association of Plasmodium falciparum kelch13 R561H genotypes with delayed parasite clearance in Rwanda: an open-label, single-arm, multicentre, therapeutic efficacy study. Lancet Infect Dis. 2021, 21 (8), 1120–1128.

7. Balikagala, B.; Fukuda, N.; Ikeda, M.; Katuro, O. T.; Tachibana, S. I.; Yamauchi, M.; Opio, W.; Emoto, S.; Anywar, D. A.; Kimura, E.; Palacpac, N. M. Q.; Odongo-Aginya, E. I.; Ogwang, M.; Horii, T.; Mita, T., Evidence of Artemisinin-Resistant Malaria in Africa. N Engl J Med. 2021, 385 (13), 1163–1171.

8. White, N. J., Antimalarial drug resistance. The Journal of Clinical Investigation 2004, 113 (8), 1084–1092.

9. Wongsrichanalai, C.; Sibley, C. H., Fighting drug-resistant Plasmodium falciparum: the challenge of artemisinin resistance. Clinical microbiology and infection: the official publication of the European Society of Clinical Microbiology and Infectious Diseases 2013, 19 (10), 908–916.

10. Waller, R. F.; McFadden, G. I., The apicoplast: a review of the derived plastid of apicomplexan parasites. Current issues in molecular biology 2005, 7 (1), 57–79.

11. Striepen, B., The apicoplast: a red alga in human parasites. Essays in biochemistry 2011, 51, 111–125.

12. Seeber, F.; Soldati-Favre, D., Metabolic pathways in the apicoplast of apicomplexa. International review of cell and molecular biology 2010, 281, 161–228.

13. Yeh, E.; DeRisi, J. L., Chemical rescue of malaria parasites lacking an apicoplast defines organelle function in blood-stage Plasmodium falciparum. PLoS biology 2011, 9 (8), e1001138.

14. Kuschner, R.A.; Heppner, D.G.; Andersen, S.L.; Wellde, B.T.; Hall, T; Schneider, I; Ballou, W.R.; Foulds, G; Sadoff, J.C.; Schuster, B; et al. Azithromycin prophylaxis against a chloroquine-resistant strain of Plasmodium falciparum. Lancet. 1994 Jun 4;343(8910):1396–7. doi: 10.1016/s0140-6736(94)92526-7.

15. Seaberg, L.S.; Parquette, A.R.; Gluzman, I.Y.; Phillips, G.W. Jr.; Brodasky, T.F.; Krogstad, D.J., Clindamycin Activity Against Chloroquine-Resistant Plasmodium falciparum. Journal of Infectious Diseases 1984, 150 (6), 904–911.

16. Rieckmann, K.H.; McNamara, J.V.; Willerson, D.; Kass, L.; Frischer, J.; Carson, P.E., Effects of tetracycline against chloroquine-resistant and chloroquine-sensitive Plasmodium falciparum. Am J Trop Med Hyg 1971, 20, 811815.

17. Dahl, E. L.; Rosenthal, P. J., Multiple antibiotics exert delayed effects against the Plasmodium falciparum apicoplast. Antimicrobial agents and chemotherapy 2007, 51 (10), 3485–3490.

18. Pradel, G.; Schlitzer, M., Antibiotics in malaria therapy and their effect on the parasite apicoplast. Current molecular medicine 2010, 10 (3), 335–349.

19. Reguera, R. M.; Redondo, C. M.; Gutierrez de Prado, R.; Perez-Pertejo, Y.; Balana-Fouce, R., DNA topoisomerase I from parasitic protozoa: a potential target for chemotherapy. Biochimica et biophysica acta 2006, 1759 (3-4), 117–131.

20. Ralph, S. A.; van Dooren, G. G.; Waller, R. F.; Crawford, M. J.; Fraunholz, M. J.; Foth, B. J.; Tonkin, C. J.; Roos, D. S.; McFadden, G. I., Tropical infectious diseases: metabolic maps and functions of the Plasmodium falciparum apicoplast. Nature reviews. Microbiology 2004, 2 (3), 203–216.

21. Lindner, S. E.; Llinas, M.; Keck, J. L.; Kappe, S. H., The primase domain of PfPrex is a proteolytically matured, essential enzyme of the apicoplast. Molecular and biochemical parasitology 2011, 180 (2), 69–75.

22. Seow, F.; Sato, S.; Janssen, C. S.; Riehle, M. O.; Mukhopadhyay, A.; Phillips, R. S.; Wilson, R. J.; Barrett, M. P., The plastidic DNA replication enzyme complex of Plasmodium falciparum. Molecular and biochemical parasitology 2005, 141 (2), 145–153.

23. Schoenfeld, T. W.; Murugapiran, S. K.; Dodsworth, J. A.; Floyd, S.; Lodes, M.; Mead, D. A.; Hedlund, B. P., Lateral gene transfer of family A DNA polymerases between thermophilic viruses, aquificae, and apicomplexa. Molecular biology and evolution 2013, 30 (7), 1653–1664.

24. Aurrecoechea, C.; Brestelli, J.; Brunk, B. P.; Dommer, J.; Fischer, S.; Gajria, B.; Gao, X.; Gingle, A.; Grant, G.; Harb, O. S.; Heiges, M.; Innamorato, F.; Iodice, J.; Kissinger, J. C.; Kraemer, E.; Li, W.; Miller, J. A.; Nayak, V.; Pennington, C.; Pinney, D. F.; Roos, D. S.; Ross, C.; Stoeckert, C. J., Jr.; Treatman, C.; Wang, H., PlasmoDB: a functional genomic database for malaria parasites. Nucleic acids research 2009, 37 (Database issue), D539–43.

25. Milton, M. E.; Choe, J. Y.; Honzatko, R. B.; Nelson, S. W., Crystal Structure of the Apicoplast DNA Polymerase from Plasmodium falciparum: The First Look at a Plastidic A-Family DNA Polymerase. Journal of molecular biology 2016, 428 (20), 3920–3934.

26. Miller, M. E.; Parrott, E. E.; Singh, R.; Nelson, S. W., A High-Throughput Assay to Identify Inhibitors of the Apicoplast DNA Polymerase from Plasmodium falciparum. Journal of biomolecular screening 2014, 19 (6), 966–972.

27. Baraldi, P. G.; El-Kashef, H.; Manfredini, S.; Pineda de las Infantas, M. J.; Romagnoli, R.; Spalluto, G., A Mild One-Pot Synthesis of Thieno[3,4-c]pyrazoles and Their Conversion into Pyrazole Analogs of o-Quinodimethane. Synthesis 1998, 1998 (09), 1331–1334.

28. Information, N.C.F.B. Source=ChEMBL, AID=781324 https://pubchem.ncbi.nlm.nih.gov/bioassay/781324 (Accessed May 22,2020)

29. Reha-Krantz, L.J. The use of 2-aminopurine fluor, escence to study DNA polymerase function. Methods Mol Biol. 2009, 521, 381–396.

30. Kuzmic, P., DynaFit--a software package for enzymology. Methods in enzymology 2009, 467, 247–280.

31. Burnham, K.P. D. R. A., Model Selection and Multimodel Inference. 2ed.; Springer-Verlag New York: 2002.

32. Schwarz, G., Estimating the Dimension of a Model. The Annals of Statistics 1978, 6 (2), 461–464.

33. Wang, W.; Malcolm, B.A., Two-stage PCR protocol allowing introduction of multiple mutations, deletions and insertions using QuickChange Site-Directed Mutagenesis. BioTechniques 1999, 26 (4), 680–682.

