## Supplemental file1 for "Promising Antimalarials targeting Apicoplast DNA Polymerase from *Plasmodium falciparum*"

Table of Contents

| Title | Page number |
|---|---|
| Protocols for in vitro assays | S2 |
| Synthetic Procedures | S7 |
| $^{1}$H NMR Spectra for all intermediates and final compounds | S23 |
| References | S41 |

Protocols for in vitro assays:

**Expression and Purification of apPOL–** Steady-state kinetics and x-ray crystallography were performed with exonuclease deficient apPOL polymerase ($Asp^{53}$→$Asn/Glu^{55}$→Gln). *E. coli* BL21 (DE3) cells were transformed with the pSUMO-$apPOL^{exo-}$ vector that supplies an hexahistidine tagged SUMO fusion protein at the N-terminus of apPOL. A single colony was obtained from a LB/kanamycin agar plate and transferred to flasks containing 100 ml LB/kanamycin medium. After shaking the flasks for 16h at 37°C, 10-ml aliquots of the LB starter culture were transferred to two flasks each containing 1-liter of LB/kanamycin. The flasks were shaken (225 rpm) at 37°C until an $A_{600}$ of 0.8 was reached. The flasks were then cooled to 16°C, after which apPOL expression was induced by adding 200 μM (final) isopropyl β-D-1-thiogalactopyranoside (IPTG). After 16 h with constant shaking at 16°C, the cells were collected by centrifugation at 4000 rpm for 10 minutes. The cell pellet was then resuspended in 20 mL nickel binding buffer (20 mM Tris, 500 mM NaCl, 5 mM imidazole, 10% Glycerol, pH 8, 4°C) and passed through a homogenizer three times at 12,000 psi. The lysate was then centrifuged at 17000 rpm for 1 hour and the supernatant was loaded onto a 3 ml Ni-agarose column. The column was washed with 200 ml of binding buffer and then with 100 ml of high salt wash buffer (20 mM Tris, 1 M NaCl, 5 mM imidazole and 10% glycerol, pH 8.0, 4°C) followed by 30 ml of low salt wash buffer (20 mM Tris-HCl, 500 mM NaCl, 20 mM imidazole and 10% glycerol, pH 8.0, 4°C). Then the protein was eluted with 20 mM Tris-HCl, 400 mM NaCl, and 150 mM imidazole. The protein was then further purified using a 150 ml size exclusion column containing TOYOPEARL HW-50F resin, which was previously equilibrated with 20 mM Tris-HCl, 400 mM NaCl, pH 8.0 at 4°C. Eluted fractions were analyzed using a standard Bradford assay, followed by SDS-PAGE. Fractions containing pure apPOL were pooled and incubated with SUMO protease for 16 hours at 4°C at a 1:50 protein to SUMO protease ratio. Following removal of the SUMO tag, the protein solution was passed through the nickel-NTA-agarose column, which retains the unwanted SUMO fragment and the SUMO protease but allows tag-free apPOL to pass through. Protein purity was confirmed by SDS-PAGE using Coomassie Blue stain and concentrated to 20 mg/ml using an extinction coefficient of $\varepsilon_{280}$ =56750 $M^{-1}$ $cm^{-1}$. Protein destined for crystallization was used immediately, whereas protein used for activity assays was then brought to 20% glycerol and flash-frozen in liquid nitrogen and stored at 193 K.

**Kinetic Experiments–** Measurement of apPOL$^{exo-}$ DNA polymerase activity employed a DNA hairpin substrate[1] in Polymerase Reaction Buffer (PRB) 20 mM Tris-acetate, 10 mM magnesium acetate, 50 mM potassium acetate, pH 8.0 at 25 °C. Mechanisms of inhibition were determined by measuring polymerase activity at 20nM enzyme, 50nM DNA comprised of a 33nt primer within hairpin and double stranded region with 3'Cy3 dye and 5'quencher-polymerase extends primer displacing double stranded region releasing dye from quencher to produce signal-and several concentrations of an equimolar mixture of dNTPs ranging from 0.5-8$K_M$ increasing by two-fold. This assay was tested for each concentration of inhibitor ranging from 0.25-4 $IC_{50}$, increasing by two-fold, including a 0μM inhibitor control. Assays were carried out in PRB, where enzyme plus DNA and inhibitor were incubated for two minutes before reactions were initiated by addition of dNTPs. Time courses were collected in real-time using a Cary Eclipse Fluorescence Spectrophotometer with excitation and emission wavelengths of 545 nm and 570 nm, respectively. Initial velocities were plotted against dNTP concentration to generate a set of reaction saturation curves each representing a different concentration of inhibitor. Using Dynafit software[2], the data was fit to multiple models of inhibition including competitive, noncompetitive, mixed, and uncompetitive for a single binding site inhibitor mechanism. The model discrimination analysis component of the software was used to determine which model the data fit best to; when model discrimination chose more than one possible model the best-fit parameters ΔAIC and ΔBIC were used to discriminate which model the data was most consistent with. Results from the one site model fitting suggested possibility of a second binding site therefore, the data was fit to a two-site inhibitor binding model where each site was either competitive, noncompetitive, uncompetitive, mixed, or different combinations of all four. For the inhibition assays, the inhibitor was incubated with apPOL$^{exo-}$ for 2 minutes prior to the addition of the DNA substrate and dNTPs for the time course of at least 2 minutes. Initial velocities were calculated from the slope of the linear portion of the time course and the data were fit to a variety of inhibition mechanisms using the Dynafit software.[2]

**Exonuclease assays:** Wild type apPOL exonuclease activity was measured in the presence and absence of each inhibitor using a 2'aminopurine blunt ended DNA substrate in 20 mM Tris-HCl, 10 mM magnesium acetate, 50 mM potassium acetate, and 0.1 mg/mL BSA pH 8.0 at 25 °C. Signal was produced by release of the 2'amino purine at the 3' end of the substrate and measured in real time using a Cary Eclipse Fluorescence Spectrophotometer with excitation and emission wavelengths of 310 nm and 375nm, respectively. Single concentrations of each inhibitor were incubated with WTapPOL for 2 minutes and DNA was added to initiate the reaction. Linear regions of the reaction time courses were used to measure the initial velocity and calculate the specific activity of the enzyme in the presence and absence of inhibitor. All inhibitor reactions were done in triplicate with a no inhibitor control, and specific activity values are displayed as averages with standard deviations. Two of the most potent inhibitors displaying >90% decrease in specific activity of enzyme compared to no inhibitor control were subjected to $IC_{50}$ evaluation using the same assay.

**Parasite killing assays:** Inhibitors were subjected for parasite killing growth assays utilizing acridine orange DNA/RNA binding dye and flow cytometry as described.[3] Briefly, full $IC_{50}$ killing curves were measured by plating inhibitors and DMSO controls in 96 well plates at 1% the volume of $NF54^{attb}$ parasite cultures added. Parasites were plated at 1% parasitemia and allowed to grow for 72 hrs. Flow cytometer measurements were taken at 48- and 72-hour time marks. Inhibitor growths were normalized to controls and percentage growth values were plotted against the log[Inhibitor] to extract the $EC_{50}$. IPP rescue assay was initially evaluated on parent compound 5a to confirm apicoplast targeting, 5a was plated at 20 µM (at 1% of parasite volume as done before) and incubated with parasites under 4 conditions in triplicate: regular RPMI media, regular RPMI media + 200µM IPP, regular RPMI media + Minocyclin, and RPMI media + Minocyclin + IPP. Parasitemia measurements were taken at 72- and 96-hr time points and parasite growth for each condition was normalized to DMSO control and compared between two biological replicates-each of which contained triplicate technical replicates.

**Generating pfapPOLexo- W512 mutants-** Primers were purchased from either IDT or Iowa State University DNA Facility. Primers for generating exonuclease deficient apPOL were designed as described.[4] Mutants of pfapPOL$^{exo-}$ were generated by a two-stage PCR protocol using Quick Change site directed mutagenesis as described.[5] Briefly, 50uL single primer extension reactions using 10uM forward or reverse primer were conducted using 1X Phusion Polymerase High Fidelity Master Mix (1 U of Phusion High Fidelity Polymerase final), and 100ng of pSUMO-pfapPOLexo- as a template for 10 PCR cycles-these reactions result in extended primers that possess longer regions complementary to the vector template and aid in prevention of primer dimer formation. A second 50uL PCR reaction was subsequently done using equal volumes of the forward and reverse single primer extension reactions with an additional 1U of Phusion polymerase and ran for the standard 35 PCR cycles. This second PCR reaction mixture was then subjected to overnight DPN1 digestion and transformed into E. Cloni chemically competent cells and plated on LB-agar plates containing 30ug/mL Kanamycin. Colonies were selected and plasmids extracted, using a Qiagen Miniprep Kit, and sequenced for correct mutation using Iowa State University DNA Facility.

**Primer List**

| **Primer** | **Sequence** |
|---|---|
| **P1 (POM1exo-F)** | 5’TCAAATACTGTGGTCTGaATATTcAAACCACGGGCCTGGAA3’ |
| **P2 (POM1exo-R)** | 5’TTCCAGGCCCGTGGTTTgAATATtCAGACCACAGTATTTGA3’ |
| **P3 (W512AFor)** | 5’CGAACATTACAAAGGCATCTACAAAgcGCACAATCAGGTTAAAC3’ |
| **P4 (W512ARev)** | 5’GTTTAACCTGATTGTGCgcTTTGTAGATGCCTTTGTAATGTTCG3’ |

| | |
|---|---|
| **P5 (W512F For)** | 5'CAAAGGCATCTACAAATtcCACAATCAGGTTAAAC3' |
| **P6 (W512F Rev)** | 5'GTTTAACCTGATTGTGgaATTTGTAGATGCCTTTG3' |

Note: Mutations are shown in lower case

**Crystallization and Data Collection**– In general, the inhibitors exhibited limited solubility in aqueous buffers (as determined visually). As the urea-based derivatives appeared to be the most soluble in aqueous buffers, we initiated the crystallization trials with **7a**. The protein (20 mg/ml) was passed through a 0.22 μm cellulose acetate spin filter after which powdered inhibitor was added incrementally to the point of saturation and incubated at room temperature for 10 minutes. Excess (undissolved) inhibitor was then removed by centrifugation. A Mosquito crystallization robot was used to dispense protein and the crystallization screening solution onto 96-well sitting-drop plates. From the 400 conditions screened, a single condition produced crystals. The result was replicated and optimized manually by the method hanging-drop vapor diffusion. The final crystallization condition used was 0.1 M sodium citrate tribasic dihydrate pH 5.5, 400 mM NaCl, and 30% w/v polyethylene glycol 1000. The addition of 400 mM NaCl was necessary to prevent the crystals from dissolving. After approximately 7 days of incubation at 22 $^{o}$C, the crystal was looped and immediately plunged into liquid nitrogen for storage and subsequent data collection. Diffraction data were collected at 100 K using a Pilatus 3 6M detector on Advanced Photon Source beamline 23-ID-D. 600 frames were collected using an oscillation range of 0.2° with a 20 μm beam at a crystal to detector distance of 450 mm. HKL-3000 was used for data processing. Molecular replacement was performed using the PHENIX software suite with 5DKU as a model, followed by model building and refinement using Phenix-refine and Coot.

### Synthetic Procedures

**General Chemical Synthesis Information:** All commercially available reagents and solvents were used directly without further purification unless otherwise noted. Reactions were monitored either by thin-layer chromatography (carried out on silica plates, silica gel 60 $F_{254}$, Merck) and visualized under UV light. Flash chromatography was performed using silica gel 60 as stationary phase performed under positive air pressure. $^1$H and $^{19}$F NMR spectra were recorded in $CDCl_3$ or $d_6$-DMSO on a Bruker Avance spectrometer operating at 300 MHz at ambient temperature. All peaks are reported in ppm on a scale downfield from TMS and using the residual solvent peak in $CDCl_3$ (H $\delta$ = 7.26) or TMS ($\delta$ = 0.0) as an internal standard. Data for $^1$H NMR are reported as follows: chemical shift (ppm, scale), multiplicity (s =singlet, d = doublet, t = triplet, q = quartet, m = multiplet and/or multiplet resonances, dd = double of doublets, dt = double of triplets, br = broad), coupling constant (Hz), and integration. All high-resolution mass spectra (HRMS) were measured on Waters Q-Tof Premier mass spectrometer using electrospray ionization (ESI) time-of-flight (TOF). All analytical HPLC analyses were performed using a Shimadzu system comprised of an LC-20AT pump, DGU-14A degasser, CBM-20A system controller, and a SPD-M10AVP photodiode array detector. The HPLC was operated using Shimadzu Client/Server Version 7.4 software installed on a Dell Optiplex GX400 PC. The column used was Restek Allure PFP Propyl unless otherwise indicated. The mobile phase was composed of water buffered with 0.1 % trifluoroacetic acid (TFA) and acetonitrile also buffered with 0.1 % TFA. The HPLC analyses were obtained by running a gradient of 5 % - 95 % buffered acetonitrile over 30 minutes with a flow rate of 1.000 mL/min. Select final products were purified using reverse-phase semi-preparative HPLC. The Shimadzu system was composed of the following parts: two LC-10AT pumps (one for each solvent), SPD-M10AVP photodiode array detector, and an SCL-10AVP system controller. The HPLC system was operated using Shimadzu EZStart Version 7.4 control software installed on a Dell Optiplex 755 PC. The stationary phase used for separation was a Phenomenex Luna PFP (2) reverse phase column. The mobile phase was composed of water buffered with 0.1 % TFA and acetonitrile also buffered with 0.1 % TFA. Various gradients were used for separation of products, but most commonly a 5 % - 95 % buffered acetonitrile gradient over 40 minutes was used. Reaction progress was monitored utilizing analytical HPLC (methods described above) or thin layer chromatography (TLC). The TLC plates used were glass-backed 0.25 mm TLC Silica Gel 60 plates containing fluorescence indicator F264 (EMD Sciences). The TLC plates were visualized by one or more of the following three methods: (1) UV absorption/fluorescence (254 nm and 366 nm), (2) Iodine chamber, (3) cerium molybdate stain with subsequent heating. Flash chromatography was used to purify many intermediate compounds. This was accomplished using Silica Gel 60 (particle size 0.040 – 0.063 mm; 230 – 400 mesh ASTM) and various solvent systems. The solvent ratios described are volume/volume. Removal of organic solvents was accomplished by first drying the organic solvent over $Na_2SO_4$ before concentrating on a rotary evaporator. Aqueous solutions were concentrated by using either a rotary evaporator or lyophilizer.

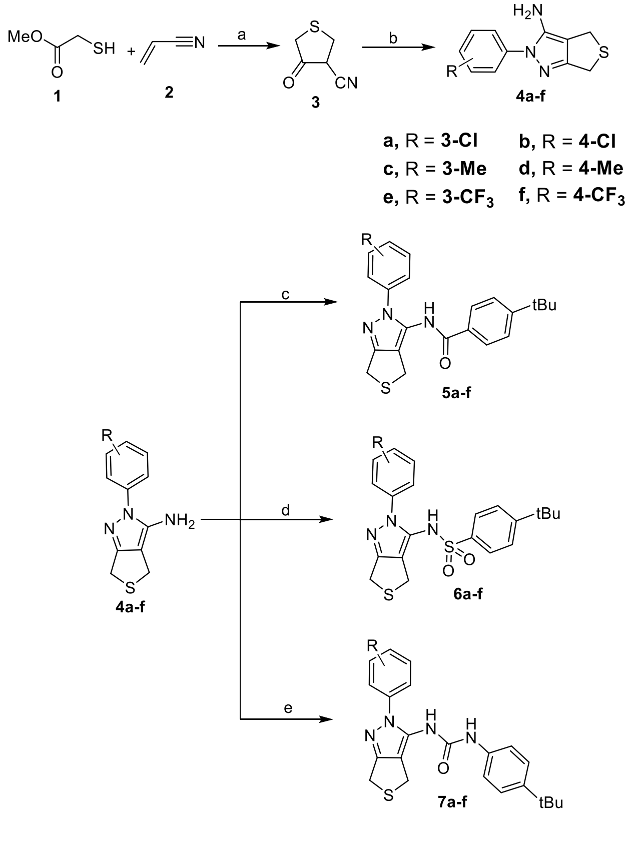


***4-oxotetrahydrothiophene-3-carbonitrile*** (**3**): To a dried round bottom flask was added sodium methoxide (5.4M solution in methanol) (3.3 mmol, 1.2 equiv.) and allowed to stir at 0 °C for 15 mins, to which was added methyl mercaptoacetate (2.83 mmol, 1 equiv.) dropwise at 0 $^{o}$C. The solution was allowed to stir for 15 mins. To this was added acrylonitrile (2.83 mmol, 1 equiv.) and the reaction was then stirred at ambient temperature for 3 hours. Methanol was then removed, and the residue was dissolved in water and extracted with diethyl ether (50 ml) to remove residual ester. The aqueous phase was then acidified with 4N hydrochloric acid and extracted with diethyl ether (2x50ml). The organic phases were collected, dried over $Na_2SO_4$ and concentrated *in vacuo* to give 4-oxotetrahydrothiophene-3-carbonitrile as yellow liquid (151 mg, 42%), which was used for next step without further purification. HRMS (ESI$^-$) for $C_5H_4NOS$ (M-H$^+$) 126.0023 (Negative mode).

**General Experimental Procedure A**: ***2-(3-chlorophenyl)-2,6-dihydro-4H-thieno[3,4-c] pyrazol-3-amine* (4a): To a solution of 4-oxotetrahydrothiophene-3-carbonitrile** (**3**) (30 mg, 0.24mmol, 1 equiv.) in ethanol, was added 3-chlorophenyl hydrazine (43 mg, 0.24mmol, 1 equiv.) and the resulting solution was refluxed for 6 hours. Once the reaction was complete, ethanol was removed under vacuum and the residue was dissolved in ethyl acetate which was washed with 5% NaOH solution. The organic layers were collected, concentrated and the resulting crude product was purified by column chromatography eluting in 5:1:: hexanes:ethylacetate, yielding 2-(3-chlorophenyl)-2,6-dihydro-4H-thieno[3,4-c] pyrazol-3-amine (**4a**) as a white solid (37 mg, 63%). $^1$H NMR (300 MHz, $CDCl_3$): δ 7.62 (t, *J* = 2.0 Hz, 1H), 7.51 – 7.45 (m, 1H), 7.42 (t, *J* = 7.8 Hz, 1H), 7.36 – 7.30 (m, 1H), 3.99 (d, *J* = 1.2 Hz, 2H), 3.81 (d, *J* = 1.2 Hz, 2H), 3.77 (s, 2H). HRMS (ESI) m/z: [M + H]$^+$ Calcd for $C_{11}H_{11}ClN_3S$ 252.0362, found 252.0358. Retention factor (TLC)=0.48(4:1::Hexanes:ethylacetate).

***2-(4-chlorophenyl)-2,6-dihydro-4H-thieno[3,4-c] pyrazol-3-amine*** (**4b**): Synthesized from 4-oxotetrahydrothiophene-3-carbonitrile (**3**) (30 mg, 0.24mmol, 1 equiv.) and 4-chlorophenyl hydrazine (42 mg, 0.24mmol, 1 equiv.) following General Experimental Procedure A to provide intermediate **4b** as cream solid (39 mg, 65% yield). $^1$H NMR (300 MHz, $CDCl_3$): δ 7.56-7.40 (m, 4H), 3.99 (m, 2H), 3.81 (m, 2H),

3.73 (bs,2H). HRMS (ESI) m/z: $[M + H]^+$ Calcd for $C_{11}H_{11}ClN_3S$ 252.0362, found 252.0365. Retention factor (TLC)=0.5(4:1::Hexanes:ethylacetate).

***2-(3-methylphenyl)-2,6-dihydro-4H-thieno[3,4-c] pyrazol-3-amine*** (**4c**) (PC-01-162): Synthesized from 4-oxotetrahydrothiophene-3-carbonitrile (**3**) (30 mg, 0.24mmol, 1 equiv.) and 3-methylphenyl hydrazine (35 mg, 0.24mmol, 1 equiv.) following General Experimental Procedure A to provide intermediate **4c** as tan solid (30 mg, 54%). $^1$H NMR (300 MHz, $CDCl_3$) δ 7.34 (m, 3H), 7.17 (d, *J* = 7.3 Hz, 1H), 3.99 (s, 2H), 3.80 (s, 2H), 3.73 (bs, 2H), 2.41 (s, 3H). HRMS (ESI) m/z: $[M + H]^+$ Calcd for $C_{12}H_{14}N_3S$ 232.0908, found 232.0922. Retention factor (TLC)=0.40 (4:1::Hexanes:ethylacetate).

***2-(4-methylphenyl)-2,6-dihydro-4H-thieno[3,4-c] pyrazol-3-amine*** (**4d**) (PC-01-172C): Synthesized from 4-oxotetrahydrothiophene-3-carbonitrile (**3**) (30 mg, 0.24mmol, 1 equiv.) and 3-methylphenyl hydrazine (35 mg, 0.24mmol, 1 equiv.) following General Experimental Procedure A to provide intermediate **4d** as a tan solid (33 mg, 58%). $^1$H NMR (300 MHz, $CDCl_3$) δ 7.40 (d, *J* = 8.4 Hz, 2H), 7.28 (t, *J* = 4.0 Hz, 2H), 4.00 (m, 12), 3.82 (m, 2H), 3.72 (s, 2H), 2.41 (s, 3H). HRMS (ESI) m/z: $[M + H]^+$ Calcd for $C_{12}H_{14}N_3S$ 232.0908, found 232.0903. Retention factor (TLC)=0.40 (4:1::Hexanes:ethylacetate).

***2-(3-trifluoromethylphenyl)-2,6-dihydro-4H-thieno[3,4-c] pyrazol-3-amine*** (**4e**) (PC-01-156): Synthesized from 4-oxotetrahydrothiophene-3-carbonitrile (**3**) (30 mg, 0.24mmol, 1 equiv.) and 3-trifluoromethylphenyl hydrazine (44 mg, 0.24mmol, 1 equiv.) following General Experimental Procedure A to provide intermediate **4e** as a yellow solid (49 mg, 72%). $^1$H NMR (300 MHz, $CDCl_3$): δ 7.96-7.90 (m, 1H), 7.81-7.78 (m, 1H), 7.63-7.61 (m, 2H), 4.07-4.00 (m, 2H), 3.83-3.82 (m, 2H), 3.76 (bs,2H). $^{19}$F NMR (282 MHz, $CDCl_3$) δ -62.69 (s). HRMS (ESI) m/z: $[M + H]^+$ Calcd for $C_{12}H_{11}F_3N_3S$ 286.0626, found 286.0620. Retention factor (TLC)=0.30 (4:1::Hexanes:ethylacetate).

***2-(4-trifluoromethylphenyl)-2,6-dihydro-4H-thieno[3,4-c] pyrazol-3-amine*** (**4f**) (PC-01-172B): Synthesized from 4-oxotetrahydrothiophene-3-carbonitrile (**3**) (30 mg, 0.24mmol, 1 equiv.) and 4-trifluoromethylphenyl hydrazine (44 mg, 0.24mmol, 1 equiv.) following General Experimental Procedure A to provide intermediate **4f** as a yellow solid (46 mg, 69%). $^{1}$H NMR (300 MHz, $CDCl_3$) δ 7.75 (s, 4H), 4.00 (s, 2H), 3.82 (s, 2H), 3.79 (bs, 2H). $^{19}$F NMR (282 MHz, $CDCl_3$) δ -62.37 (s). HRMS (ESI) m/z: $[M + H]^+$ Calcd for $C_{12}H_{11}F_3N_3S$ 286.0626, found 286.0626. Retention factor (TLC)=0.20 (4:1::Hexanes:ethylacetate).

**General Experimental Procedure B**: ***4-(tert-butyl)-N-(2-(3-chlorophenyl)-2,6-dihydro-4H-thieno[3,4-c]pyrazol-3-yl)benzamide*** (**5a**) (PC-01-150 & PC-01-171D): To an oven dried round bottom flask was added 2-(3-chlorophenyl)-2,6-dihydro-4H-thieno[3,4-c] pyrazol-3-amine (**4a**) (90 mg, 0.35mmol, 1 equiv.) followed by anhydrous dichloromethane. To this was added di-isopropyl amine (80 ml, 0.43mmol, 1.2 equiv.) and the resulting solution was allowed to stir at 0 °C for 15 mins. To this was added 4-*tert* butyl benzoyl chloride (82 ml, 0.43mmol, 1.2 equiv.) and the resulting solution was allowed to stir at ambient temperature for 12 hours. Once complete, the solvent was removed under vacuum. The residue left was suspended in hexanes and filtered to give crude solid product which was purified by column chromatography using dichloromethane and methanol as eluents to furnish pure 4-(*tert*-butyl)-*N*-(2-(3-chlorophenyl)-2,6-dihydro-4*H*-thieno[3,4-*c*]pyrazol-3-yl)benzamide (**5a**) (52 mg, 35%) as a tan solid. $^{1}$H NMR (300 MHz, $CDCl_3$) δ 7.79 (bs, 1H), 7.74-7.71 (m, 2H), 7.58-7.57 (m, 1H), 7.52-6.50 (m, 2H), 7.44-7.37 (m, 3H), 4.10 (s, 2H), 4.05 (s, 2H), 1.36 (s, 9H). HRMS (ESI) m/z: $[M + H]^+$ Calcd for $C_{22}H_{23}ClN_3OS$ 412.1250, found 412.1251. Retention factor (TLC)= 0.5 (4:1::Hexanes:ethylacetate).

***4-(tert-butyl)-N-(2-(4-chlorophenyl)-2,6-dihydro-4H-thieno[3,4-c]pyrazol-3-yl)benzamide*** (**5b**) (PC-01-171E): Synthesized from 2-(4-chlorophenyl)-2,6-dihydro-4H-thieno[3,4-c] pyrazol-3-amine (**4b**) (90 mg, 0.35mmol, 1 equiv.) and 4-*tert* butyl benzoyl chloride (82 ml, 0.43mmol, 1.2 equiv.) following General Experimental Procedure B to provide compound **5b** as a cream solid (62 mg, 42%). $^{1}$H NMR (300 MHz,

$CDCl_3$) δ 8.08 (s, 1H), 7.70-7.67 (m, 2H), 7.49-7.46 (m, 2H), 7.44-7.35 (m, 4H), 3.975 (d, *J*= 3Hz, 4H), 1.35 (s,9H). HRMS (ESI) m/z: $[M + H]^+$ Calcd for $C_{22}H_{23}ClN_3OS$ 412.1250, found 412.1244. Retention factor (TLC)= 0.5 (4:1::Hexanes:ethylacetate).

***4-(tert-butyl)-N-(2-(3-methylphenyl)-2,6-dihydro-4H-thieno[3,4-c]pyrazol-3-yl)benzamide*** (**5c**) (PC-01-171C): Synthesized from 2-(3-methylphenyl)-2,6-dihydro-4H-thieno[3,4-c] pyrazol-3-amine (**4c**) (80 ml, 0.43mmol, 1.0 equiv.) and 4-*tert* butyl benzoyl chloride (82 ml, 0.43mmol, 1.0 equiv.) following General Experimental Procedure B to provide compound **5c** as a cream solid (67 mg, 48%). $^1$H NMR (300 MHz, $CDCl_3$) δ 8.05 (bs, 1H), 7.70-7.68 (m, 2H), 7.49-7.46 (m, 2H), 7.40-7.35 (m, 1H), 7.31 (bs, 1H), 7.25-7.22 (m, 2H), 4.10 (s, 2H), 3.99 (s, 2H), 2.41 (s, 3H), 1.34 (s, 9H). HRMS (ESI) m/z: $[M + H]^+$ Calcd for $C_{23}H_{26}N_3OS$ 392.1796, found 392.18. Retention factor (TLC)= 0.4 (5:1::Hexanes:ethylacetate).

***4-(tert-butyl)-N-(2-(4-methylphenyl)-2,6-dihydro-4H-thieno[3,4-c]pyrazol-3-yl)benzamide*** (**5d**) (PC-01-184): Synthesized from 2-(4-methylphenyl)-2,6-dihydro-4H-thieno[3,4-c] pyrazol-3-amine (**4d**) (82 mg, 0.35mmol, 1.0 equiv.) and 4-*tert* butyl benzoyl chloride (82 ml, 0.43mmol, 1.0 equiv.) following General Experimental Procedure B to provide compound **5d** as a cream solid (63 mg, 46%). $^1$H NMR (300 MHz, $CDCl_3$) δ 7.93(s, 1H), 7.78-7.62 (m, 2H), 7.56-7.41 (m, 2H), 7.37-7.29 (m, 4H), 4.19-3.95 (m, 4H), 2.44 (t, *J*=18 Hz, 3H), 1.35 (s, 9H). HRMS (ESI) m/z: $[M + H]^+$ Calcd for $C_{23}H_{26}N_3OS$ 392.1796, found 392.18. Retention factor (TLC)= 0.4 (5:1::Hexanes:ethylacetate).

*4-(tert-butyl)-N-(2-(3-trifluoromethylphenyl)-2,6-dihydro-4H-thieno[3,4-c]pyrazol-3-yl)benzamide* (**5e**) (PC-01-170B/158/171B): Synthesized from 2-(3-trifluoromethylphenyl)-2,6-dihydro-4H-thieno[3,4-c] pyrazol-3-amine (**4e**) (102 mg, 0.35mmol, 1.0 equiv.) and 4-*tert* butyl benzoyl chloride (82 ml, 0.43mmol, 1.0 equiv.) following General Experimental Procedure B to provide compound **5e** as a yellow solid (93 mg, 58%). $^1$H NMR (300 MHz, $CDCl_3$) δ 7.84 (bs, 1H), 7.76-7.64 (m, 6H), 7.52 (d, *J*= 9Hz, 2H), 4.095 (d,

*J*= 9Hz, 4H), 1.36 (t, *J*= 18 Hz, 9H). $^{19}$F NMR (282 MHz, $CDCl_3$) δ -62.64 (s). LRMS (ESI) m/z: $[M + H]^+$ Calcd for $C_{23}H_{23}F_3N_3OS$ 446.14, found 446.10. Retention factor (TLC)= 0.3 (4:1::Hexanes:ethylacetate).

***4-(tert-butyl)-N-(2-(4-trifluoromethylphenyl)-2,6-dihydro-4H-thieno[3,4-c]pyrazol-3-yl)benzamide*** (**5f**) (PC-01-171F): Synthesized from 2-(4-trifluoromethylphenyl)-2,6-dihydro-4H-thieno[3,4-c] pyrazol-3-amine (**4f**) (102 mg, 0.35mmol, 1.0 equiv.) and 4-*tert* butyl benzoyl chloride (82 ml, 0.43mmol, 1.0 equiv.) following General Experimental Procedure B to provide compound **5f** as a yellow solid (86 mg, 54%). $^1$H NMR (300 MHz, $CDCl_3$) δ 7.84 (bs, 1H), 7.76-7.64 (m, 6H), 7.52 (d, *J*= 9Hz, 2H), 4.05 (bs, 4H), 1.36 (s, 9H). $^{19}$F NMR (282 MHz, $CDCl_3$) δ -62.43 (s). HRMS (ESI) m/z: $[M + H]^+$ Calcd for $C_{23}H_{23}F_3N_3OS$ 446.1514, found 446.1509. Retention factor (TLC)= 0.3 (4:1::Hexanes:ethylacetate).

**General Experimental Procedure C**: **4-(*tert*-butyl)-*N*-(2-(3-chlorophenyl)-2,6-dihydro-4*H*-thieno[3,4-*c*]pyrazol-3-yl) benzene sulfonamide** (**6a**) (PC-01-164): To an oven dried round bottom flask was added 2-(3-chlorophenyl)-2,6-dihydro-4H-thieno[3,4-c] pyrazol-3-amine (**4a**) (100 mg, 0.32 mmol, 1 equiv.) followed by anhydrous DMF and the resulting solution was allowed to stir at 0 °C for 10 minutes. To this was added sodium hydride (60% suspension in mineral oil, 0.64mmol, 2 equiv.) in portions. The solution was allowed to stir at same temperature until effervescence ceased after which was added 4-tert butyl benzenesulfonyl chloride (94 mg, 0.40mmol, 1 equiv.). The reaction was stirred at 0 °C for 15 mins and then at rt for additional 60 minutes. The reaction was monitored by TLC and once complete, DMF was removed under vacuum and the residue dissolved in ethyl acetate which was washed with a saturated solution of ammonium chloride, followed by water and brine. The ethyl acetate fractions were collected, concentrated and purified by column chromatography by running a gradient from hexanes to ethyl acetate. The fractions containing the product were concentrated to give **6a** (96 mg, 54%) as cream solid. $^1$H NMR (300 MHz, $CDCl_3$) δ 7.54-7.40 (m, 4.5H), 7.26-7.14 (m, 2H), 7.00-6.97 (m, 1.5H), 3.95 (t, *J* = 18 Hz, 2H), 3.64 (t, *J* = 18 Hz, 2H), 1.33 (t, *J*= 18 Hz, 9H). HRMS (ESI) m/z: $[M + H]^+$ Calcd for $C_{21}H_{23}ClN_3O_2S_2$ 448.0920, found 448.0915. Retention factor (TLC)= 0.5 (4:1::Hexanes:ethylacetate).

***4-(tert-butyl)-N-(2-(4-chlorophenyl)-2,6-dihydro-4H-thieno[3,4-c]pyrazol-3-yl) benzene sulfonamide*** (**6b**) (PC-01-180A): Synthesized from 2-(4-chlorophenyl)-2,6-dihydro-4H-thieno[3,4-c] pyrazol-3-amine (**4b**) (102 mg, 0.35mmol, 1.0 equiv.) and 4-tert butyl benzenesulfonyl chloride (94 mg, 0.40mmol, 1 equiv.) following General Experimental Procedure C to provide compound **6b** as a cream solid (93 mg, 52%). $^1$H NMR (300 MHz, $CDCl_3$) δ 7.45 (dd, *J*=24, 6 Hz, 4H), 7.25-7.21 (m, 2H), 7.14 (s, 1H), 7.07-6.92 (m, 2H), 3.98 (t, *J* = 18 Hz, 2H), 3.65 (t, *J* = 18 Hz, 2H), 1.36 (t, *J*= 18 Hz, 9H). HRMS (ESI) m/z: $[M + H]^+$ Calcd for $C_{21}H_{23}ClN_3O_2S_2$ 448.9208, found 448.0945. Retention factor (TLC)=0.50(4:1::Hexanes:ethylacetate).

***4-(tert-butyl)-N-(2-(3-methylphenyl)-2,6-dihydro-4H-thieno[3,4-c]pyrazol-3-yl) benzene sulfonamide*** (**6c**) (PC-01-178B): Synthesized from 2-(3-methylphenyl)-2,6-dihydro-4H-thieno[3,4-c] pyrazol-3-amine (**4c**) (60 mg, 0.21mmol, 1 equiv.) and 4-tert butyl benzenesulfonyl chloride (51 mg, 0.21mmol, 1 equiv.) following General Experimental Procedure C to provide compound **6c** as a tan solid (53 mg, 58%). $^1$H NMR (300 MHz, $CDCl_3$) δ 7.58-7.36 (m, 4H), 7.14-7.12 (m, 2H), 6.99 (s, 1H), 6.75 (bs, 1H) 6.69-6.66 (m, 1H), 4.04-3.91 (3, 2H), 3.87-3.74 (m, 2H), 2.28 (t, *J*= 21Hz, 3H), 1.35 (t, *J*= 18 Hz, 9H). HRMS (ESI) m/z: $[M + H]^+$ Calcd for $C_{22}H_{26}N_3O_2S_2$ 428.1466, found 428.1469. Retention factor (TLC)= 0.4 (5:1::Hexanes:ethylacetate).

***4-(tert-butyl)-N-(2-(4-methylphenyl)-2,6-dihydro-4H-thieno[3,4-c]pyrazol-3-yl) benzene sulfonamide*** (**6d**) (PC-01-180C): Synthesized from 2-(4-methylphenyl)-2,6-dihydro-4H-thieno[3,4-c] pyrazol-3-amine (**4d**) (60 mg, 0.21mmol, 1 equiv.) and 4-tert butyl benzenesulfonyl chloride (51 mg, 0.21mmol, 1 equiv.) following General Experimental Procedure C to provide compound **6d** as a beige solid (50 mg, 56%). $^1$H NMR (300 MHz, $CDCl_3$) δ 7.54-7.34 (m, 4H), 7.21 (bs, 1H), 7.14 (s, 1H), 7.05-6.75 (m, 4H), 3.95 (t, *J* = 18 Hz, 2H), 3.72 (t, *J* = 18 Hz, 2H), 2.35 (t, *J*=18Hz, 3H), 1.35 (t, *J*= 18 Hz, 9H). HRMS (ESI) m/z: $[M + H]^+$ Calcd for $C_{22}H_{26}N_3O_2S_2$ 428.1466, found 428.1467. Retention factor (TLC)= 0.4 (5:1::Hexanes:ethylacetate).

***4-(tert-butyl)-N-(2-(3-trifluoromethylphenyl)-2,6-dihydro-4H-thieno[3,4-c]pyrazol-3-yl) benzene sulfonamide*** (**6e**) (PC-01-178A): Synthesized from 2-(3-trifluoromethylphenyl)-2,6-dihydro-4H-thieno[3,4-c] pyrazol-3-amine (**4e**) (80 mg, 0.34mmol, 1 equiv.) and 4-tert butyl benzenesulfonyl chloride (81 mg, 0.34mmol, 1 equiv.) following General Experimental Procedure C to provide compound **6e** as a beige solid (97 mg, 58%). $^1$H NMR (300 MHz, $CDCl_3$) δ 7.57-7.33 (m, 9H), 3.96 (t, *J* = 18 Hz, 2H), 3.57 (t, *J* = 18 Hz, 2H), 1.32 (t, *J*= 18 Hz, 9H). $^{19}$F NMR (282 MHz, $CDCl_3$) δ -62.30- -62.59 (m). HRMS (ESI) m/z: $[M + H]^+$ Calcd for $C_{22}H_{23}F_3N_3O_2S_2$ 482.1184, found 482.1188. Retention factor (TLC)= 0.3 (4:1::Hexanes:ethylacetate).

***4-(tert-butyl)-N-(2-(4-trifluoromethylphenyl)-2,6-dihydro-4H-thieno[3,4-c]pyrazol-3-yl) benzene sulfonamide*** (**6f**) (PC-01-180B): Synthesized from 2-(4-trifluoromethylphenyl)-2,6-dihydro-4H-thieno[3,4-c] pyrazol-3-amine (**4f**) (80 mg, 0.34mmol, 1 equiv.) and 4-tert butyl benzenesulfonyl chloride (81 mg, 0.34mmol, 1 equiv.) following General Experimental Procedure C to provide compound **6f** as a beige solid (87 mg, 52%). $^1$H NMR (300 MHz, $CDCl_3$) δ 7.55-7.47 (m, 4H), 7.41-7.37 (m, 2H), 7.90 (s, 1H), 7.28 (s, 1H), 3.98 (t, *J* = 18 Hz, 2H), 3.62 (t, *J* = 18 Hz, 2H), 1.33 (t, *J*= 18 Hz, 9H). $^{19}$F NMR (282 MHz, $CDCl_3$) δ -62.35- -62.49 (m). HRMS (ESI) m/z: $[M + H]^+$ Calcd for $C_{22}H_{23}F_3N_3O_2S_2$ 482.1184, found 482.1184. Retention factor (TLC)= 0.3 (4:1::Hexanes:ethylacetate).

***1-(4-(tert-butyl)phenyl)-3-(2-(3-chlorophenyl)-2,6-dihydro-4H-thieno[3,4-c]pyrazol-3-yl)urea*** (**7a**) (PC-01-186): Synthesized from 2-(3-chlorophenyl)-2,6-dihydro-4H-thieno[3,4-c] pyrazol-3-amine (**4a**) (50 mg, 0.2 mmol, 1.0 equiv.) and 4-tert butyl phenylisocyanate (36 ml, 0.2mmol, 1.0 equiv.) following General Experimental Procedure C to provide compound **7a** as a cream solid (38 mg, 45%). $^1$H NMR (300 MHz, DMSO-$d_6$) δ 9.12 (t, *J*= 18 Hz, 1H), 8.69 (bs, 1H), 7.63 (s, 1H), 7.57—7.54 (m, 2H), 7.50-7.45 (m, 1H), 7.33-7.23 (m, 4H), 3.92 (d, *J*= 27 Hz, 4H), 1.24 (s, 9H). HRMS (ESI) m/z: $[M + H]^+$ Calcd for $C_{22}H_{24}ClN_4OS$ 427.1359, found 427.1360. Retention factor (TLC)= 0.2 (4:1::Hexanes:ethylacetate).

***1-(4-(tert-butyl)phenyl)-3-(2-(4-chlorophenyl)-2,6-dihydro-4H-thieno[3,4-c]pyrazol-3-yl)urea*** (**7b**) (PC-01-188): Synthesized from 2-(4-chlorophenyl)-2,6-dihydro-4H-thieno[3,4-c] pyrazol-3-amine (**4b**) (50 mg, 0.2 mmol, 1.0 equiv.) and 4-tert butyl phenylisocyanate (36 ml, 0.2mmol, 1.0 equiv.) following General Experimental Procedure C to provide compound **7b** as a beige solid (36 mg, 42%). $^{1}$H NMR (300 MHz, DMSO-$d_6$) δ 8.97 (s, 1H), 8.54 (bs, 1H), 7.61-7.52 (m, 4H), 7.33—7.22 (m, 4H), 3.96 (s, 2H), 3.88 (s, 2H), 1.24 (s, 9H). HRMS (ESI) m/z: $[M + H]^+$ Calcd for $C_{22}H_{24}ClN_4OS$ 427.1359, found 427.1348. Retention factor (TLC)= 0.2 (4:1::Hexanes:ethylacetate).

***1-(4-(tert-butyl)phenyl)-3-(2-(3-methylphenyl)-2,6-dihydro-4H-thieno[3,4-c]pyrazol-3-yl)urea*** (**7c**) (PC-01-190): Synthesized from 2-(3-methylphenyl)-2,6-dihydro-4H-thieno[3,4-c] pyrazol-3-amine (**4c**) (50 mg, 0.2 mmol, 1.0 equiv.) and 4-tert butyl phenylisocyanate (39 ml, 0.2 mmol, 1.0 equiv.) following General Experimental Procedure C to provide compound **7c** as a beige solid (49 mg, 54%). $^{1}$H NMR (300 MHz, DMSO-$d_6$) δ 8.87 (s, 1H), 8.35 (s, 1H), 7.44-7.22 (m, 8H), 3.925 (d, *J*= 15Hz, 4H), 2.37 (s, 3H), 1.24 (s, 9H). HRMS (ESI) m/z: $[M + H]^+$ Calcd for $C_{23}H_{27}N_4OS$ 407.1905, found 407.1903. Retention factor (TLC)= 0.3 (3:1::Hexanes:ethylacetate).

***1-(4-(tert-butyl)phenyl)-3-(2-(4-methylphenyl)-2,6-dihydro-4H-thieno[3,4-c]pyrazol-3-yl)urea*** (**7d**) (PC-01-192): Synthesized from 2-(4-methylphenyl)-2,6-dihydro-4H-thieno[3,4-c] pyrazol-3-amine (**4d**) (50 mg, 0.2 mmol, 1.0 equiv.) and 4-tert butyl phenylisocyanate (39 ml, 0.2 mmol, 1.0 equiv.) following General Experimental Procedure C to provide compound **7d** as a tab solid (42 mg, 48%). $^{1}$H NMR (300 MHz, DMSO-$d_6$) δ 8.85 (s, 1H), 8.32 (s, 1H), 7.40—7.22 (m, 8H), 3.92 (d, *J*= 15 Hz, 4H), 1.24 (s, 9H). HRMS (ESI) m/z: $[M + H]^+$ Calcd for $C_{23}H_{27}N_4OS$ 407.1905, found 407.1910. Retention factor (TLC)= 0.3 (3:1::Hexanes:ethylacetate).

***1-(4-(tert-butyl)phenyl)-3-(2-(3-trifluoromethylphenyl)-2,6-dihydro-4H-thieno[3,4-c]pyrazol-3-yl)urea*** (**7e**) (PC-01-194): Synthesized from 2-(3-trifluoromethylphenyl)-2,6-dihydro-4H-thieno[3,4-c] pyrazol-3-

amine (**4e**) (50 mg, 0.17 mmol, 1.0 equiv.) and 4-tert butyl phenylisocyanate (32 ml, 0.2 mmol, 1.0 equiv.) following General Experimental Procedure C to provide compound **7e** as a tab solid (37 mg, 46%). $^1$H NMR (300 MHz, DMSO-$d_6$) δ 8.93 (s, 1H), 8.53 (s, 1H), 7.88 (bs, 2H), 7.78-7.76 (m, 2H), 7.28 (s, 4H), 3.94 (d, *J*= 30 Hz, 4H), 1.24 (s, 9H). $^{19}$F NMR (282 MHz, DMSO-$d_6$) δ -61.17 (s). HRMS (ESI) m/z: [M + H]$^+$ Calcd for $C_{23}H_{24}F_3N_4OS$ 461.1623, found 461.1622. Retention factor (TLC)= 0.3 (2:1::Hexanes:ethylacetate).

***1-(4-(tert-butyl)phenyl)-3-(2-(4-trifluoromethylphenyl)-2,6-dihydro-4H-thieno[3,4-c]pyrazol-3-yl)urea*** (**7f**) (PC-01-196): Synthesized from 2-(4-trifluoromethylphenyl)-2,6-dihydro-4H-thieno[3,4-c] pyrazol-3-amine (**4f**) (50 mg, 0.17 mmol, 1.0 equiv.) and 4-tert butyl phenylisocyanate (32 ml, 0.2 mmol, 1.0 equiv.) following General Experimental Procedure C to provide compound **7f** as a tab solid (34 mg, 42%). $^1$H NMR (300 MHz, DMSO-$d_6$) δ 8.95 (s, 1H), 8.62 (bs, 1H), 7.92-7.78 (m, 4H), 7.33-7.23 (m, 4H), 3.94 (d, *J*= 30 Hz, 4H), 1.24 (s, 9H). $^{19}$F NMR (282 MHz, DMSO-$d_6$) δ -60.76 (s). HRMS (ESI) m/z: [M + H]$^+$ Calcd for $C_{23}H_{24}F_3N_4OS$ 461.1623, found 461.1630. Retention factor (TLC)= 0.3 (2:1::Hexanes:ethylacetate).

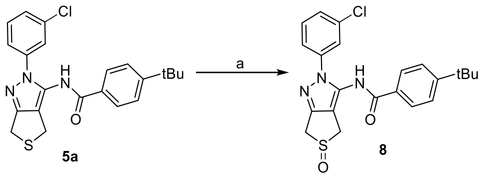


***4-(tert-butyl)-N-(2-(3-chlorophenyl)-5-oxido-2,6-dihydro-4H-thieno[3,4-c]pyrazol-3-yl)benzamide*** (**8**) (PC-02-06): To a solution of **5a** (20 mg, 0.04 mmol, 1 equiv.) in dichloromethane was added silica gel (1.5 equiv.) and acetic anhydride (6 ml, 0.05 mmol, 1.1 equiv.) and the resulting solution was allowed to stir at rt for 10 minutes. To this was added hydrogen peroxide (2 ml, 0.05 mmol, 1.2 equiv.) and the resulting solution was stirred at room temperature for 24 hours. The reaction was then concentrated and purified by column chromatography to give **8** as a white solid in quantitative yield (20 mg). $^1$H NMR (300 MHz, $CDCl_3$) δ 9.25 (s, 1H), 7.83 (d, *J* = 8.3 Hz, 2H), 7.66 (s, 1H), 7.46 (m, 4H), 4.43 (d, *J* = 16.3 Hz, 1H), 4.10 (d, *J* =

16.1 Hz, 1H), 3.62 (dd, $J$ = 36.6, 16.2 Hz, 2H), 1.36 (s, 9H). HRMS (ESI) m/z: $[M + H]^+$ Calcd for $C_{22}H_{23}ClN_3O_2S$ 428.1199, found 428.1179. Retention factor (TLC)= 0.2 (1:1::Hexanes:ethylacetate).

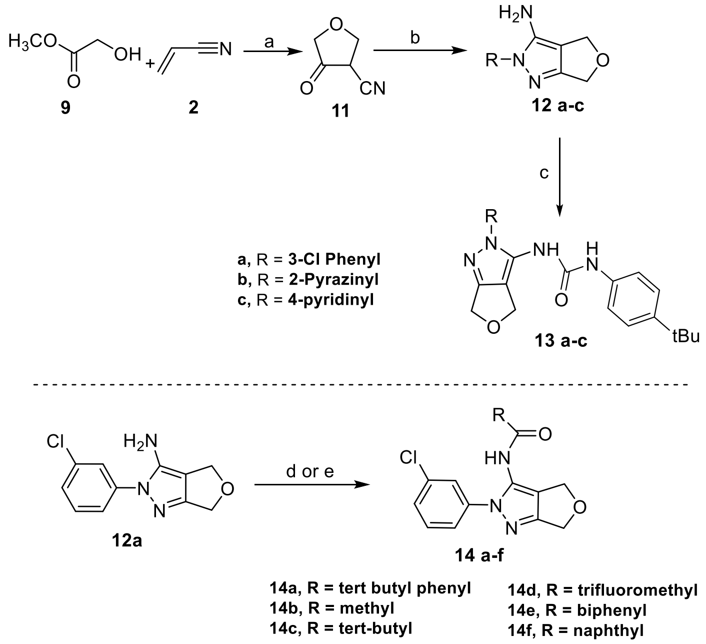


***4-oxotetrahydrofuran-3-carbonitrile*** (**11**) (PC-01-212): To dry THF at 0 °C was added NaH (266 mg, 11 mmol, 2eq.) and stirred at 0 °C for 10 minutes. To this was added a solution of methyl 2-hydroxyacetate (429 ml, 5.5 mmol, 1 equiv.) in dry THF and the solution was stirred at 0 °C for 10 minutes. To this was added acrylonitrile (366 ml, 5.5 mmol, 1 equiv.) and the reaction was refluxed for 2 hours. Monitor the reaction on TLC, once complete, the reaction was quenched with sat. solution of $NaHCO_3$ and extracted with ethyl ether. The aqueous layer was acidified to pH 1 and then extracted with dichloromethane. The organic fractions were concentrated to give crude **31** as yellow oil. (289 mg, 48% yield). Retention factor (TLC)= 0.3 (1:1::Hexanes:ethylacetate). Crude **11** was used for further reactions without further purification.

***2-(3-chlorophenyl)-2,6-dihydro-4H-furo[3,4-c]pyrazol-3-amine*** (**12a**) (PC-01-214): To a solution of 4-oxotetrahydrofuran-3-carbonitrile (**10**) (90 mg, 0.81 mmol, 1equiv.) in ethanol, was added 3-chlorophenyl hydrazine (190 mg, 0.81 mmol, 1equiv.) and the resulting solution was refluxed for 3 hours. Once the reaction was complete, ethanol was removed under vacuum and the residue was dissolved in ethyl acetate which was washed with 5% NaOH solution. The organic layers were collected, concentrated and the resulting crude product was purified by column chromatography eluting in 2: 1:: hexanes:ethylacetate, yielding pure **12** (111 mg, 58%). $^1$H NMR (300 MHz, $CDCl_3$) δ 7.46 (m, 4H), 4.83 (m, 4H), 3.84 (bs, 2H). HRMS (ESI) m/z: $[M + H]^+$ Calcd for $C_{11}H_{11}ClN_3O$ 236.0590, found 236.0547. Retention factor (TLC)=0.2 (1:1::Hexanes:ethylacetate).

***2-(pyrazin-2-yl)-4,6-dihydro-2H-furo[3,4-c]pyrazol-3-amine*** **(12b) (PC-02-50):** To a solution of 4-oxotetrahydrofuran-3-carbonitrile (**10**) (90 mg, 0.81 mmol, 1 equiv.) in ethanol, was added 4-hydrazinylpyrimidine (90 mg, 0.81 mmol, 1 equiv.) and the resulting solution was refluxed for 2 hours. Once the reaction was complete, ethanol was removed under vacuum and the residue was dissolved in ethyl acetate which was washed with 5% NaOH solution. The organic layers were collected, concentrated and the resulting crude product was purified by column chromatography eluting in 1:2:: hexanes:ethylacetate, to furnish **12b** as a beige solid (111 mg, 64%). $^1$H NMR (300 MHz, $CDCl_3$) δ 9.28 (d, *J* = 1.5 Hz, 1H), 8.39 (d, *J* = 2.7 Hz, 1H), 8.26 (dd, *J* = 2.7, 1.5 Hz, 1H), 5.80 (s, 2H), 4.83 (s, 4H). HRMS (ESI) m/z: $[M + H]^+$ Calcd for $C_9H_{10}N_5O$ 204.0885, found: 204.0874. Retention factor (TLC)= 0.2 (1:1::Hexanes:ethylacetate).

***2-(pyridin-4-yl)-4,6-dihydro-2H-furo[3,4-c]pyrazol-3-amine*** (**12c**) **(PC-02-51):** To a solution of 4-oxotetrahydrofuran-3-carbonitrile (**10**) (90 mg, 0.81 mmol, 1 equiv.) in ethanol, was added 4-hydrazineylpyridine (87 mg, 0.81 mmol, 1 equiv.) and the resulting solution was refluxed for 2 hours. Once the reaction was complete, ethanol was removed under vacuum and the residue was dissolved in ethyl acetate which was washed with 5% NaOH solution. The organic layers were collected, concentrated and the resulting crude product was purified by column chromatography using dichloromethane in methanol as eluting solvent to furnish **12c** as a tan solid (111 mg, 72%). $^1$H NMR (300 MHz, $CDCl_3$) δ 8.68 (dd, *J* = 4.6, 1.6 Hz, 2H), 7.64 (dd, *J* = 4.6, 1.6 Hz, 2H), 4.83 (s, 4H), 3.99 (s, 2H). HRMS (ESI) m/z: $[M + H]^+$ Calcd for $C_{10}H_{11}N_4O$ 203.0933, found: 203.0940. Retention factor (TLC)= 0.3 (10:1:: Dichloromethane: Methanol).

***1-(4-(tert-butyl)phenyl)-3-(2-(3-chlorophenyl)-2,6-dihydro-4H-furo[3,4-c]pyrazol-3-yl)urea*** (**13a**) (PC-III-149)**:** Synthesized from 2-(3-chlorophenyl)-2,6-dihydro-4H-furo[3,4-c]pyrazol-3-amine (**12a**) (50 mg, 0.2 mmol, 1.0 equiv.) and 4-tert butyl phenylisocyanate (36 ml, 0.2mmol, 1.0 equiv.) following General Experimental Procedure C to provide compound **13a** as a cream solid (51 mg, 58%). $^{1}$H NMR (300 MHz, $CDCl_3$) δ 7.43 (s, 1H), 7.37 – 7.29 (m, 5H), 7.16 (d, *J* = 8.6 Hz, 2H), 5.06 (s, 2H), 4.84 (s, 2H), 1.30 (s, 10H). HRMS (ESI) m/z: $[M + H]^+$ Calcd for $C_{22}H_{24}ClN_4O_2$ 411.1588, found 411.1590.

***1-(4-(tert-butyl)phenyl)-3-(2-(pyrazin-2-yl)-4,6-dihydro-2H-furo[3,4-c]pyrazol-3-yl)urea*** (**13b**) (PC-02-53A)**:** Synthesized from 2-(pyrazin-2-yl)-4,6-dihydro-2H-furo[3,4-c]pyrazol-3-amine **(12b)** (50 mg, 0.24 mmol, 1.0 equiv.) and 4-tert butyl phenylisocyanate (43 ml, 0.24 mmol, 1.0 equiv.) following General Experimental Procedure C to provide compound **13b** as a cream solid (63 mg, 68%). $^{1}$H NMR (300 MHz, $CDCl_3$) δ 10.97 (s, 1H), 9.30 (d, *J* = 1.4 Hz, 1H), 8.34 (d, *J* = 2.7 Hz, 1H), 7.73 (s, 1H), 7.49 (d, *J* = 8.6 Hz, 2H), 7.30 (m, 2H), 6.83 (s, 1H), 5.19 (s, 2H), 4.89 (s, 2H), 1.40 (s, 9H). HRMS (ESI) m/z: $[M + H]^+$ Calcd for $C_{20}H_{23}N_6O_2$ 379.1882, found 379.1911.

***1-(4-(tert-butyl)phenyl)-3-(2-(pyridin-4-yl)-4,6-dihydro-2H-furo[3,4-c]pyrazol-3-yl)urea*** (**13c**) **(PC-02-64):** Synthesized from 2-(pyridin-4-yl)-4,6-dihydro-2H-furo[3,4-c]pyrazol-3-amine (**12c**) (50 mg, 0.24 mmol, 1.0 equiv.) and 4-tert butyl phenylisocyanate (43 ml, 0.24 mmol, 1.0 equiv.) following General Experimental Procedure C to provide compound **13c** as a tan solid (79 mg, 83%). $^{1}$H NMR. HRMS (ESI) m/z: $[M + H]^+$ Calcd for $C_{21}H_{24}N_5O_2$ 378.1930, found 378.1946. Retention factor (TLC) = 0.5 (10:1:: Dichloromethane: Methanol).

***4-(tert-butyl)-N-(2-(3-chlorophenyl)-2,6-dihydro-4H-furo[3,4-c]pyrazol-3-yl)benzamide*** (**14a**) (PC-01-216): Synthesized from 2-(3-chlorophenyl)-2,6-dihydro-4H-furo[3,4-c]pyrazol-3-amine (**12a**) (40 mg, 0.17mmol, 1 equiv.) and 4-*tert* butyl benzoyl chloride (67 ml, 0.34 mmol, 2.0 equiv.) following General

Experimental Procedure B to provide compound **14a** as a yellow solid (42 mg, 62%). $^1$H NMR (300 MHz, $CDCl_3$) δ 8.09 (s, 1H), 7.71-7.67 (m, 2H), 7.61-7.60 (m, 2H), 7.53-7.43 (m, 5H) 5.18 (s, 2H) 4.91 (s, 2H), 1.35 (s, 9H). HRMS (ESI) m/z: $[M + H]^+$ Calcd for $C_{22}H_{23}ClN_3O_2$ 396.1479, found 396.1471. Retention factor (TLC)= 0.3 (1:1::Hexanes:ethylacetate).

***N-(2-(3-chlorophenyl)-2,6-dihydro-4H-furo[3,4-c]pyrazol-3-yl)acetamide*** (**14b**) (PC-02-33): Synthesized from 2-(3-chlorophenyl)-2,6-dihydro-4H-furo[3,4-c]pyrazol-3-amine (**12a**) (80 mg, 0.34 mmol, 1 equiv.) and acetyl chloride (0.43 mmol, 1.2 equiv.) following General Experimental Procedure B to provide compound **14b** as a white solid (45 mg, 48%). $^1$H NMR (300 MHz, $CDCl_3$) δ 7.51-7.41 (m, 4H), 7.37-7.34 (m, 1H), 5.04 (s, 2H) 4.86 (s, 2H), 2.15 (s, 3H). HRMS (ESI) m/z: $[M + H]^+$ Calcd for $C_{13}H_{13}ClN_3O_2$ 278.0696, found 278.0691. Retention factor (TLC)= 0.4 (1:1::Hexanes:ethylacetate).

***N-(2-(3-chlorophenyl)-2,6-dihydro-4H-furo[3,4-c]pyrazol-3-yl)pivalamide*** (**14c**) (PC-02-36): Synthesized from 2-(3-chlorophenyl)-2,6-dihydro-4H-furo[3,4-c]pyrazol-3-amine (**12a**) (80 mg, 0.34 mmol, 1 equiv.) and pivaloyl chloride (0.43 mmol, 1.2 equiv.) following General Experimental Procedure B to provide compound **14c** as a white solid (63 mg, 58%). $^1$H NMR (300 MHz, $CDCl_3$) δ 7.58 (bs, 1H), 7.52-7.51 (m, 1H), 7.49-7.47 (m, 1H), 7.45-7.41 (m, 1H), 7.39-7.35 (m, 1H), 5.10 (s, 2H) 4.88 (s, 2H), 1.26 (s, 9H). HRMS (ESI) m/z: $[M + H]^+$ Calcd for $C_{16}H_{19}ClN_3O_2$ 320.1166, found 320.1169. Retention factor (TLC)= 0.5 (2:1::Hexanes:ethylacetate).

***N-(2-(3-chlorophenyl)-4,6-dihydro-2H-furo[3,4-c]pyrazol-3-yl)-2,2,2-trifluoroacetamide*** (**14d**) (PC-02-79): Synthesized from 2-(3-chlorophenyl)-2,6-dihydro-4H-furo[3,4-c]pyrazol-3-amine (**12a**) (80 mg, 0.34 mmol, 1 equiv.) and trifluoro-acetic anhydride (0.43 mmol, 1.2 equiv.) following General Experimental Procedure B to provide compound **14d** as a cream solid (83 mg, 78%). $^1$H NMR (300 MHz, $CDCl_3$) δ 8.40 (bs, 1H), 7.48 (m, 3H), 7.30 (m, 1H), 5.05 (s, 2H), 4.85 (s, 2H). $^{19}$F NMR (282 MHz, $CDCl_3$) δ -75.26 (s).

HRMS (ESI) m/z: $[M + H]^+$ Calcd for $C_{13}H_{10}ClF_3N_3O_2$ 332.0413, found: 332.0406. Retention factor (TLC)= 0.4 (3:1::Hexanes:ethylacetate).

***N-(2-(3-chlorophenyl)-2,6-dihydro-4H-furo[3,4-c]pyrazol-3-yl)-[1,1'-biphenyl]-4-carboxamide*** (**14e**) (PC-02-43): To a solution of 2-(3-chlorophenyl)-2,6-dihydro-4H-furo[3,4-c]pyrazol-3-amine (**12a**) (50 mg, 0.21 mmol, 1 equiv.) in acetonitrile was added N-methyl imidazole (2 mg, 0.02 mmol, 0.1 equiv.), tetramethylethylenediamine (3 mg, 0.02 mmol, 0.1 equiv.) and potassium carbonate (44 mg, 0.32 mmol, 1.5 equiv.) and the resulting solution was stirred at 0 °C for 10 minutes. To this solution was added [1,1'-biphenyl]-4-carbonyl chloride (51 mg, 0.23 mmol, 1.1 eq) and the resulting solution was stirred at 0 °C for 1 hour. Once complete, the reaction was quenched with water and extracted with ethyl acetate. The organic fractions were concentrated, and the product purified by column chromatography furnishing **14e** as white solid (41 mg, 46%). $^1H$ NMR (300 MHz, $CDCl_3$) δ 8.10 (s, 1H), 7.85-7.62 (m, 7H), 7.52-7.45 (m, 5H), 5.21 (s, 2H), 4.93 (s, 2H). HRMS (ESI) m/z: $[M + H]^+$ Calcd for $C_{24}H_{19}ClN_3O_2$ 416.1166, found 416.1156. Retention factor (TLC)= 0.6 (1:1::Hexanes:ethylacetate).

(**14f**) (PC-02-45): To a solution of 2-(3-chlorophenyl)-2,6-dihydro-4H-furo[3,4-c]pyrazol-3-amine (**12a**) (50 mg, 0.21 mmol, 1 equiv.) in acetonitrile was added N-methyl imidazole (2 mg, 0.02 mmol, 0.1 equiv.), tetramethylethylenediamine (3 mg, 0.02 mmol, 0.1 equiv.) and potassium carbonate (44 mg, 0.32 mmol, 1.5 equiv.) and the resulting solution was stirred at 0 °C for 10 minutes. To this solution was added 2-naphthoyl chloride (45 mg, 0.23 mmol, 1.1 eq) and the resulting solution was stirred at 0 °C for 1 hr. Once complete, the reaction was quenched with water and extracted with ethyl acetate. The organic fractions were concentrated, and the product purified by column chromatography furnishing **14f** as a beige solid (43 mg, 52%). $^1H$ NMR (300 MHz, $CDCl_3$) δ 8.30 (s, 1H),8.29 (s, 1H), 7.95-7.89 (m, 3H), 7.78-7.75 (m, 1H), 7.66-7.55 (m, 3H), 7.53-7.43 (m, 3H), 5.17 (s, 2H), 4.89 (s, 2H). HRMS (ESI) m/z: $[M + H]^+$ Calcd for $C_{22}H_{17}ClN_3O_2$ 390.1009, found 390.1009. Retention factor (TLC)= 0.6 (1:1::Hexanes:ethylacetate).

**[1]H NMR Spectra for all intermediates and final compounds:**

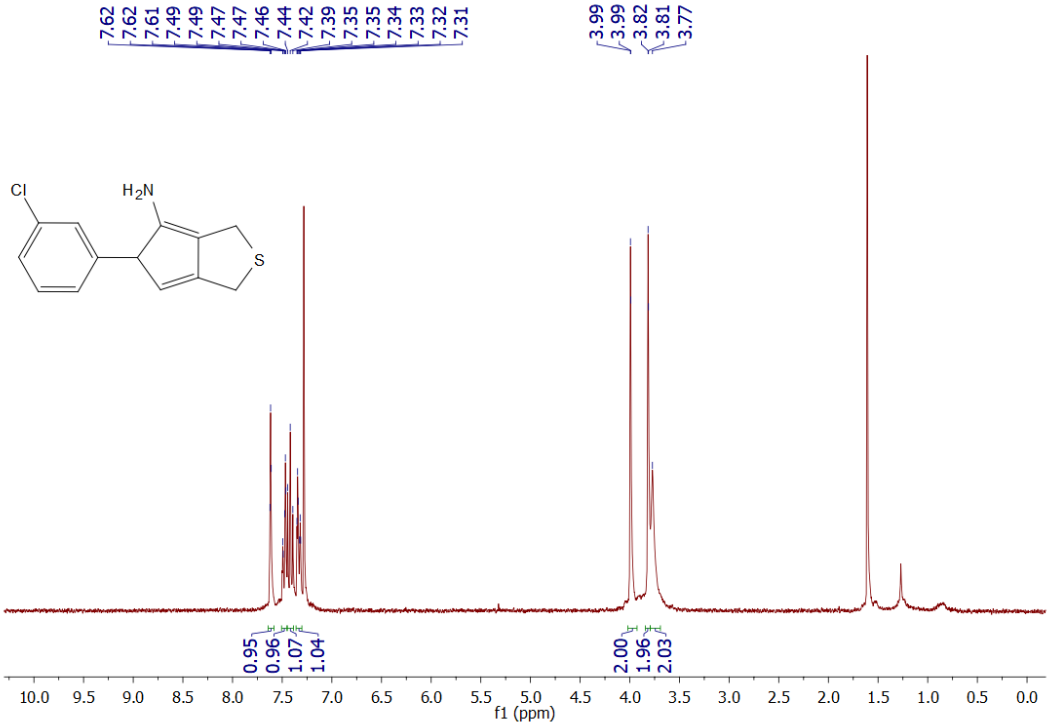


**Supplemental Figure S1**: [1]H NMR Spectrum of **4a**

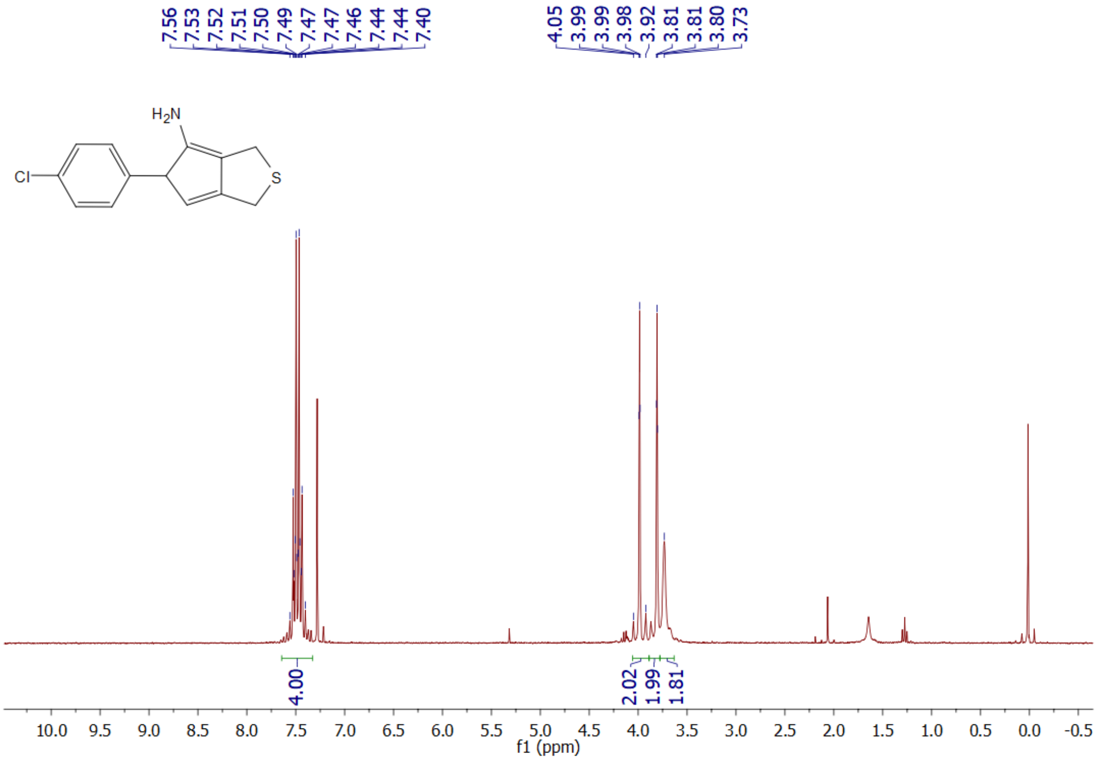


**Supplemental Figure S2**: [1]H NMR Spectrum of **4b**

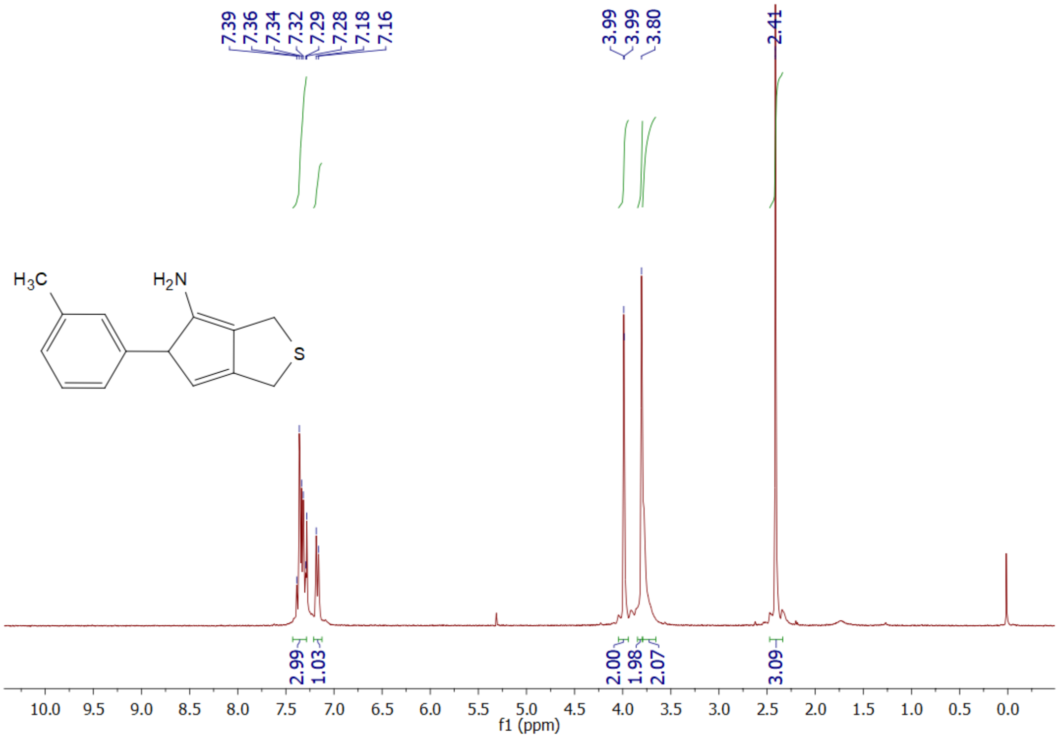


**Supplemental Figure S3**: $^{1}$H NMR Spectrum of **4c**

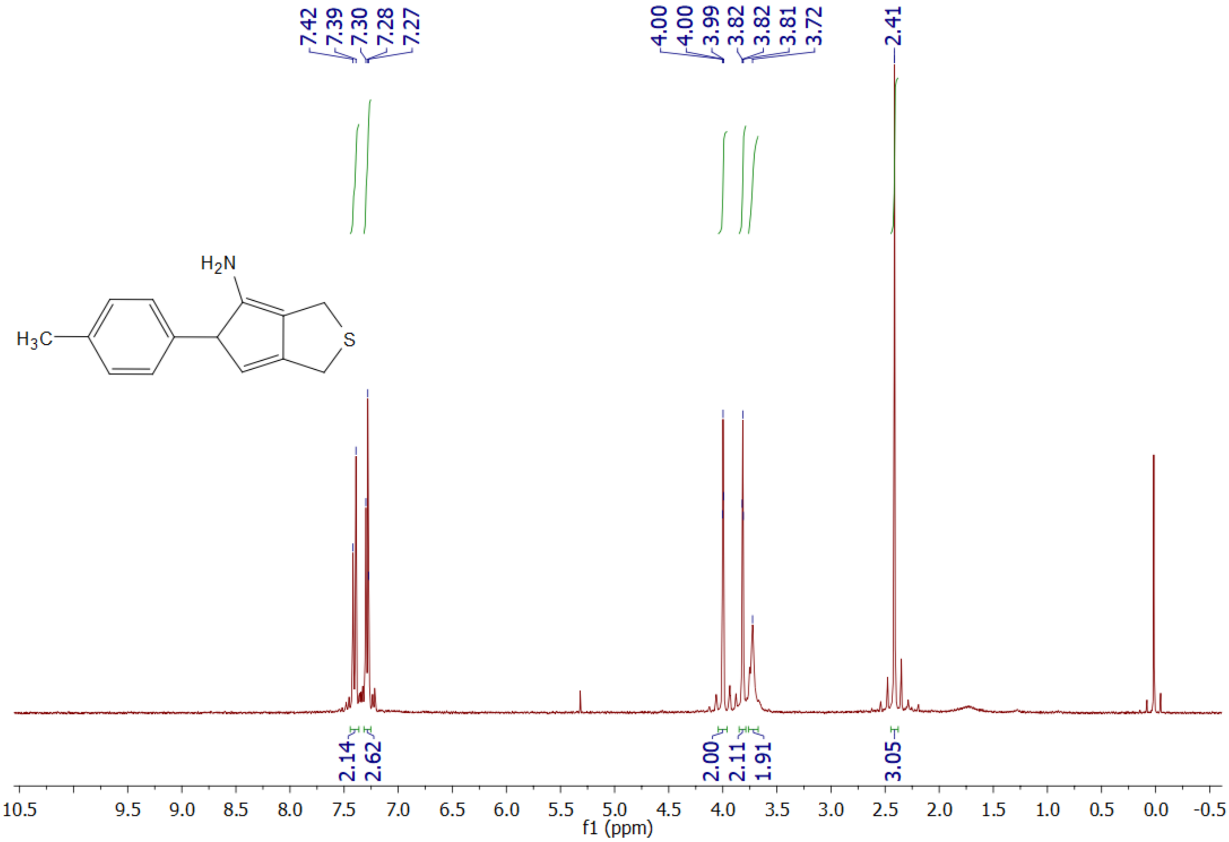


**Supplemental Figure S4**: $^{1}$H NMR Spectrum of **4d**

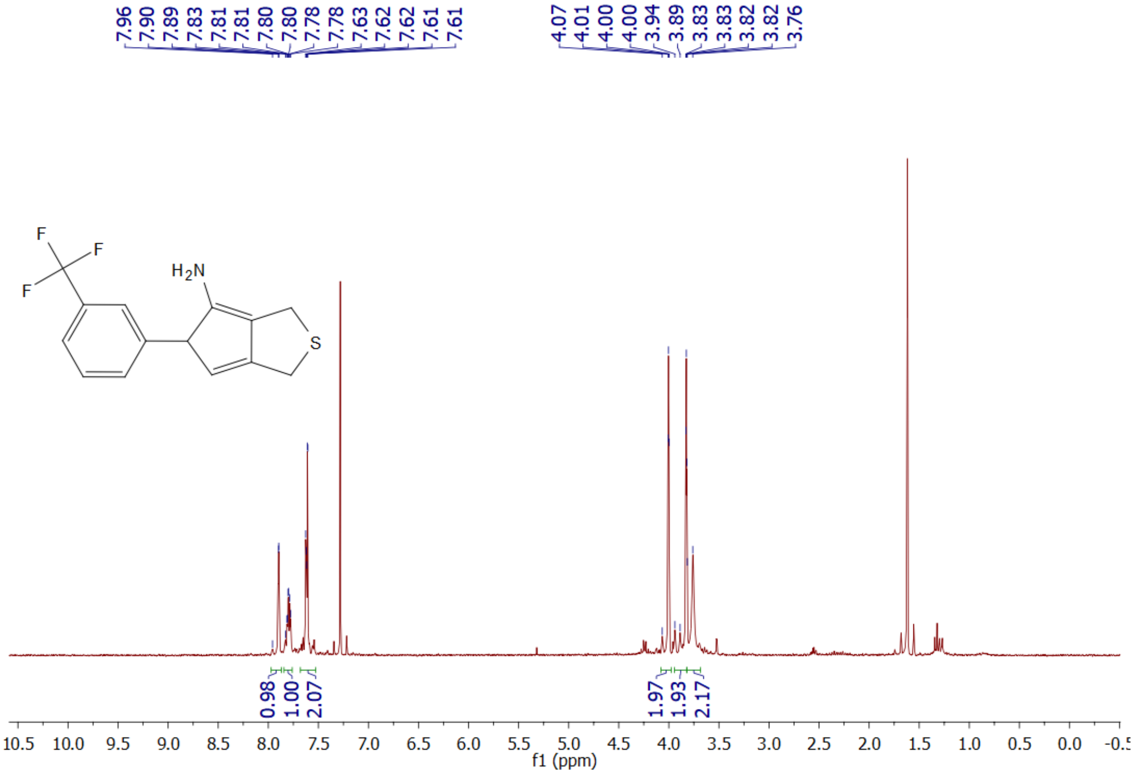


**Supplemental Figure S5**: $^{1}$H NMR Spectrum of **4e**

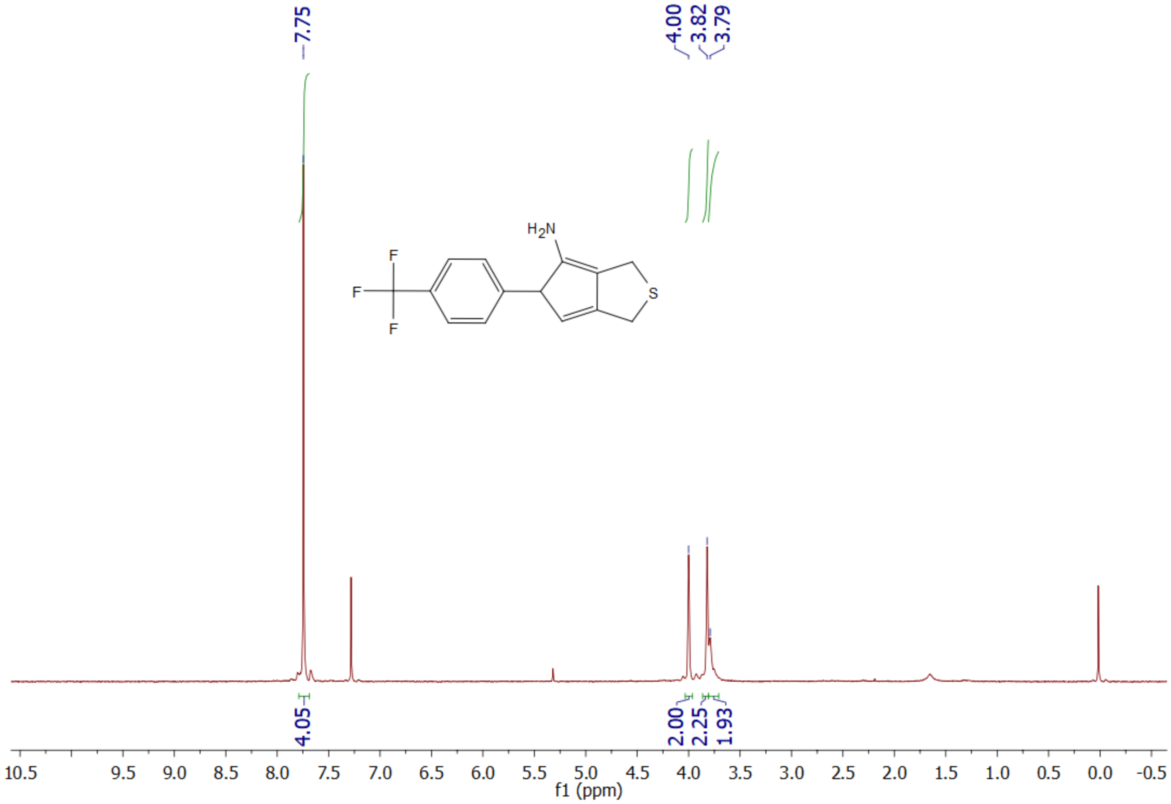


**Supplemental Figure S6**: $^{1}$H NMR Spectrum of **4f**

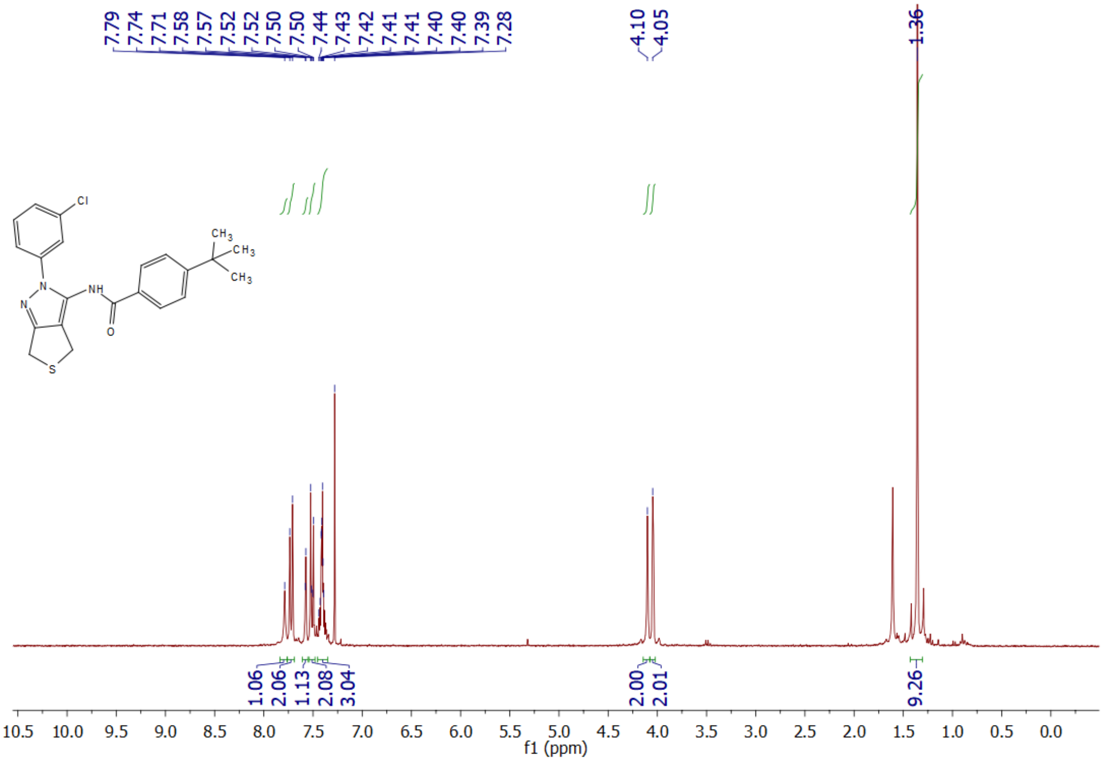

**Supplemental Figure S7**: $^{1}$H NMR Spectrum of **5a**

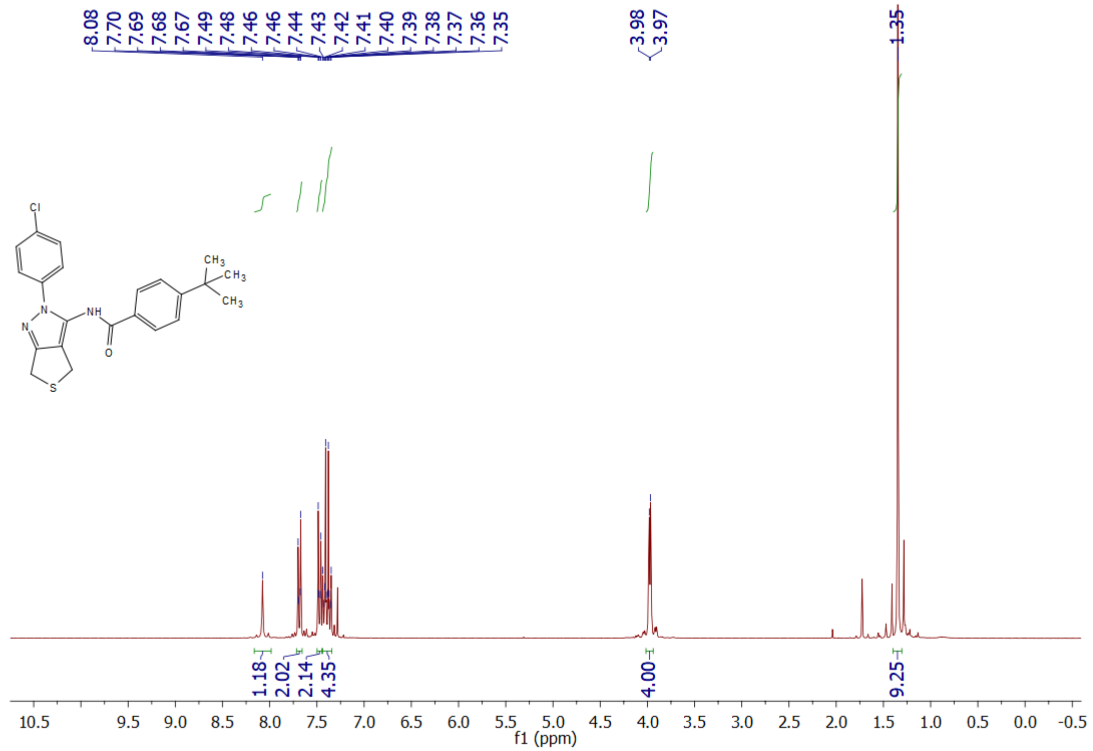

**Supplemental Figure S8**: $^{1}$H NMR Spectrum of **5b**

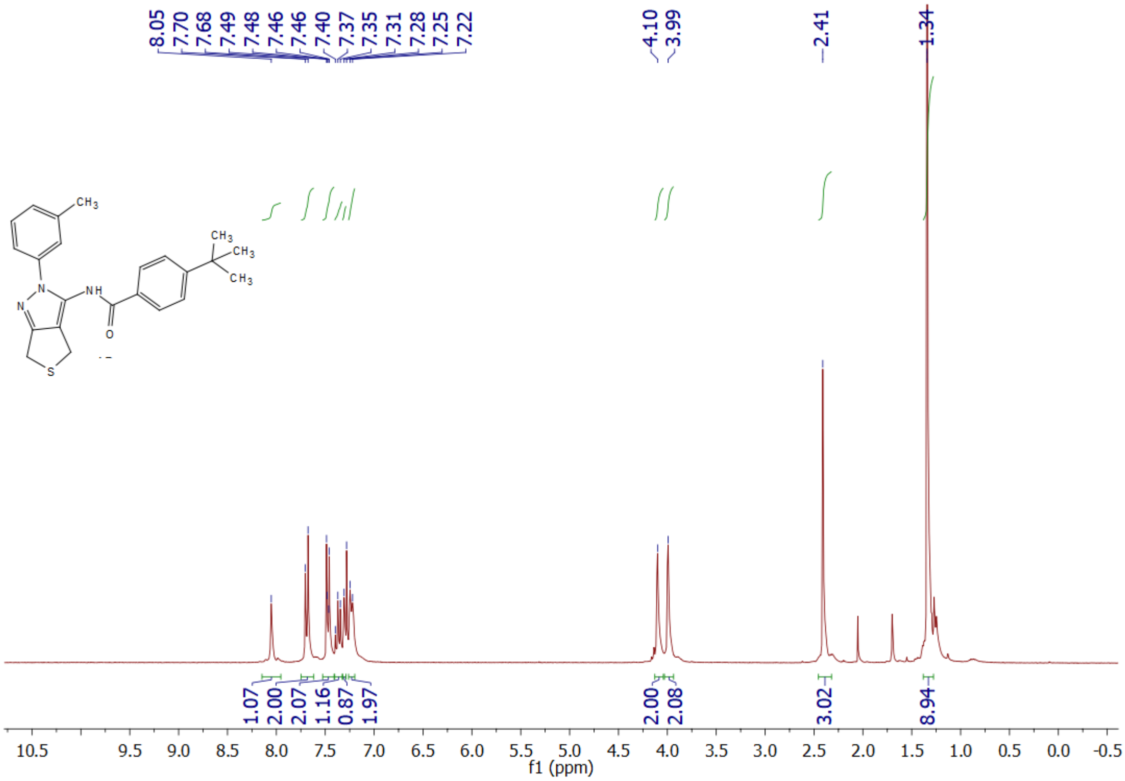


**Supplemental Figure S9**: $^{1}$H NMR Spectrum of **5c**

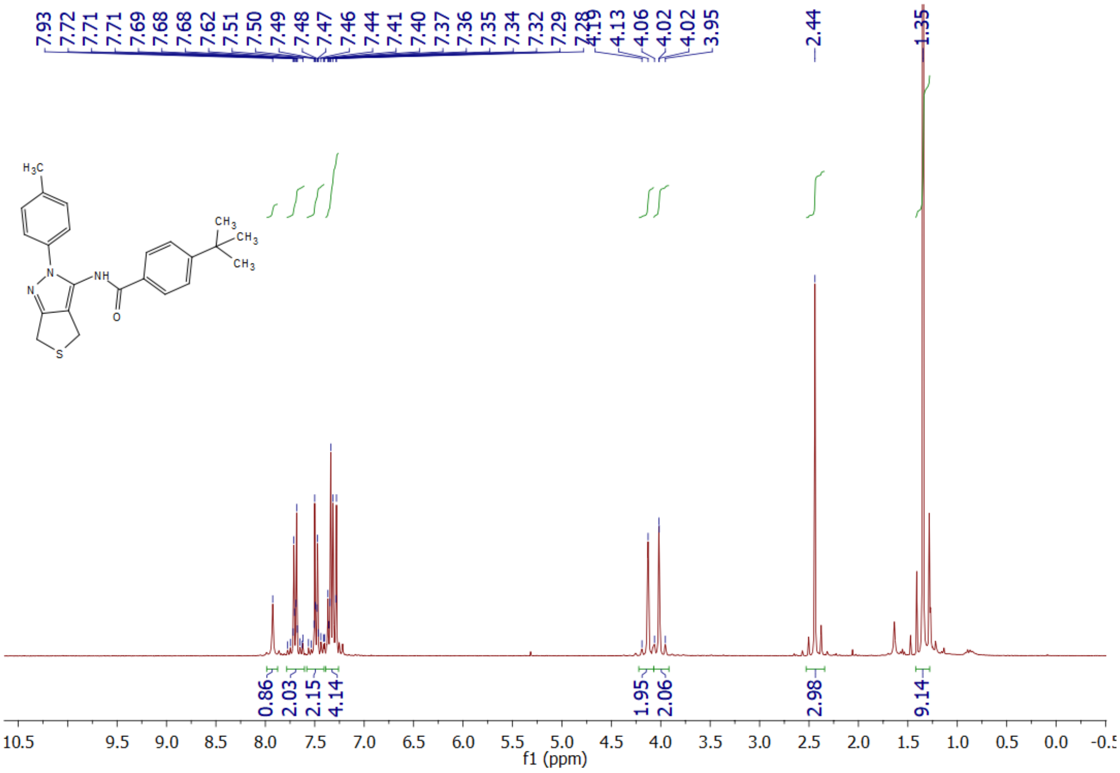


**Supplemental Figure S10**: $^{1}$H NMR Spectrum of **5d**

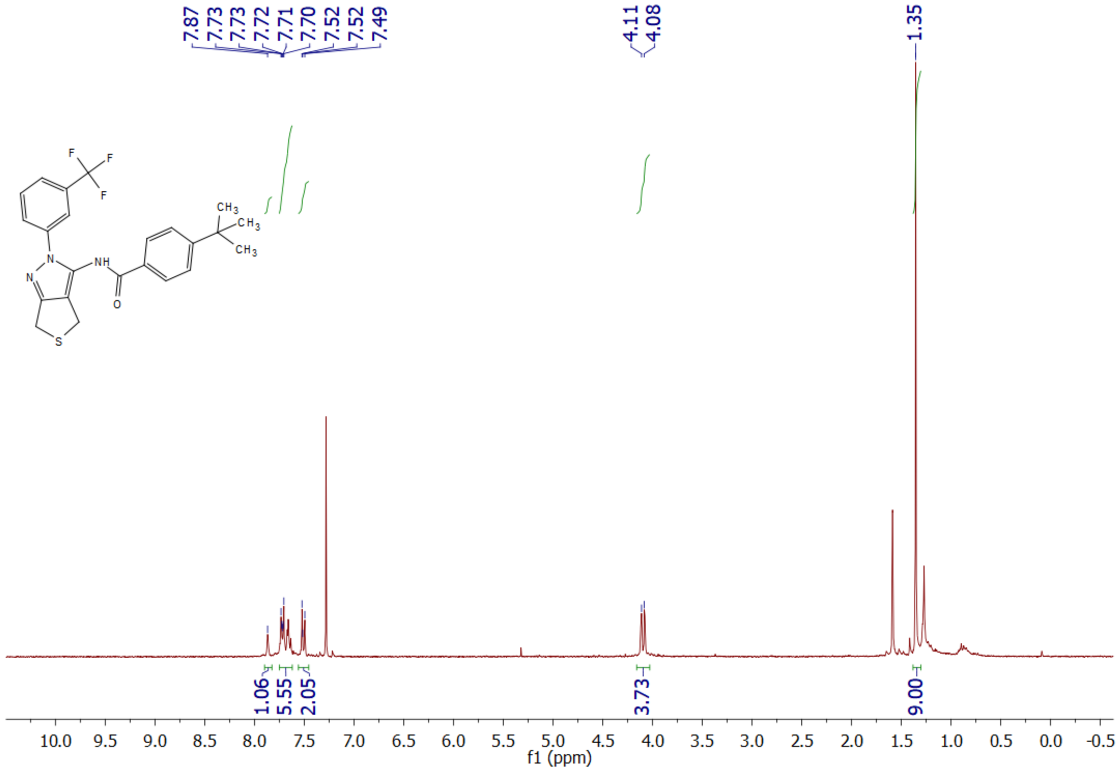


**Supplemental Figure S11**: $^{1}$H NMR Spectrum of **5e**

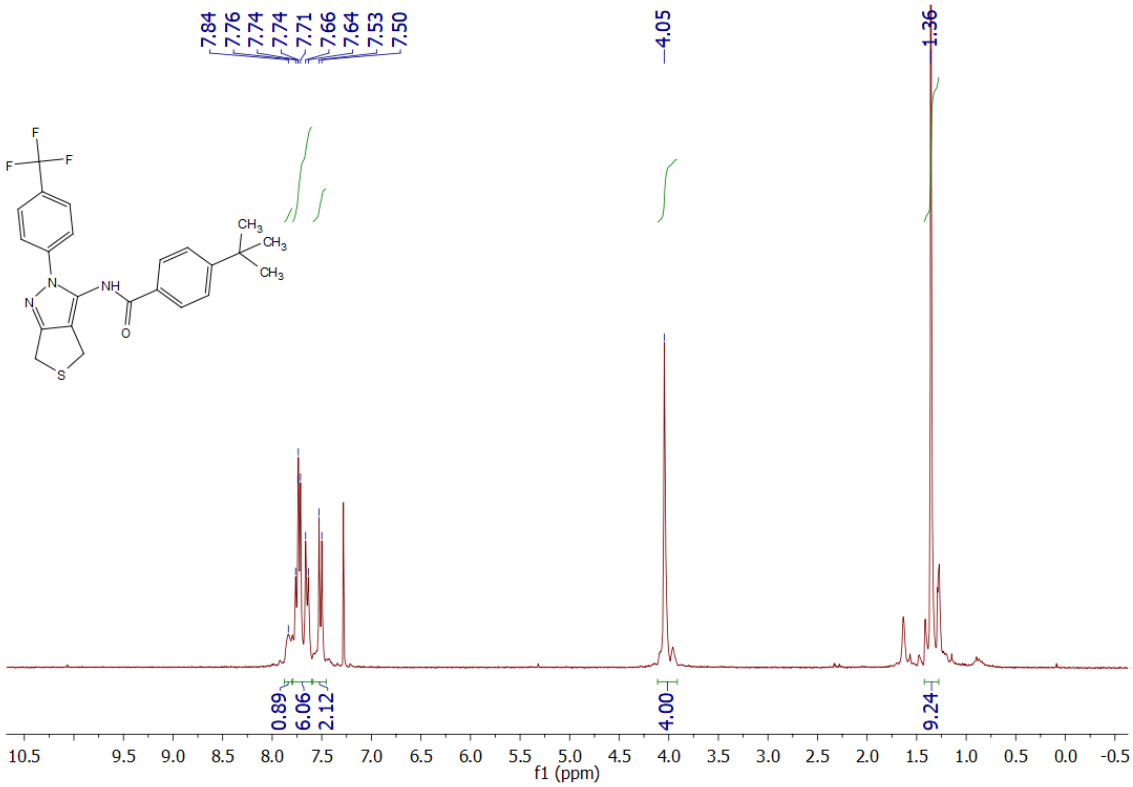


**Supplemental Figure S12**: $^{1}$H NMR Spectrum of **5f**

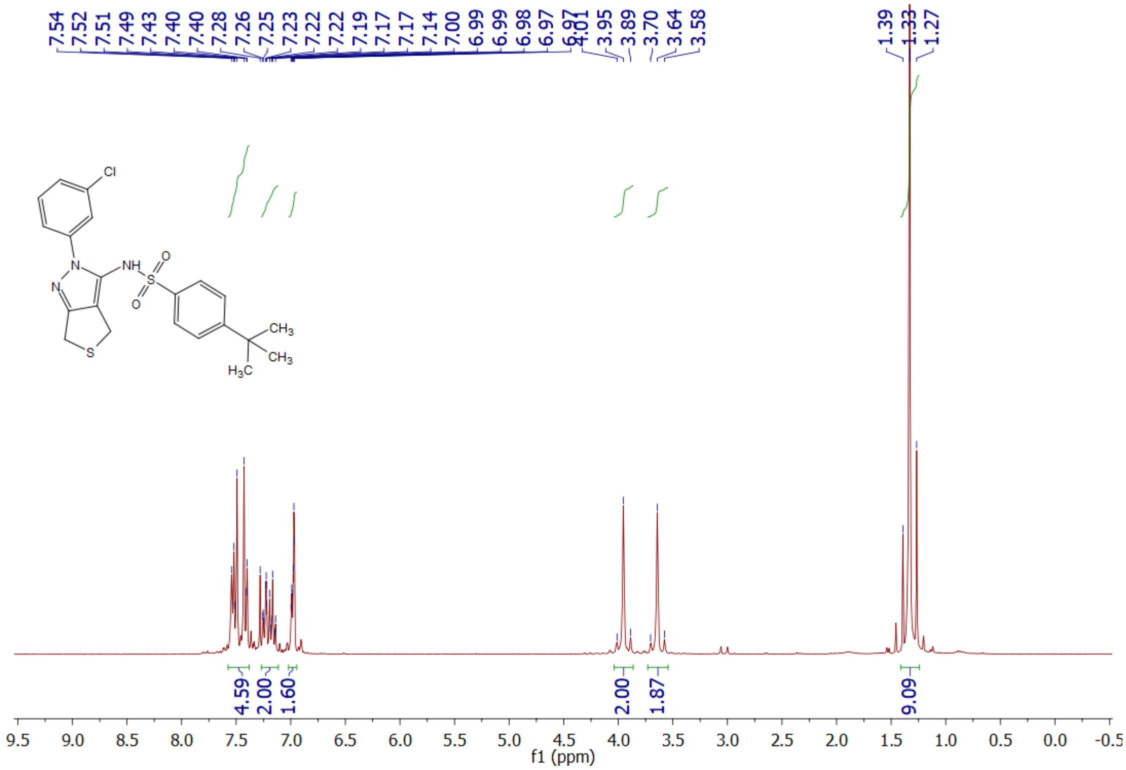


**Supplemental Figure S13**: [1]H NMR Spectrum of **6a**

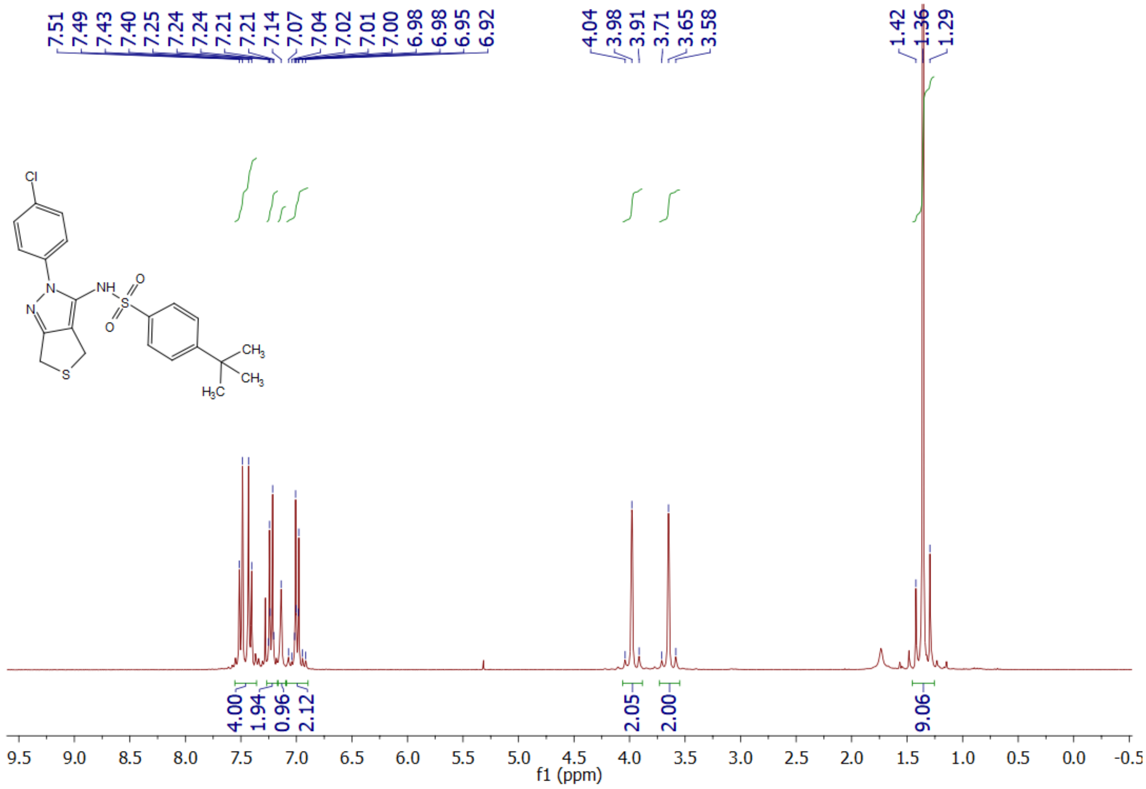


**Supplemental Figure S14**: [1]H NMR Spectrum of **6b**

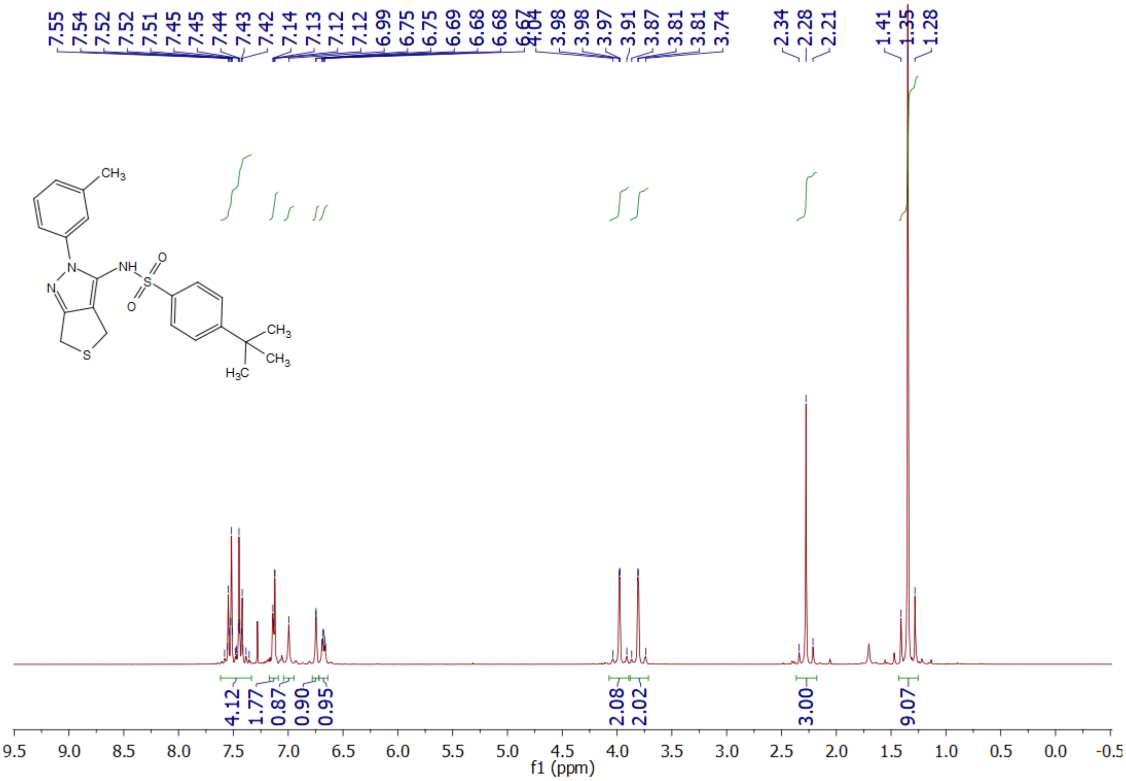


**Supplemental Figure S15**: $^{1}$H NMR Spectrum of **6c**

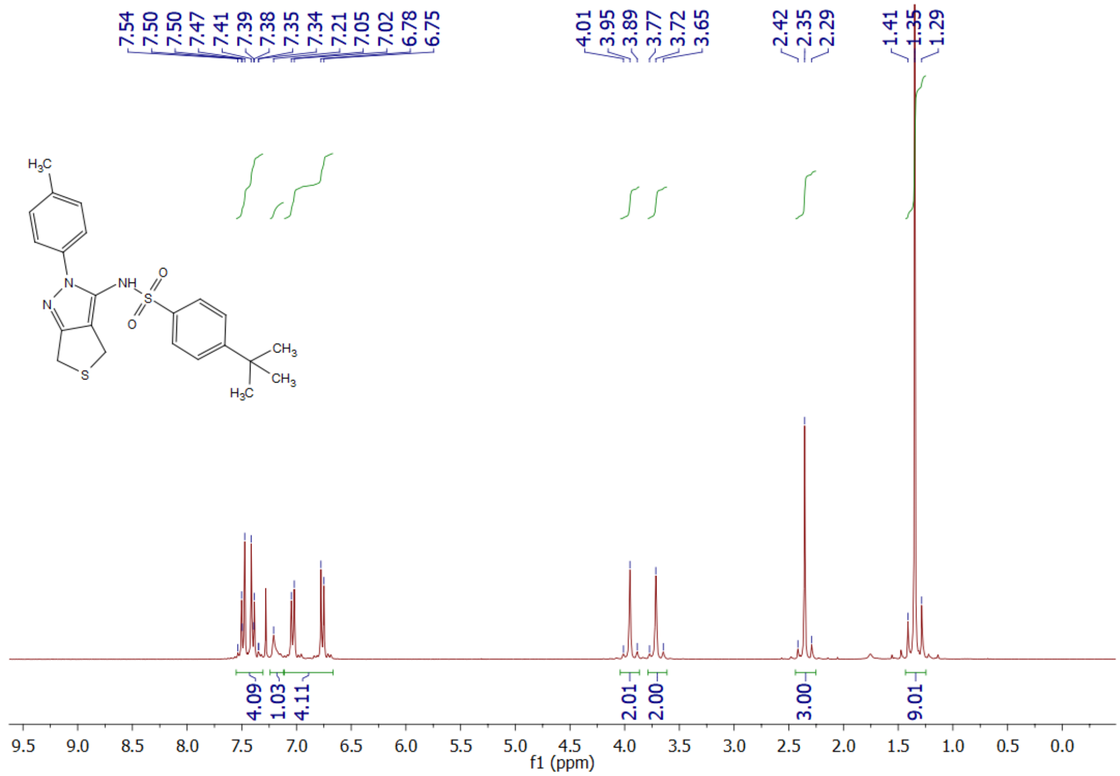


**Supplemental Figure S16**: $^{1}$H NMR Spectrum of **6d**

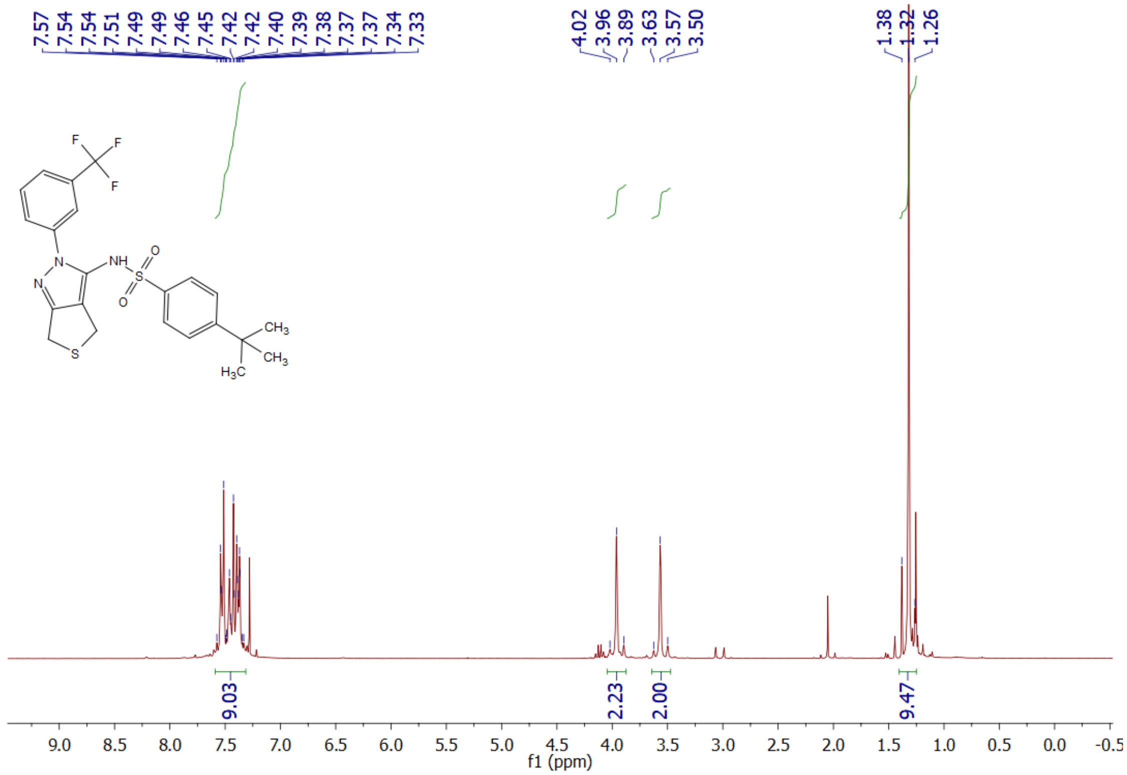


**Supplemental Figure S17**: $^{1}$H NMR Spectrum of **6e**

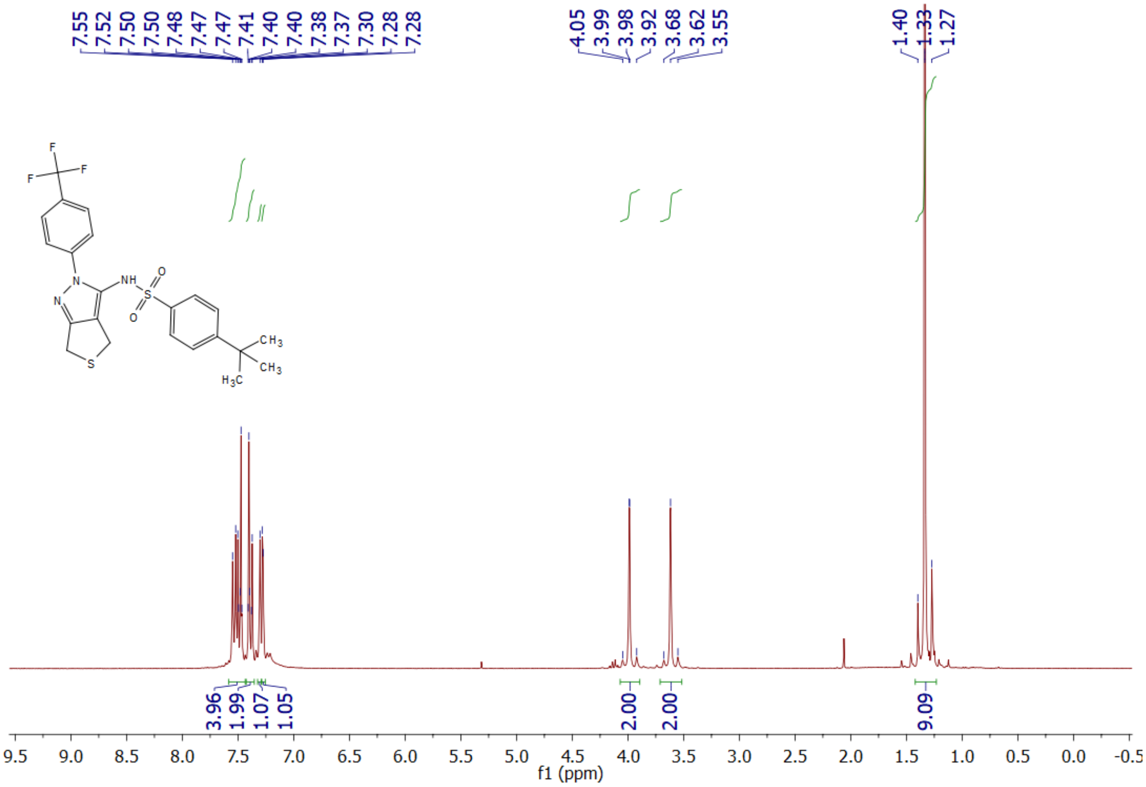


**Supplemental Figure S18**: $^{1}$H NMR Spectrum of **6f**

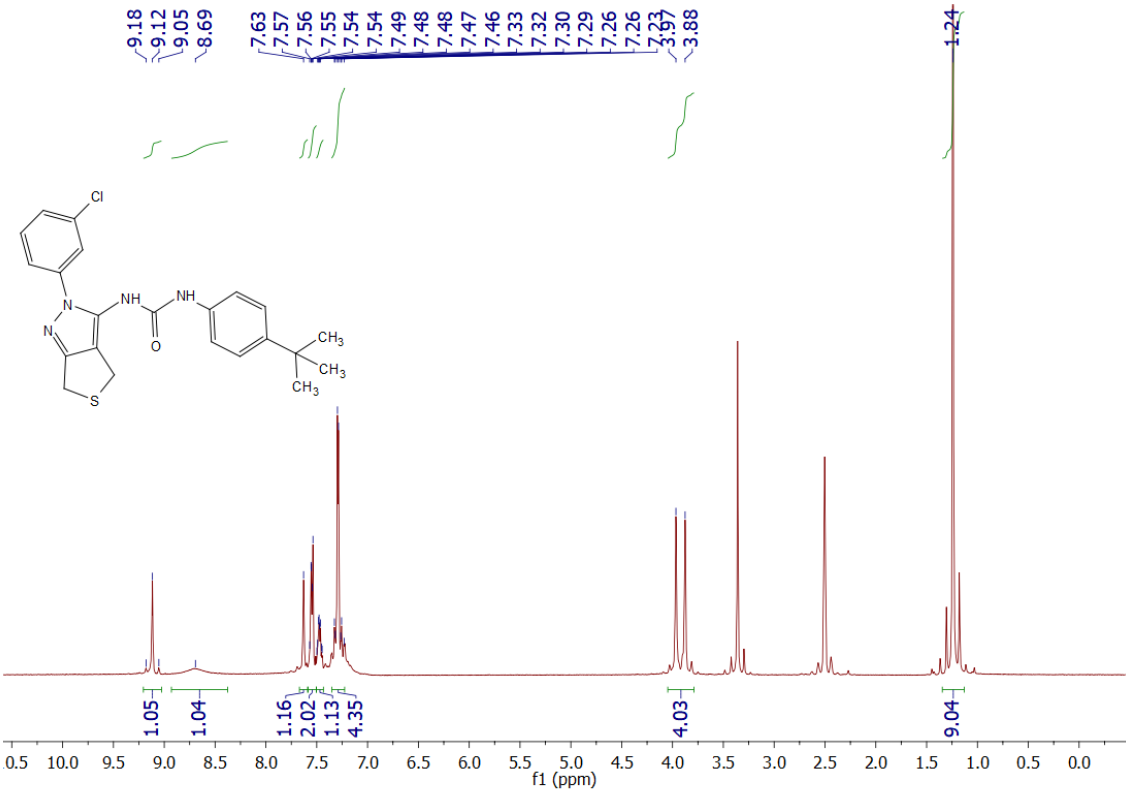

**Supplemental Figure S19**: $^{1}$H NMR Spectrum of **7a**

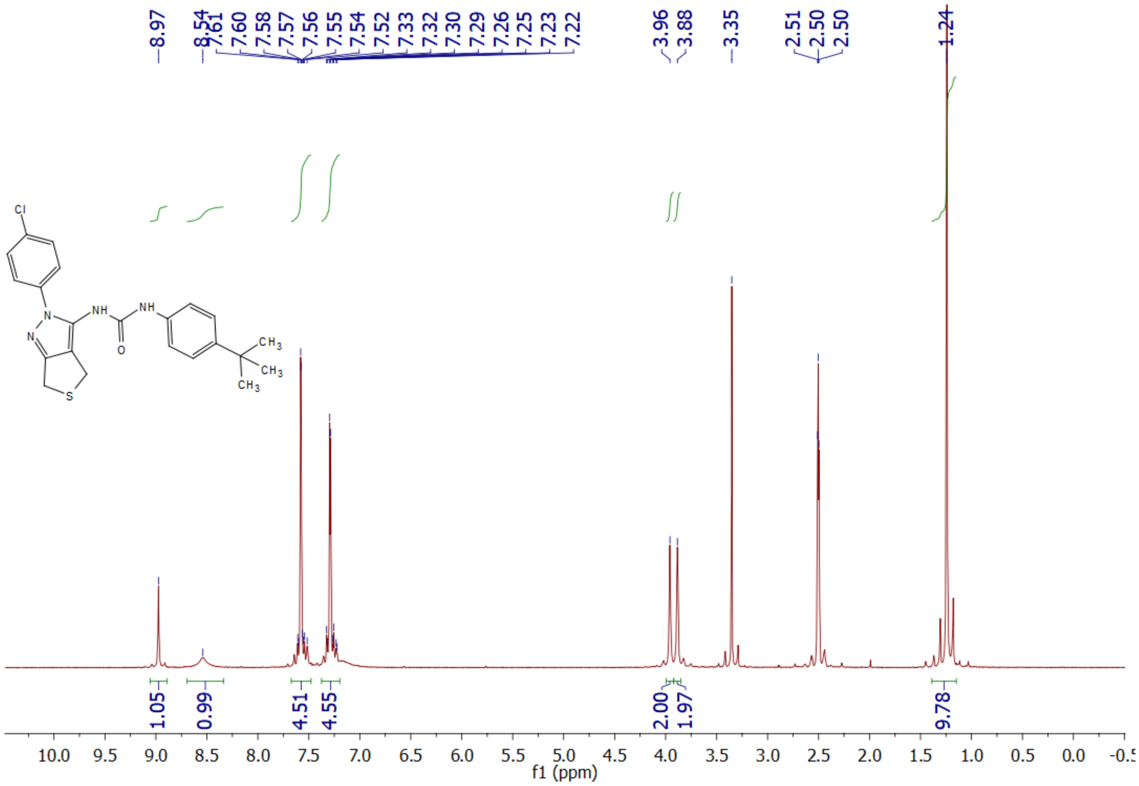

**Supplemental Figure S20**: $^{1}$H NMR Spectrum of **7b**

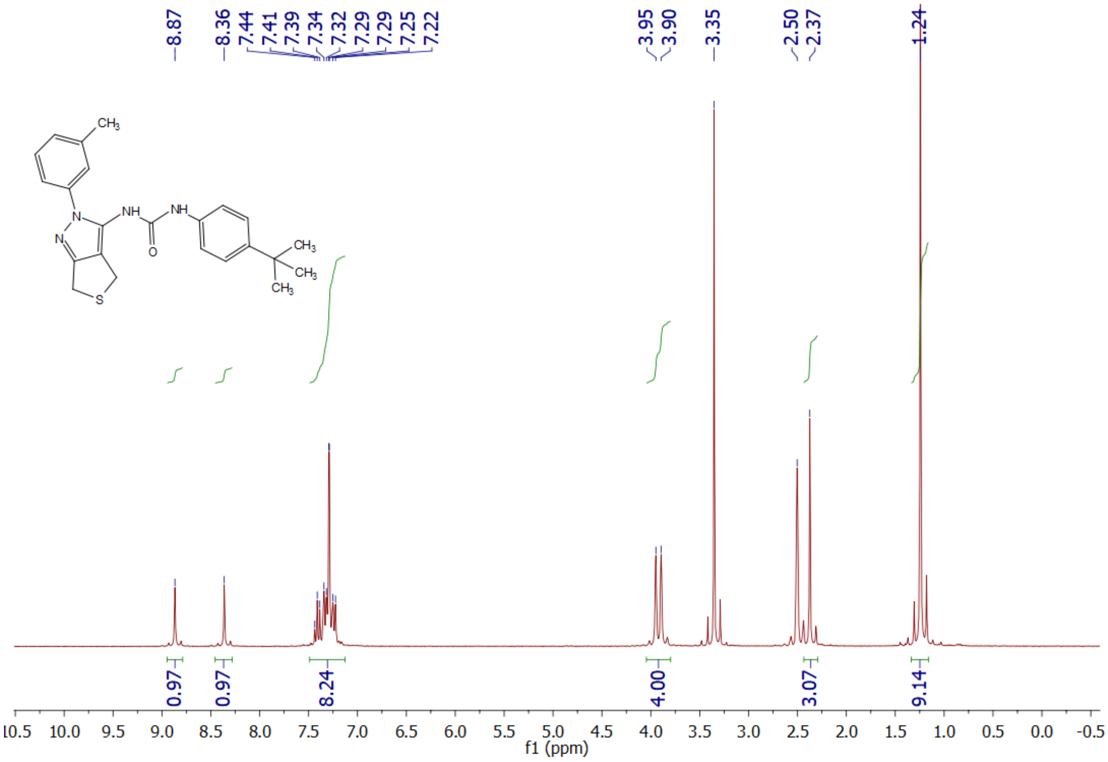


**Supplemental Figure S21**: $^{1}$H NMR Spectrum of **7c**

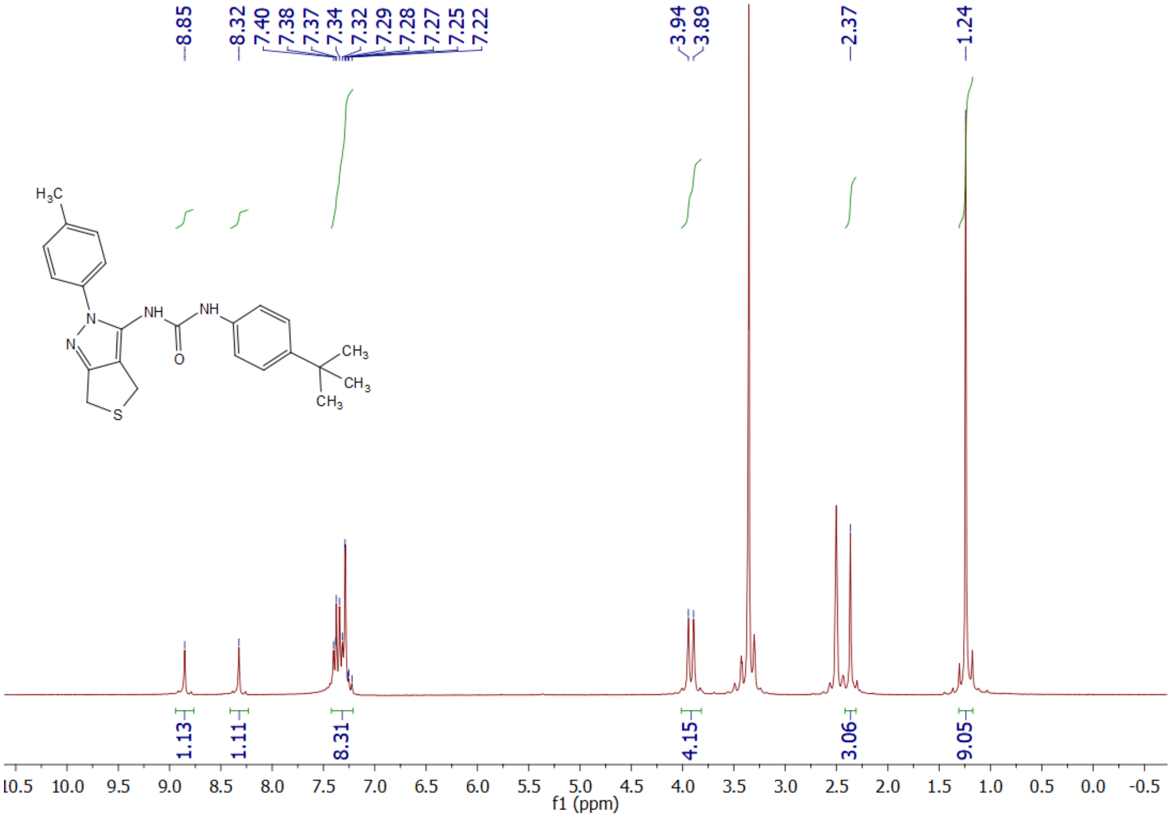


**Supplemental Figure S22**: $^{1}$H NMR Spectrum of **7d**

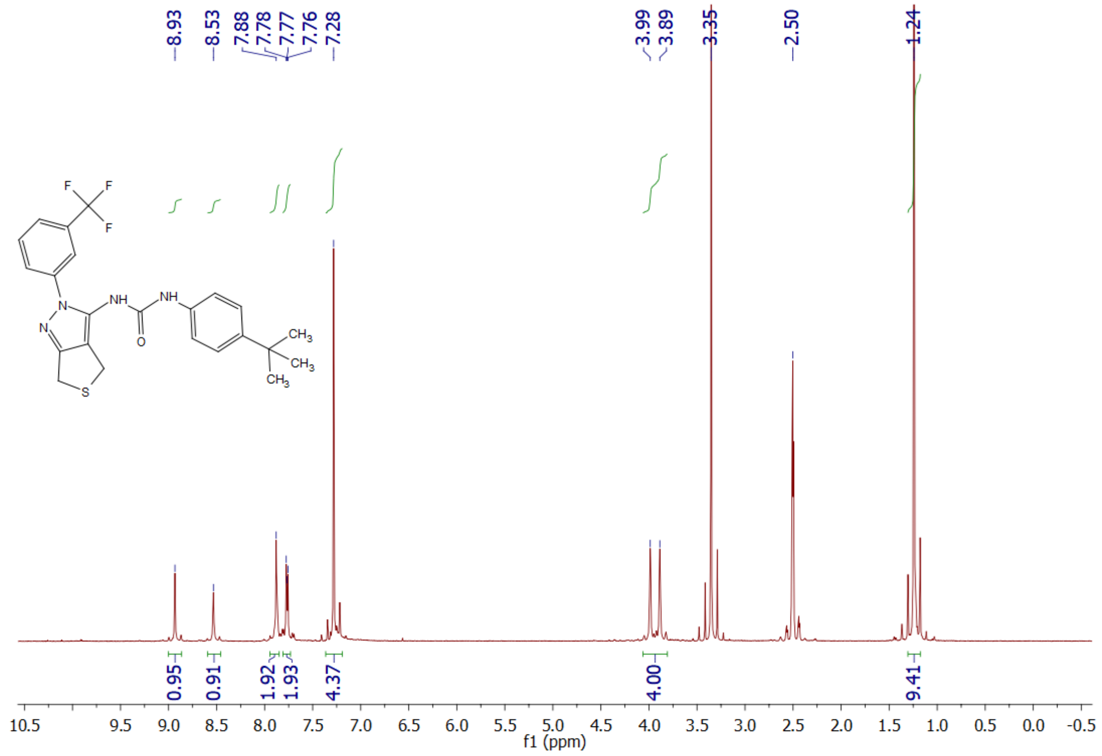


**Supplemental Figure S23**: $^{1}$H NMR Spectrum of **7e**

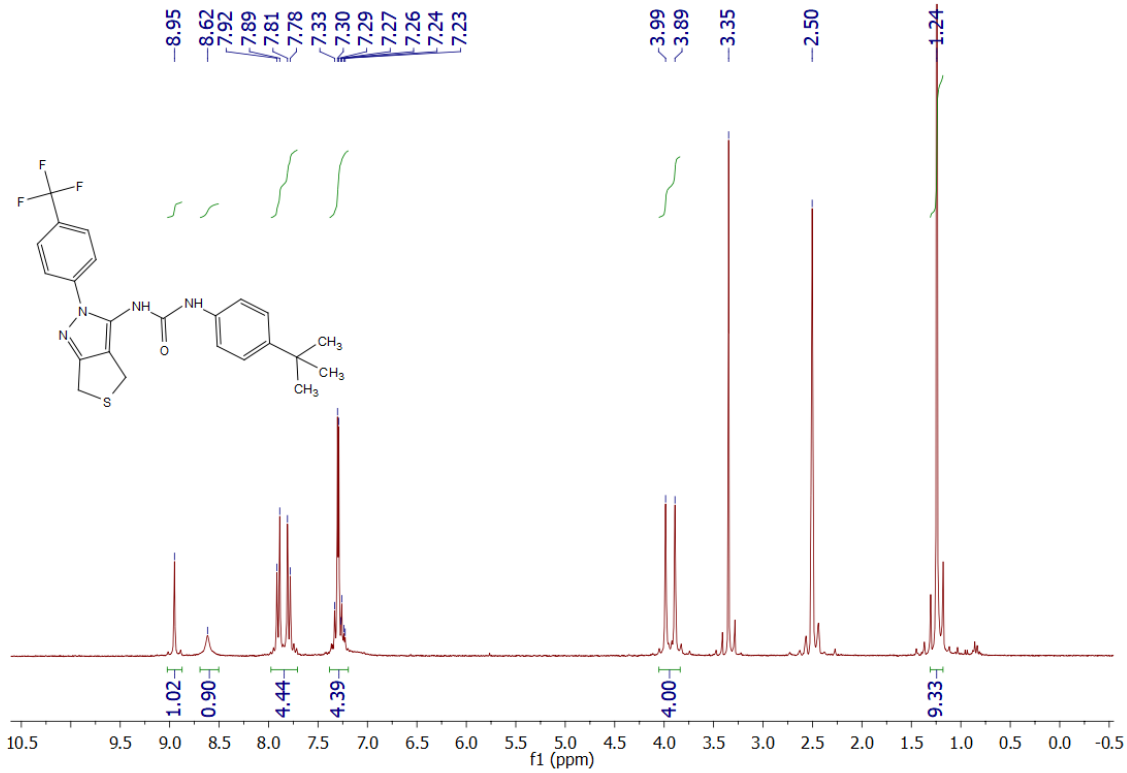


**Supplemental Figure S24**: $^{1}$H NMR Spectrum of **7e**

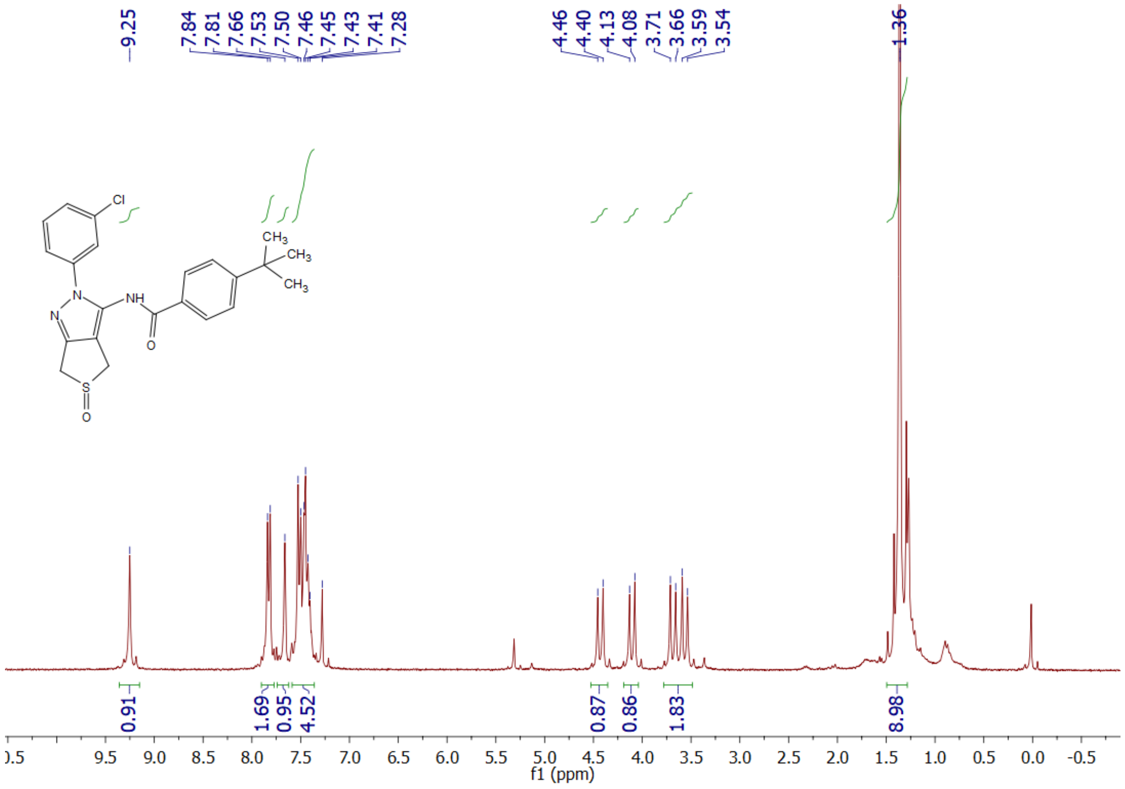


**Supplemental Figure S25**: $^{1}$H NMR Spectrum of **8**

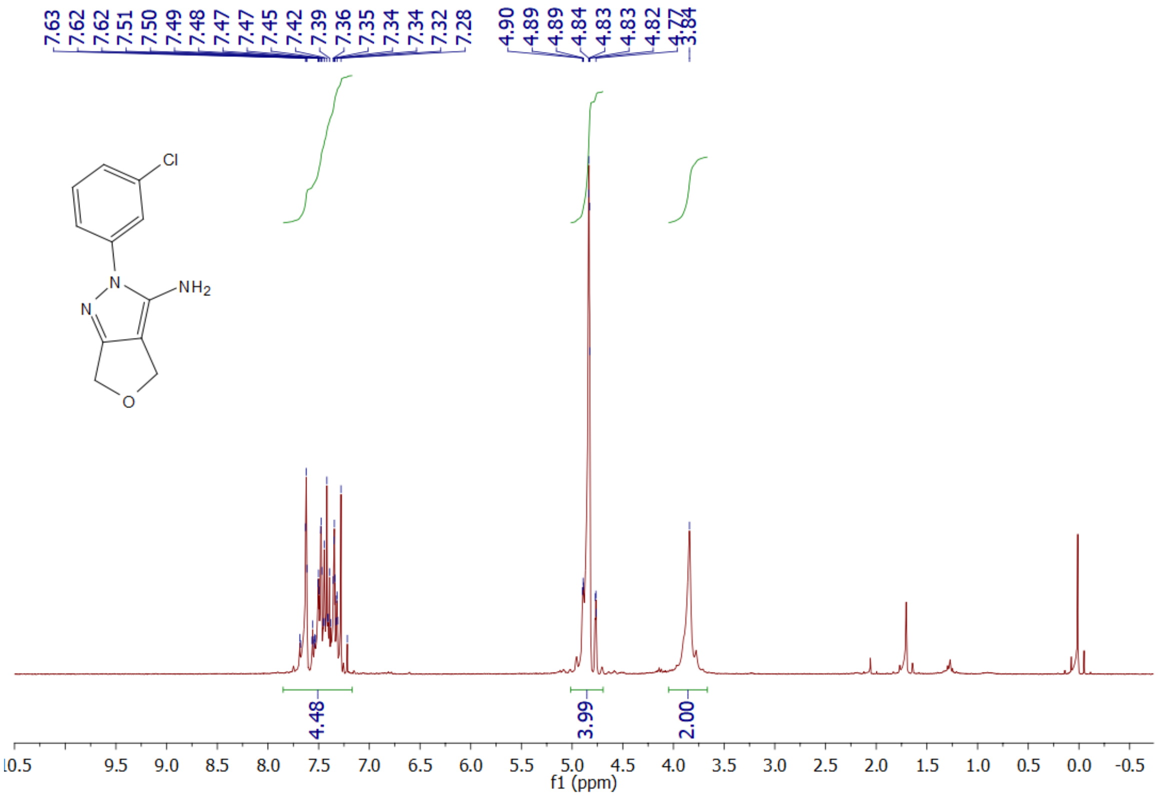


**Supplemental Figure S26**: $^{1}$H NMR Spectrum of **12a**

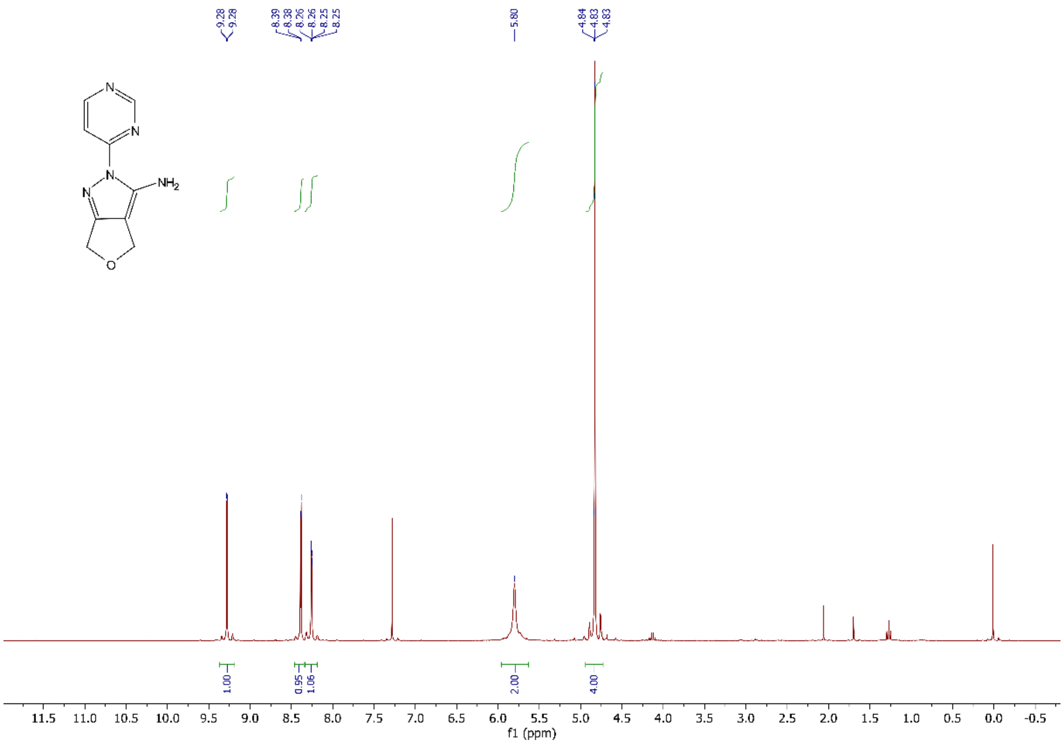


**Supplemental Figure S27**: $^{1}$H NMR Spectrum of **12b**

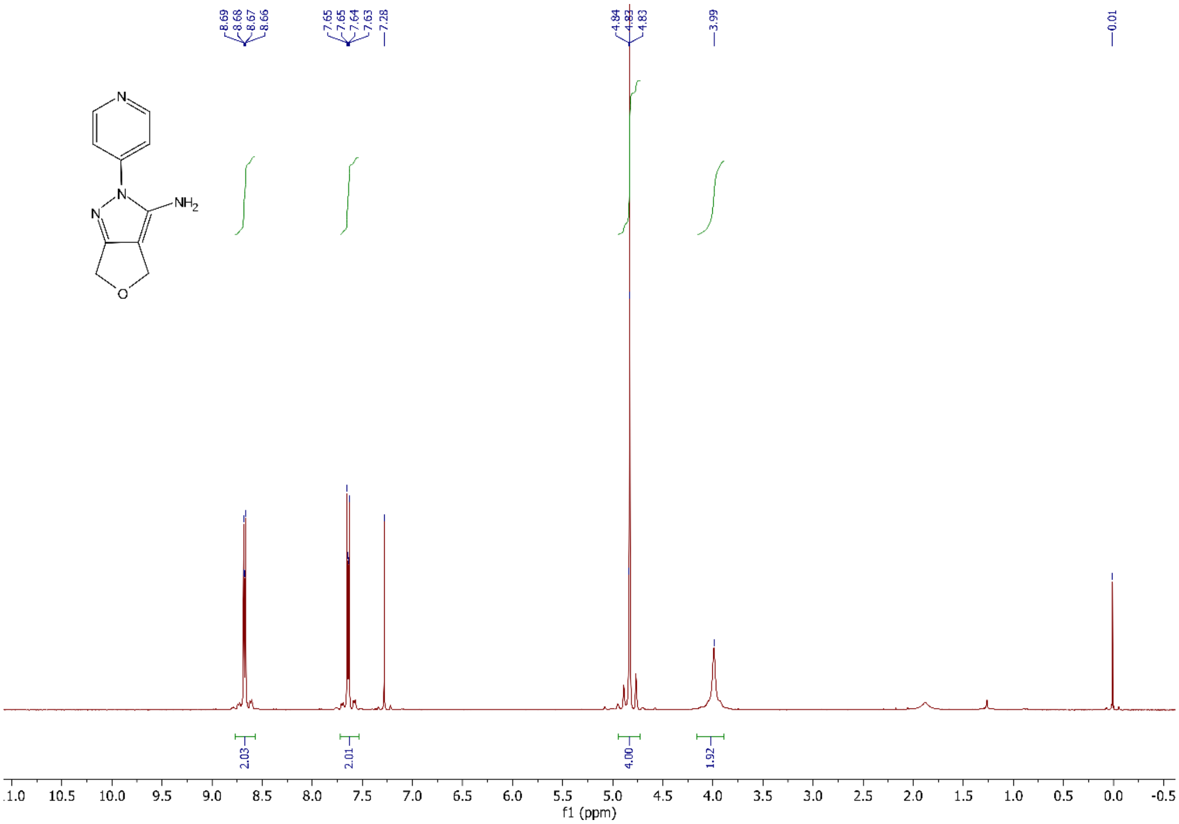


**Supplemental Figure S28**: $^{1}$H NMR Spectrum of **12c**

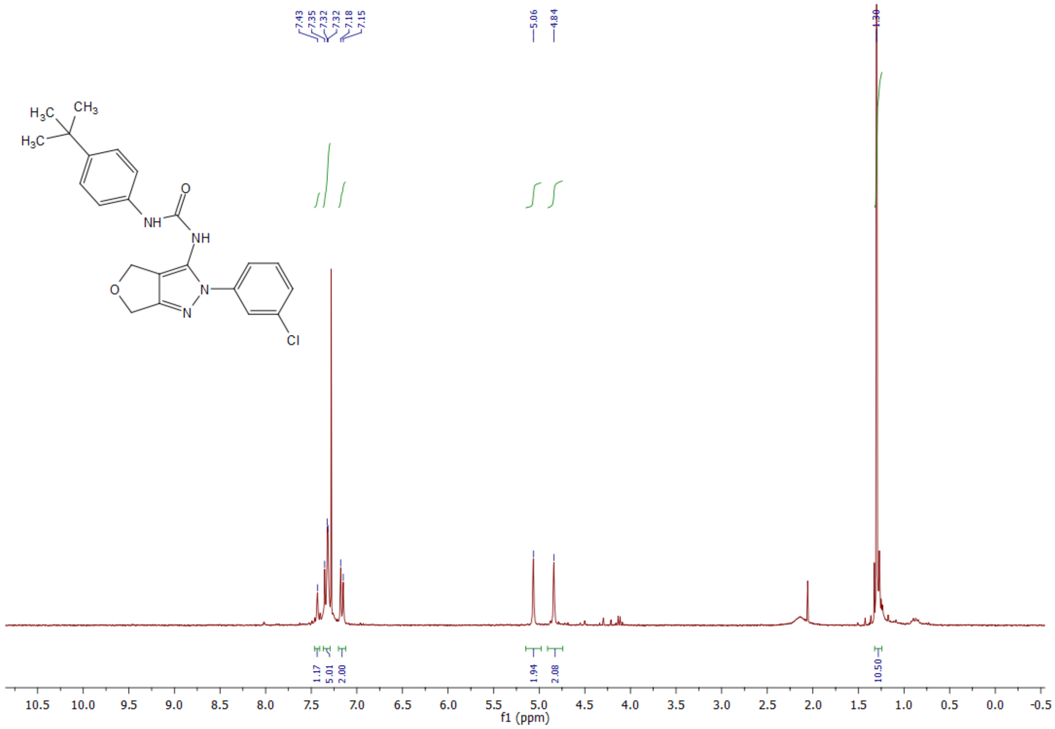

**Supplemental Figure S29**: $^{1}$H NMR Spectrum of **13a**

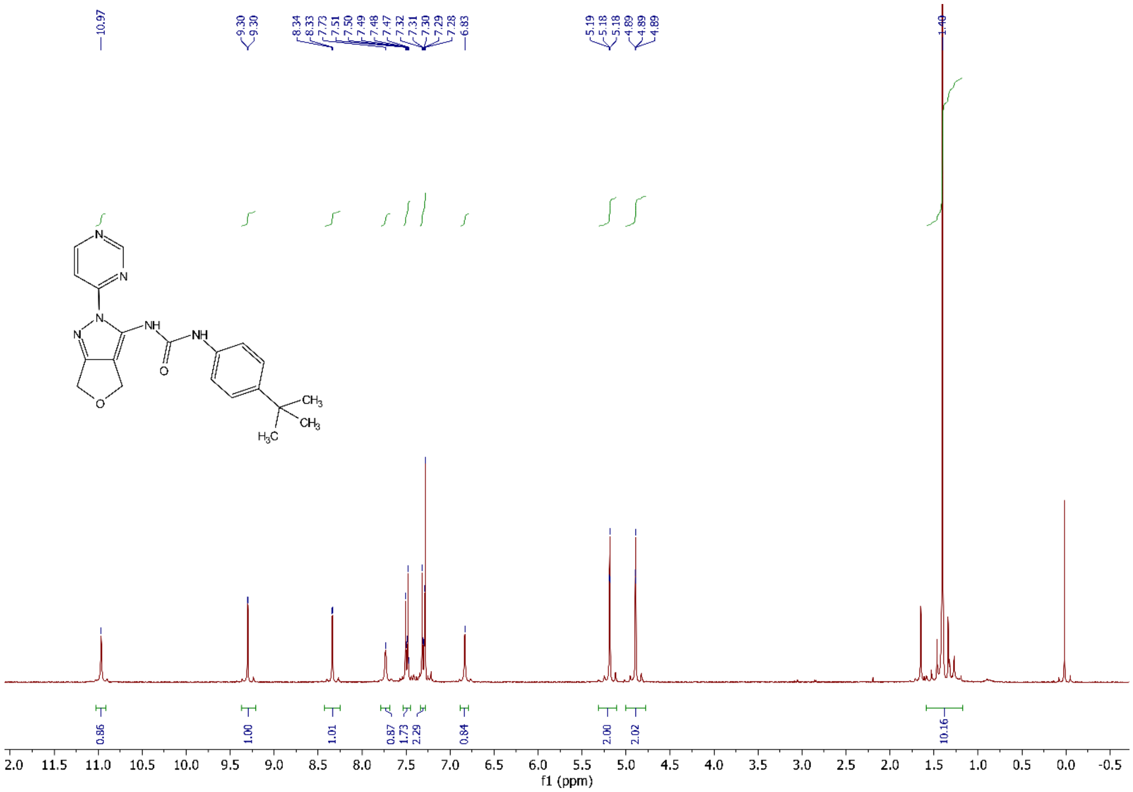

**Supplemental Figure S30**: $^{1}$H NMR Spectrum of **13b**

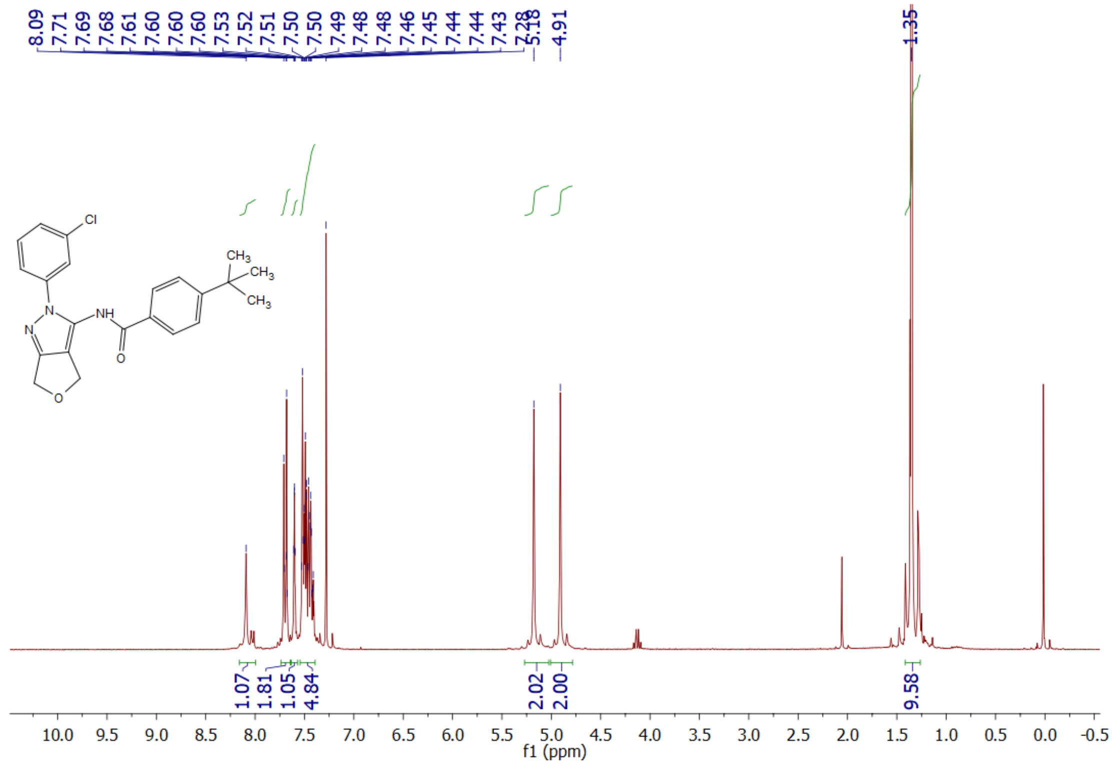

**Supplemental Figure S31**: $^{1}$H NMR Spectrum of **14a**

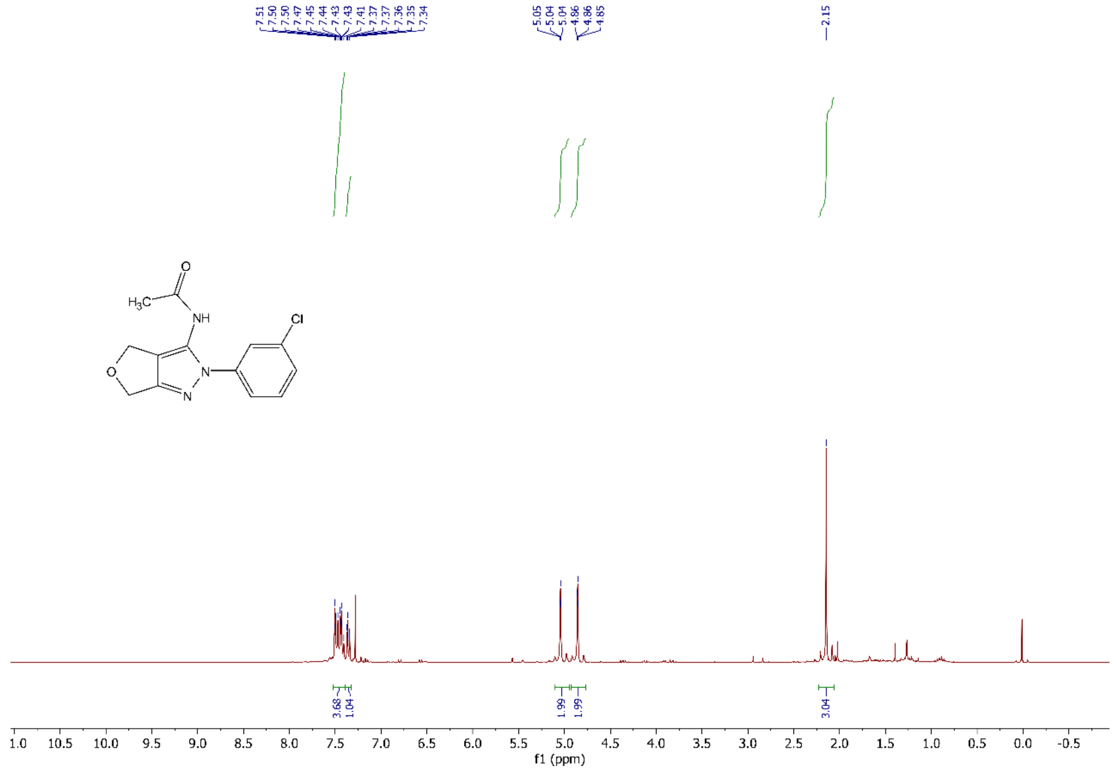

**Supplemental Figure S32**: $^{1}$H NMR Spectrum of **14b**

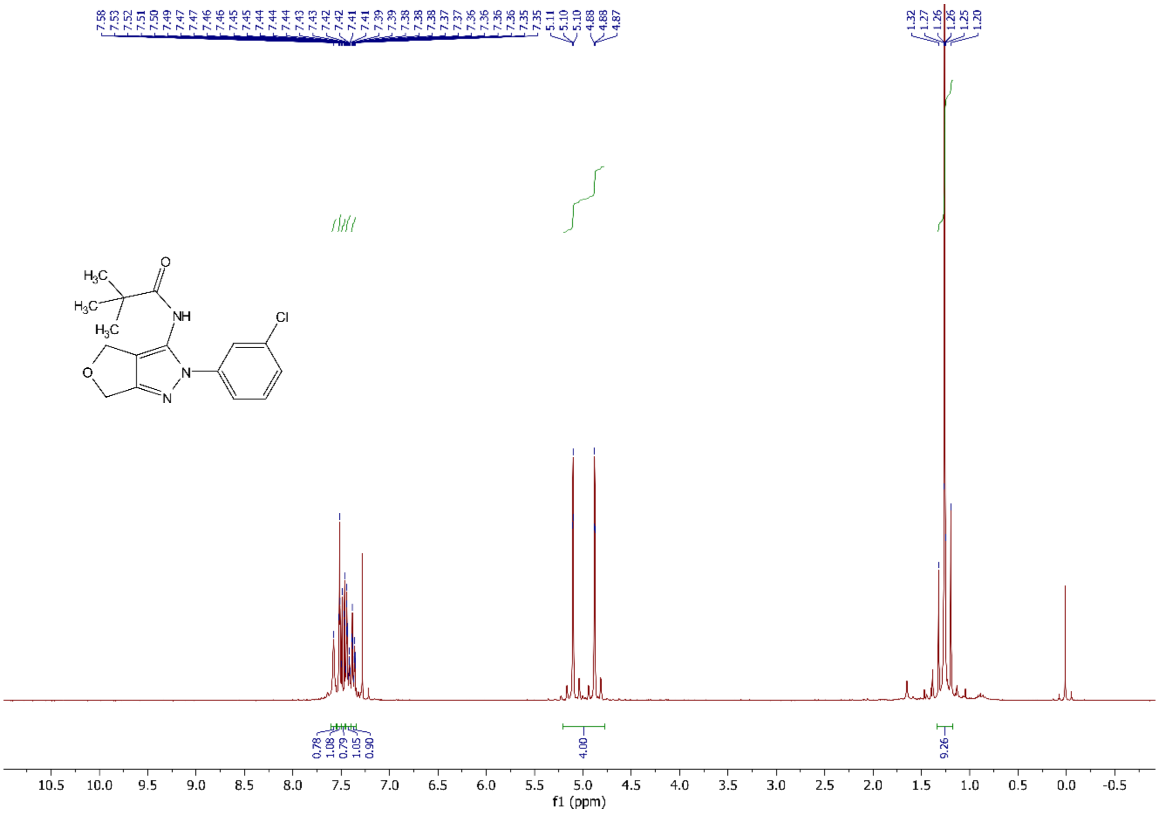

**Supplemental Figure S33**: $^{1}$H NMR Spectrum of **14c**

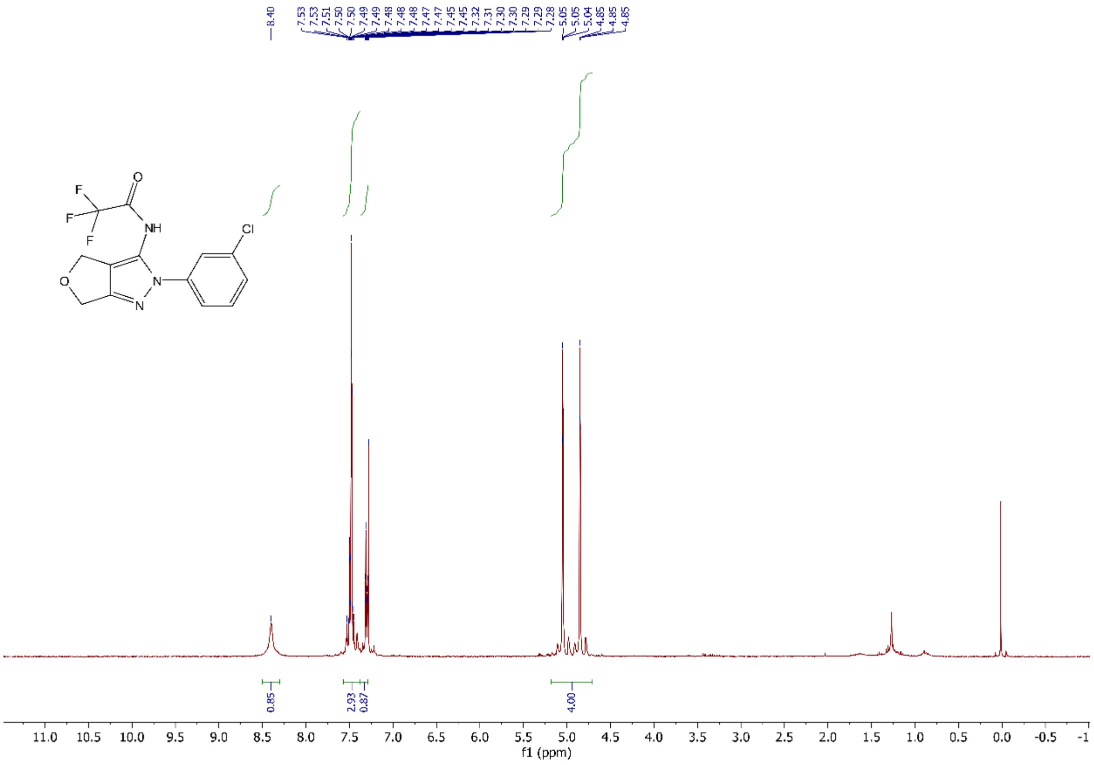

**Supplemental Figure S34**: $^{1}$H NMR Spectrum of **14d**

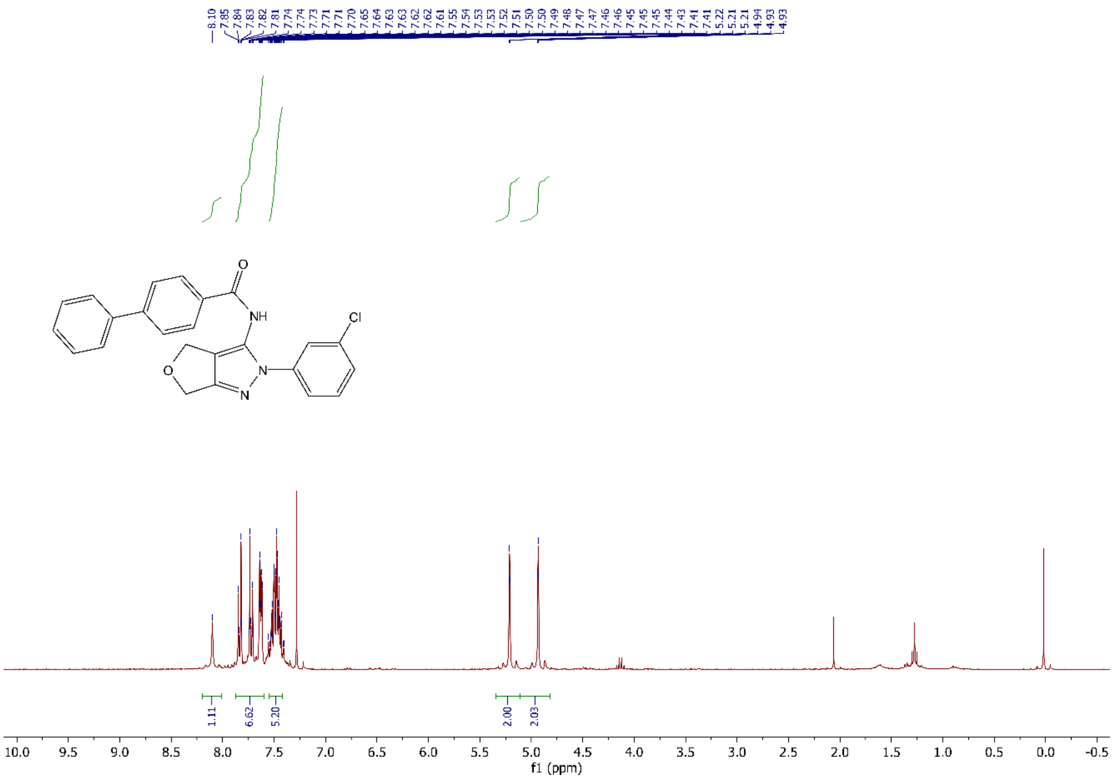


**Supplemental Figure S35**: $^1$H NMR Spectrum of **14e**

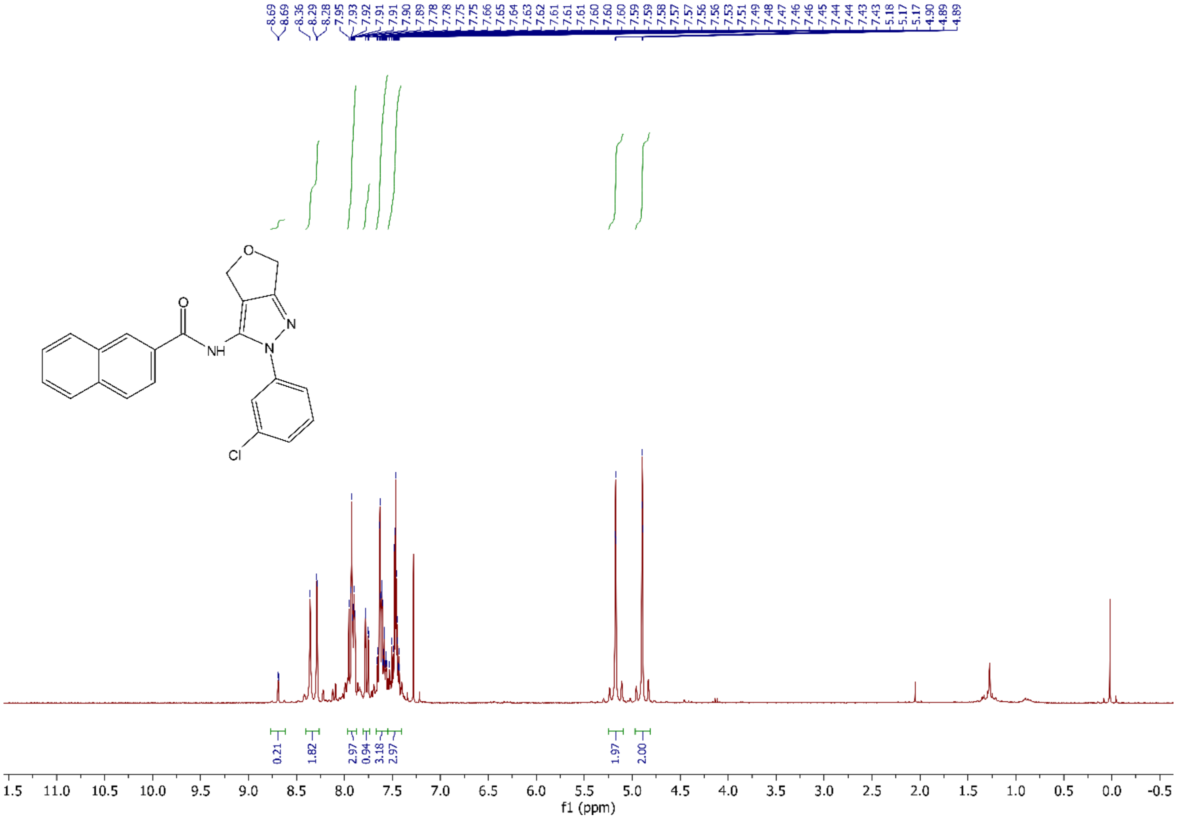


**Supplemental Figure S36**: $^1$H NMR Spectrum of **14f**

**References:**


1. Miller, M. E.; Parrott, E. E.; Singh, R.; Nelson, S. W., A High-Throughput Assay to Identify Inhibitors of the Apicoplast DNA Polymerase from Plasmodium falciparum. *Journal of biomolecular screening* **2014,** *19* (6), 966-72.
2. Kuzmic, P., DynaFit--a software package for enzymology. *Methods in enzymology* **2009**, 467, 247-280. PMID: 19897096.
3. Information, N.C.F.B. Source=ChEMBL, AID=781324 https://pubchem.ncbi.nlm.nih.gov/bioassay/781324 (accessed May 22,2020).
4. Wingert, B.M.; Parrott, E.E.; Nelson, S.W., Fidelity, mismatch, extension, and proofreading activity of the Plasmodium falciparum apicoplast DNA polymerase. *Biochemistry* **2013**, 52 (44), 7723-30.
5. Wang, W.; Malcolm, B.A., Two-stage PCR protocol allowing introduction of multiple mutations, deletions and insertions using QuickChange Site-Directed Mutagenesis. *BioTechniques* **1999**, 26 (4), 680-2.
